# From Neuropeptides to Toxins: Illuminating the Origins of Venom Complexity in Cone Snails

**DOI:** 10.1101/2025.09.26.678268

**Authors:** Thomas Lund Koch, Giada Ferrari, Sunita B Sumanam, Paula Flórez Salcedo, Maren Watkins, Kevin Chase, Ave Tooming-Klunderud, Arturo O Lluisma, Angel Yanagihara, Neil D Young, Baldomero M Olivera, Eivind A.B. Undheim, Helena Safavi-Hemami

## Abstract

New genes and gene functions are key drivers of evolutionary innovation. Venomous animals, such as cone snails, provide striking examples of gene innovation, yet the mechanisms by which toxins arise remain poorly understood. Using the *Conus textile* genome, we uncover how neuropeptide genes were recruited into the venom and neofunctionalized as doppelgänger toxins. We identify over 20 independent recruitment events that evolved dynamically across the *Conus* lineage. Rather than arising from ohnologs of a whole-genome duplication event ∼100 mya, these toxins evolved through diverse mechanisms, including exon shuffling, alternative splicing, and ectopic recombination, often facilitated by lineage-specific transposable elements. Our findings reveal a dynamic interplay between genome architecture and molecular innovation, offering broad insight into the evolution of complex gene repertoires in venoms and beyond.

**One-Sentence Summary:** Doppelgänger toxins reveal how modular gene architecture, including 5’UTR reuse and TE-driven recombination, fuels gene innovation in cone snails.

## Main Text

New genes and gene functions are fundamental drivers of evolutionary innovations that enable organisms to acquire novel traits and adapt to changing environments^1–3^. A central question in biology is how such genes and functions arise. Neofunctionalization, in which genes acquire new roles, arises from changes in either coding or regulatory regions and may occur independently of gene duplication. While new genes can arise through diverse mechanisms, the evolutionary processes linking genetic changes to novel functions remain poorly understood.

Venom systems are compelling models to investigate the relationship between new genes, new functions, and evolutionary innovation. Toxins evolve rapidly^4^, often belong to large gene families^5,6^, and have important biological functions^7^. Further, many toxins originate from non-toxin genes and undergo changes in gene expression, shifting from non-venomous tissues to specialized toxin-secreting glands in a process referred to as “recruitment”^8^. However, the short length and rapid evolution of most toxins can obscure their evolutionary origins. A notable exception is doppelgänger toxins – molecular mimics of neuropeptides such as insulin-like and oxytocin-like toxins^9,10^ – which typically retain substantial similarity to their ancestral neuropeptide gene, enabling their origins to be traced.

Cone snails, a hyper-diverse lineage of marine gastropods, offer an ideal model to investigate genetic innovations, as they have evolved some of the most complex and rapidly evolving venoms in the animal kingdom^11^. These snails produce a wide array of peptide toxins (conotoxins) to capture prey that includes worms, fish, and mollusks. Notably, although all cone snail lineages are known to use doppelgänger toxins, the more recently evolved mollusk-hunting species^12^ express a significantly higher number of these toxins^13,14^. This enrichment suggests that mollusk-hunting cone snails represent an especially powerful model for investigating the evolution of new genes, particularly given that the genomic origins of doppelgänger toxins remain largely unknown.

Recently, a whole genome duplication (WGD) in ancestral neogastropods was identified^15^, raising the possibility that this duplication event provided genetic material for evolutionary innovations. A compelling hypothesis is that duplicated neuropeptide genes arising from this event were co-opted into the venom system, giving rise to doppelgänger toxins. In other lineages, WGD has been a major driver of gene diversification and functional innovation^16,17^. For example, in flowering plants, WGD has contributed to the expansion of key developmental gene families^18^, and in vertebrates, it facilitated the evolution of distinct oxygen-binding proteins - hemoglobulin and myoglobulin from a common globulin gene^19^.

To investigate the genomic mechanisms underlying evolutionary innovation in cone snails, particularly the role of WGD in recruiting neuropeptides into the venom system, we conducted large-scale transcriptome analysis and whole genome sequencing of the mollusk-hunting “cloth-of-gold” cone, *Conus textile*. While we confirm the occurrence of a WGD, our results show that this event did not directly facilitate the evolution of doppelgänger toxins. Instead, we identify multiple genomic processes that facilitate recruitment, including modular recombination via exon shuffling and alternative splicing, often driven by transposable elements (TEs). We find that most doppelgänger toxins have recombined with the 5’ untranslated regions (5’UTRs) of existing conotoxin genes, likely enabling their expression in the venom gland and escape from pseudogenization. Our findings shed new light on how novel genes and functions arise in rapidly evolving systems and provide a broader framework for understanding rapid gene evolution and complexity across eukaryotes.

### Genome assembly of *Conus textile*

*C. textile* inhabits the shallow waters of the Indo-Pacific where it preys on gastropod mollusks (**Fig 1A, Movie S1-S2**). We sequenced the genome of a single male *C. textile* specimen (**Fig. S1**) using PacBio long-read and Hi-C data generated from genomic DNA. Our data assembled into 3.55 Gbp (∼22x PacBio HiFi read coverage) comprised of 35 long scaffolds (179 - 58 Mbp, N50 = 91.8 Mbp) (**Fig. S2, Table S1**) that are homologous to chromosomes described in other *Conus* species^20–22^. Over 59.6% of the genome is composed of TEs (Retrotransposons: 21%, DNA transposons: 17.6%, Unclassified: 11.9%, and simple repeats: 7.9%) (**Table S2**).

**Fig. 1.**
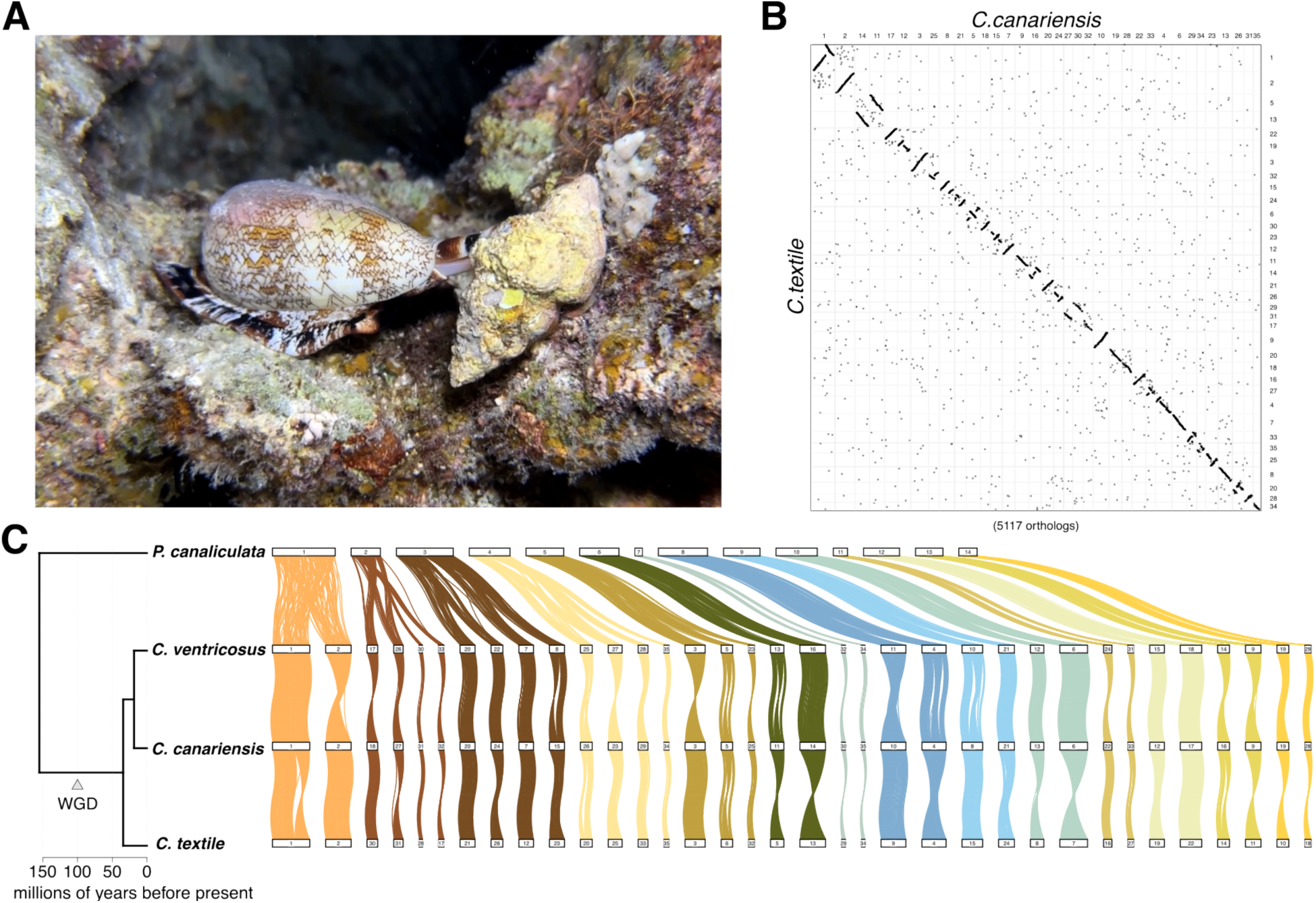
The genome of *C. textile.* **(A)** The “cloth-of-gold” cone, *C. textile,* hunting on a coral reef at night (**Movie S1**). (**B**) Oxford-dotplot showing long colinear regions between *C. textile* and *C. canariensis*. Each dot represents orthologous proteins in the *C. textile* and *C. canariensis* genomes and their location in the respective genomes. (**C**) Conserved linkage groups between *P. canaliculata* and the cone snails *C. ventricosus, C. canariensis,* and *C. textile.* Vertical lines between chromosomes link orthologous genes from conserved linkage groups. The timing of the WDG is indicated in the species tree.

Transcriptomic data from 6 different tissues (circumoesophageal nerve ring, hepatopancreas, venom gland, venom bulb, foot, and salivary gland) from the same individual were used for gene model predictions. Combined with homology-based methods, we predicted 34,172 protein-coding genes, capturing 91% of all metazoan BUSCO (**Table S3**), and 89% OMArk completeness (**Fig. S3**). Of these, 78% have similarity to well-curated proteins in EggNOGdb. Our assembled venom gland transcriptome contained 163 toxin transcripts from 33 superfamilies, with the O2 superfamily being the most abundant in both number and expression (40 transcripts, 211,192 transcripts per million) (**Table S4**). We annotated 199 full-length toxin-coding genes in the genome (127 unique proteins) distributed across 27 chromosomes (**Table S5**, **Fig. S4**).

We find long chromosomal collinearity and microsynteny between *C. textile* and *Conus canariensis*, which diverged approximately 30 million years ago ^23^ (**Fig. 1B, Figs. S5-6**). While several chromosomes show inversions, translocations were relatively rare (n = 982 of 5117 orthologs). A comparison with the golden apple snail, *Pomacea canaliculata,* confirms WGD in the ancestor of cone snails, as the chromosomes in *P. canaliculata* exhibit clear macrosynteny with *Conus* chromosomes, showing 1:2 or 1:4 correspondences (**Fig. 1C, Figs. S7-8**).

### *Conus* doppelgänger toxins did not arise from neuropeptide ohnologs

Having confirmed a WGD in the ancestors of cone snails and its retention in *C. textile,* we next investigated whether neuropeptide ohnologs (paralogs arising from WGD) provided ‘spare copies’ giving rise to doppelgänger toxins or whether they retained their ancestral neuropeptide function. The latter would be revealed by their expression in the circumoesophageal nerve ring, containing multiple neural ganglia.

We identified 61 neuropeptide families in the nerve ring transcriptomes of *C. textile*, *Conus imperialis*, and *Conus rolani,* with most families represented by two homologs (**Fig. 2A, Data S1**). Furthermore, the *C. textile* neuropeptide homologs are consistently located on homologous chromosomes, supporting their origin from WGD and designation as ohnologs (**Fig. S9**). While the overall retention rate of ohnologs across the *C. textile* genome is approximately 55%, retention among neuropeptide genes is markedly higher, at 79%.

**Fig. 2.**
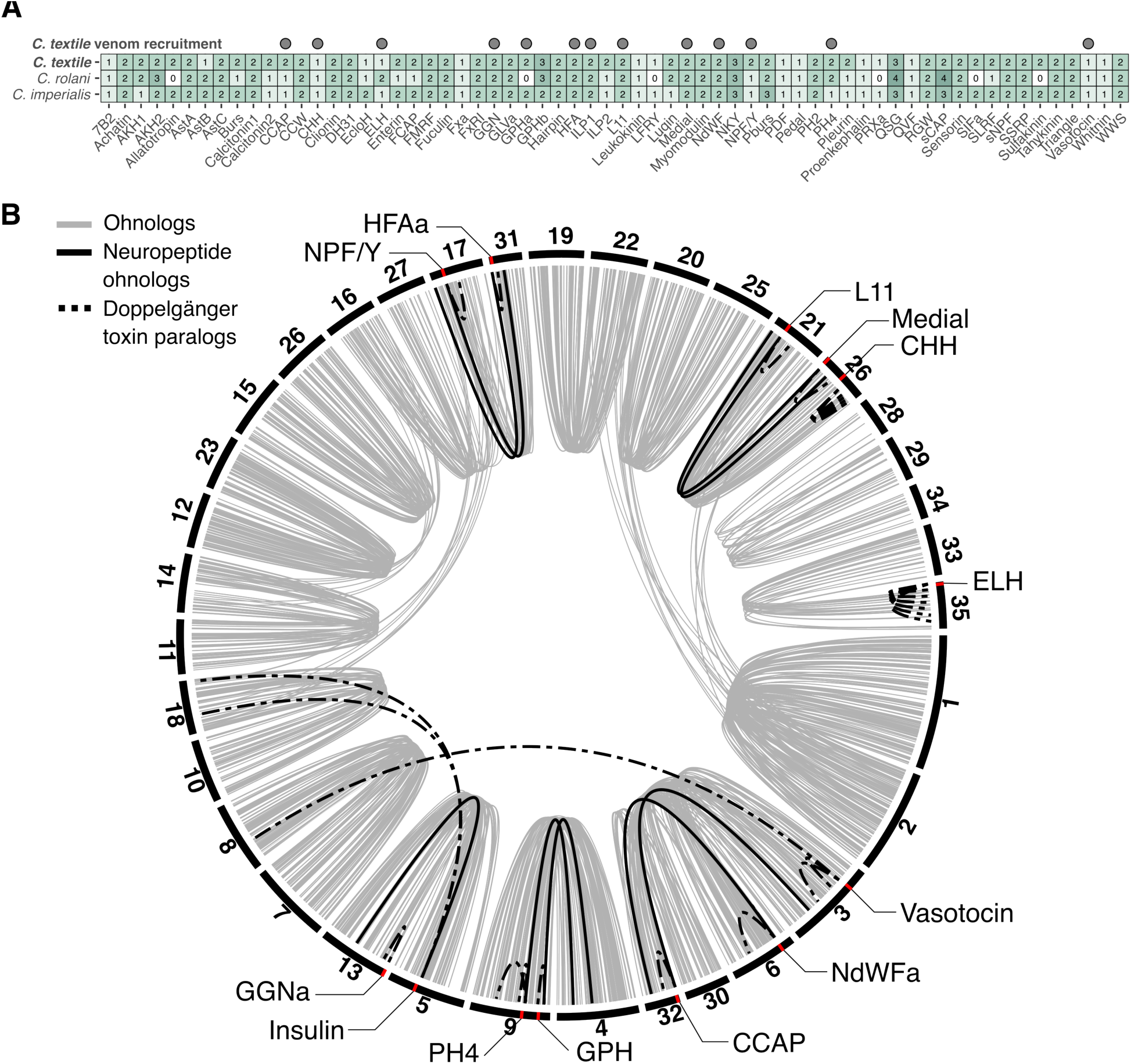
Neuropeptides and doppelgänger toxins in *C. textile.* **(A)** Presence of neuropeptide families in the nerve rings of *C. textile*, *C. imperialis,* and *C. rolani*. Numbers indicate the unique neuropeptide precursors expressed in each species’ nerve ring transcriptomes. Recruited neuropeptide families in *C. textile* are indicated. (**B**) Genomic location of neuropeptides and their corresponding doppelgänger toxins in *C. textile*. The position of the neuropeptide that gave rise to a doppelgänger toxin is indicated by a red line. Neuropeptide ohnologs (when present) are consistently located on homologous chromosomes, reflecting their origin from WGD. Except for insulin and vasotocin, doppelgänger toxins are located on the same chromosome as the neuropeptide ohnolog with which they share the highest sequence similarity. Chromosomal positions of genes have been adjusted for visual clarity.

Having annotated the complement of neuropeptides in *Conus*, we systematically searched for recruited doppelgänger toxins in *C. textile* and other species. Using venom gland transcriptomes from 45 phylogenetically diverse cone snails, we confirmed the presence of both known and several previously undescribed doppelgänger toxin families (**Fig. 3A, Table S6, Data S2-8**). These newly discovered toxins mimic the neuropeptides calcitonin, egg-laying hormone (ELH), FxRIamide, GGNamide, HFAamide, NdWFamide, and neuropeptide KY (NKY). We confirmed the presence of two of these new families (HFAamide and GGNamide) in the venom of *C. textile* (**Figs. S10-11**). The newly identified doppelgänger families have members with high expression (**Fig. S12)** and generally display several characteristics of toxins, including sequence divergence of the toxin encoding region(s) (**Fig. S13**).

**Fig. 3.**
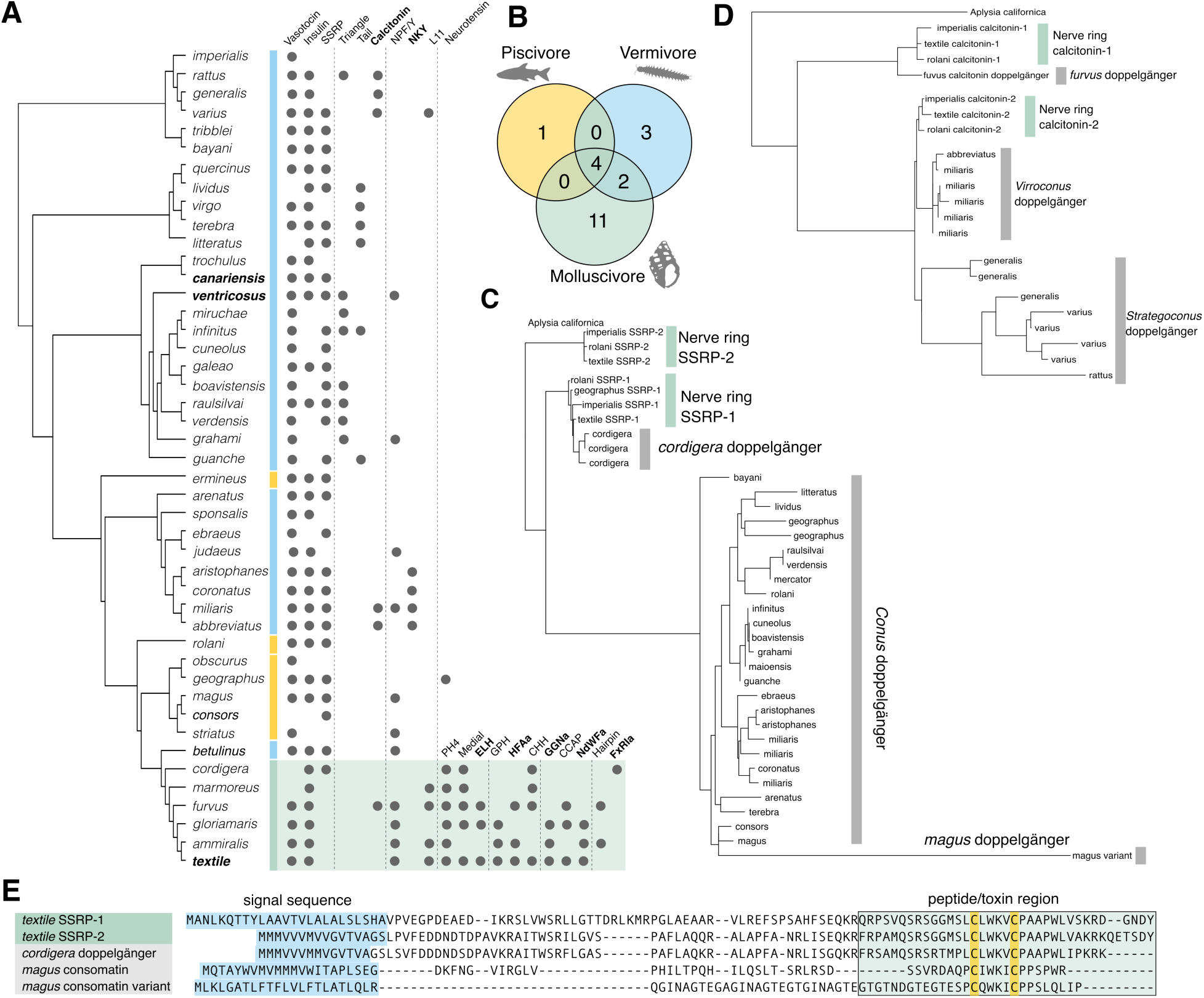
Dynamic recruitment of doppelgänger toxins in *Conus*. **(A)** Phylogenetic distribution of recruited doppelgänger toxins across *Conus* species. Prey preference is indicated by color next to the species: blue for vermivores, orange for piscivores, and green for molluscivores. Bold species have a sequenced genome; bold doppelgänger names denote newly discovered toxins. (**B**) Venn diagram of doppelgänger toxin number and overlap by prey types. (**C**) Maximum likelihood gene tree of (**C**) calcitonin and (**D**) SSRP neuropeptide precursors and their doppelgänger toxins reveal three independent recruitment events each. (**E**) Sequence alignment of *C. textile* SSRP neuropeptide precursors and representative toxin sequences from the three independent recruitment events, with predicted signal peptides (blue) and mature regions (green) highlighted.

Across the 13 doppelgänger toxin families identified in the venom gland of *C. textile*, we found that, with few exceptions, the recruited toxins are located on the same chromosome as the neuropeptide ohnolog to which they show the highest similarity. Genomic distances between the toxin and its corresponding neuropeptide gene vary from 30 kb (e.g., L11) to over 5 Mb (Vasotocin) (**Fig. 2B**). This general pattern of chromosomal colocalization is conserved in the genomes of *C. ventricosus* and *C. canariensis* (**Figs. S14-15**). We identified three doppelgänger toxins in *C. textile* (vasotocin, CHH, and ELH) for which only a single neuropeptide gene is expressed in the nerve ring and detectable in the genome. In all three cases, the doppelgänger toxins are located on the same chromosome as the prevailing neuropeptide gene.

Collectively, these findings reject a simple model in which the WGD provided spare neuropeptide copies that were directly recruited into the venom. Instead, our data support a scenario in which most neuropeptide ohnologs retained their neuropeptide function and subfunctionalized or increased in abundance through higher expression from additional gene copies. In this model, doppelgänger toxins likely arose from more recent molecular events acting on one of the ancestral ohnologs following WGD.

### Doppelgänger toxins were dynamically recruited throughout *Conus* evolution

The identified doppelgänger toxin families display dispersed phylogenetic distribution, revealing a dynamic history of recruitment across *Conus* evolution (**Fig. 3A**):

Four families – conopressins (vasotocin/oxytocin mimics), con-insulins (insulin mimics), consomatins (SSRP mimics), and conoNPY (NPF/Y mimics) – were identified in species from all three predatory types (**Fig. 3A-B**). Conopressins, con-insulins, and consomatins were further identified in each major clade, suggesting that these represent the earliest recruitments. Indeed, these three families were all recruited prior to the emergence of *Conus* and can be found in other venomous Conoidea (**Fig. S16**). These three doppelgänger families further exhibit the most substantial interchromosomal translocation and sequence divergence from the ancestral neuropeptide genes following their recruitment (**Fig. 2B, S14-15**), emphasizing how time can rapidly conceal evolutionary history and underscoring the importance of investigating recently recruited genes to uncover how new genes evolve.

For more recent recruitment events, we identified multiple cases of independent repeated recruitments of the same neuropeptide across different lineages. For example, while consomatins represent a more ancient doppelgänger family that is broadly found across many species, we previously reported that it was lost in molluscivores ^24^. However, with additional data in this study, we detect recently recruited and highly expressed consomatins in the molluscivorous *Conus cordigera* (**Figs. 3C-E, 4C, S17**). Additionally, a distinct consomatin lineage was identified in the fish hunter *Conus magus* (**Fig. 3C-E**). Similarly, also calcitonin and NPF/Y were recruited independently multiple times as demonstrated by gene tree reconstructions (**Figs. 3D, S18-36**).

Finally, we observe a pattern of expansion of doppelgänger toxins in mollusk-hunting cones, with 11 of 20 families exclusively found in these species (**Fig. 3B**). In contrast, nine families were identified in vermivorous species (three of which are exclusive to vermivores), while only members of the four ancestral families discussed above were identified across multiple piscivorous species. To date, only a single case of a doppelgänger toxin exclusively found in a fish-hunting species has been reported. This toxin, contulakin-G from *C. geographus*^25^, mimics the vertebrate hormone neurotensin and evolved from a consomatin-encoding gene (see discussion). Together, these observations reveal a reverse correlation between the number of doppelgänger toxins and the phylogenetic distance to the snail’s prey.

### Reuse of conotoxin 5’UTR in doppelgänger toxin recruitment

The high similarity and close genomic proximity between the recruited doppelgänger toxins and the corresponding neuropeptide genes establish their clear evolutionary relationship. We next investigated the molecular mechanisms underlying neuropeptide gene recruitment to form these toxins.

Though there is high overall sequence similarity between the neuropeptides and their associated doppelgänger toxins, we observed several cases where doppelgänger toxins encode signal sequences identical to established conotoxin gene superfamilies. For example, both the *C. magus* consomatin and *C. furvus* calcitonin doppelgängers have signal sequences belonging to the M-conotoxin superfamily (**Figs. 3E, 4A**), a feature we previously reported for another doppelgänger toxin ^13^. Closer inspection reveals that this similarity extends beyond the signal sequence into the entire 5’UTR (**Fig. 4A**).

**Fig. 4.**
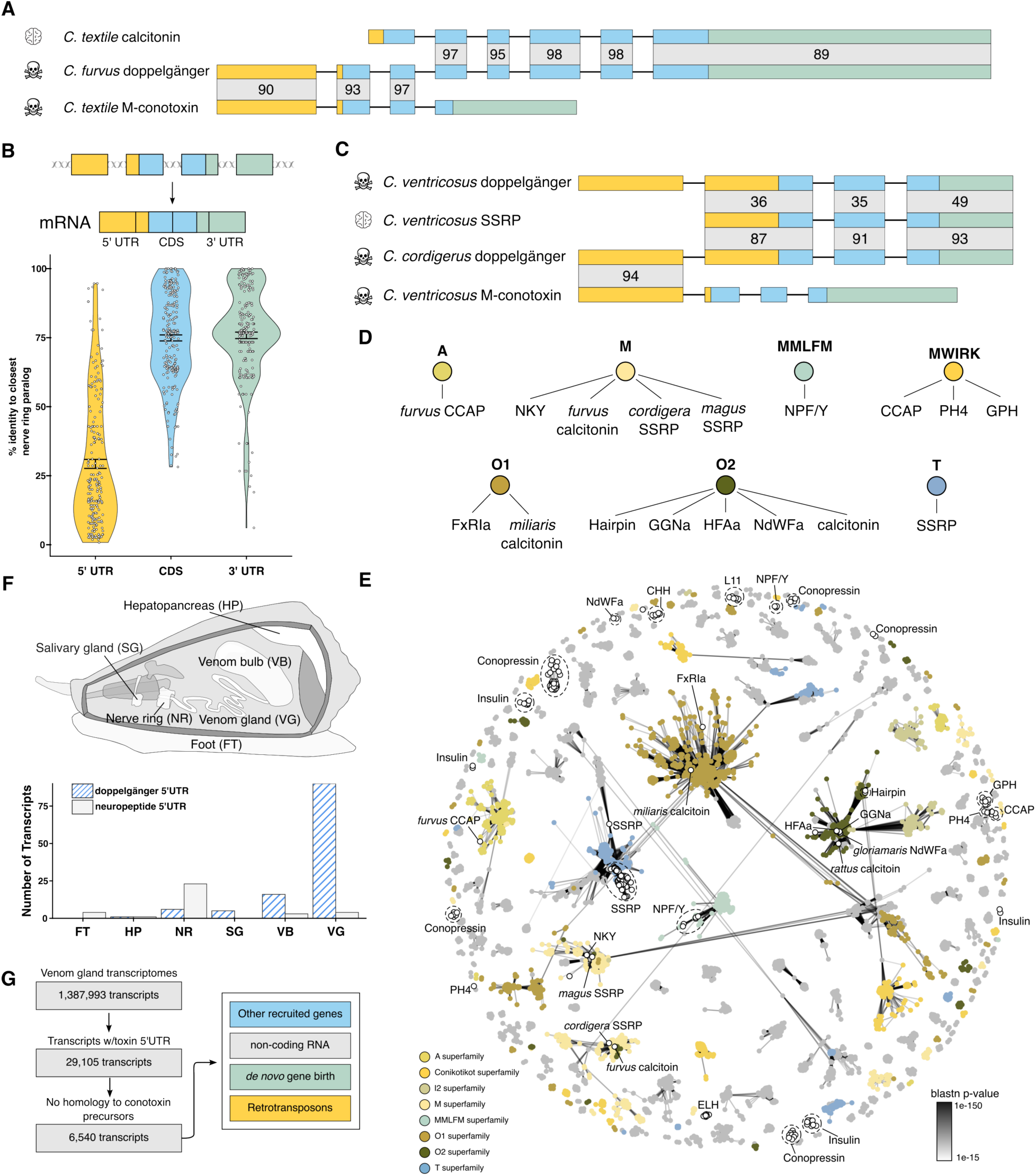
5’UTRs of established conotoxin superfamily are involved in recruiting genes to the venom gland. **(A)** *C. furvus* calcitonin doppelgänger toxins are chimeras of ancestral neuropeptide and M-conotoxin exons, with high similarity to the conotoxin gene from the 5’UTR through the signal sequence (yellow = 5’UTR, blue = CDS, grey = 3’UTR; numbers = pairwise identity). (**B**) When aligned to their ancestral neuropeptide transcript, the 5’UTRs of doppelgänger toxins are significantly more diverged (mean 31 % identity) compared to the coding regions (CDS) and 3’UTRs (mean 75% identity). (**C**) Some doppelgängers, such as SSRP toxins (consomatins), contain an additional conotoxin-derived 5’UTR exon fused with the ancestral neuropeptide gene. Numbers = pairwise identity (color code as shown in panel A). (**D**) Many doppelgänger toxin 5’UTRs trace back to recognized conotoxin family genes. (**E**) CLANS clustering of conotoxin and doppelgänger toxin 5’UTRs shows shared sequence space. Each node represents a single 5’UTR sequence, and the interconnecting lines represent the blastn p-values between the nodes. Major conotoxin superfamilies are colored. Doppelgänger toxin 5’UTRs = white circles with black outlines as annotated in the figure. (**F**) Tissue distribution of transcripts containing doppelgänger toxin or neuropeptide 5’UTRs across *C. textile* tissues. (**G**) Of the transcripts expressed in the venom gland, ∼ 2% contain conotoxin 5’UTRs. 22% of these lack a conotoxin ORF, and fall into other categories: other recruited genes, non-coding RNA, likely *de novo* evolved genes, and retrotransposons.

Given that the 5’UTR and upstream genomic regions are known to influence tissue-specific gene expression ^26^, this suggests a mechanism of regulatory element co-option, whereby doppelgänger toxins acquire venom gland-specific expression by recombining with 5’UTRs and potentially upstream regulatory elements of preexisting conotoxin genes. Strikingly, although only a subset of doppelgängers possesses a conotoxin signal sequence, nearly all doppelgänger 5’UTRs closely resemble the 5’UTRs of established conotoxin gene superfamilies (**Figs. 4DE**). Consistent with this, their 5’UTRs share very little sequence similarity (an average of 31%) with the most closely related neuropeptide transcript, compared to an average of 75% identity of the coding sequences (CDS) and 3’UTRs (**Fig. 4B**). This pattern suggests that the 5’UTRs of doppelgänger toxins frequently arise from existing conotoxins during recruitment, likely facilitating venom gland-specific expression. Supporting this, conotoxin-like 5’UTRs are expressed almost exclusively in the venom gland compared to other tissues (**Fig. 4F**). An exception to this pattern is seen in con-insulins and most consomatins, where all regions of the transcripts show similar levels of divergence and lack any obvious resemblance to conotoxin 5’UTRs, possibly because their more ancient recruitment has obscured any traces of their original regulatory structure (**Fig. S37**).

Several cases clearly illustrate the 5’UTR-mediated recruitment mechanism. For example, the 5’UTR of the recently recruited consomatin gene in *C. cordigera* shares 94% identity with the 5’UTR of M-superfamily toxins, while the remainder of the sequence is ∼90% identical to the SSRP-2 neuropeptide gene (**Fig. 4C**). NKY doppelgängers and the calcitonin doppelgänger in *C. furvus* similarly share M superfamily 5’UTRs (**Figs. 4A,D-E**). Cono-NPY doppelgängers contain 5’UTRs that align with the MMLFM toxin superfamily, while O2-superfamily 5’UTRs are found in several other doppelgänger genes (i.e., doppelgängers of hairpin, GGNamide, HFAamide, and a subset of NdWFamide and calcitonin) (**Figs. 4D-E**). O1-superfamily 5’UTRs are found in doppelgängers of FxRIamide and the *Conus miliaris* calcitonin doppelgänger, and CCAP, GPH, and PH4 all share a common 5’UTR (**Figs. 4D-E**).

Upon further examination, we observe that this recruitment mechanism is not limited to doppelgängers but can be observed in other toxins. For instance, conoporins, which are related to molluscan pore-forming proteins, appear to have been recruited through fusion with an O1-superfamily 5’UTR. Similarly, we previously observed chimeric toxins that contain mature toxin domains from the T-superfamily but have 5’UTR and signal sequence from the L-superfamily ^11^.

Moreover, we identified additional non-conotoxin transcripts with conotoxin-derived 5’UTR, including genes encoding retrotransposons, other genes, unknown open reading frames (some with predicted signal sequences), and likely non-coding RNAs, which may play functional roles in the venom system (**Fig. 4G**). Notably, several conotoxin transcripts contain remnants of degraded retrotransposons (**Fig. S38**).

Collectively, these results support a model in which new paralogs of neuropeptide genes are recruited into the venom system through recombination with existing conotoxin 5’UTRs. This regulatory switch likely confers the molecular signatures necessary for venom gland expression, representing a key mechanism in the evolutionary expansion and functional diversification of cone snail toxins. Here, we propose the term ‘toxi-driver’ for these 5′ UTR exons, which appear to be repeatedly reused through recombination with neuropeptide genes to generate novel doppelgänger toxins.

### Intron phase and repeat landscape facilitate exon shuffling in doppelgänger toxin evolution

Mechanistically, the observed recruitment pattern is consistent with exon shuffling— recombination of pre-existing genic fragments to form novel genes. Local duplications, translocations, inversions, and ectopic recombination can all lead to a novel chimeric gene containing the 5’ region of one gene and the 3’ region of another gene. Similar processes have been observed in other organisms, including humans and *Drosophila*^27^.

Several of these mechanisms are known to be greatly facilitated by TEs^28^. The genome of *C. textile* contains ∼60% TEs, with similar numbers reported for *C. ventricosus, C. canariensis,* and other neogastropods. This is notably higher than in other mollusks such as the caenogastropod *P. canaliculata*^29^ (**Fig. 5A**). There is almost no overlap of the TE families in *Conus, P.canaliculata,* and another gastropod belonging to the Conoidean superfamily, *Pymorhynchus buccinoides*. Even within *Conus,* most TEs appear to be species-specific (**Fig. 5B**).

**Fig. 5.**
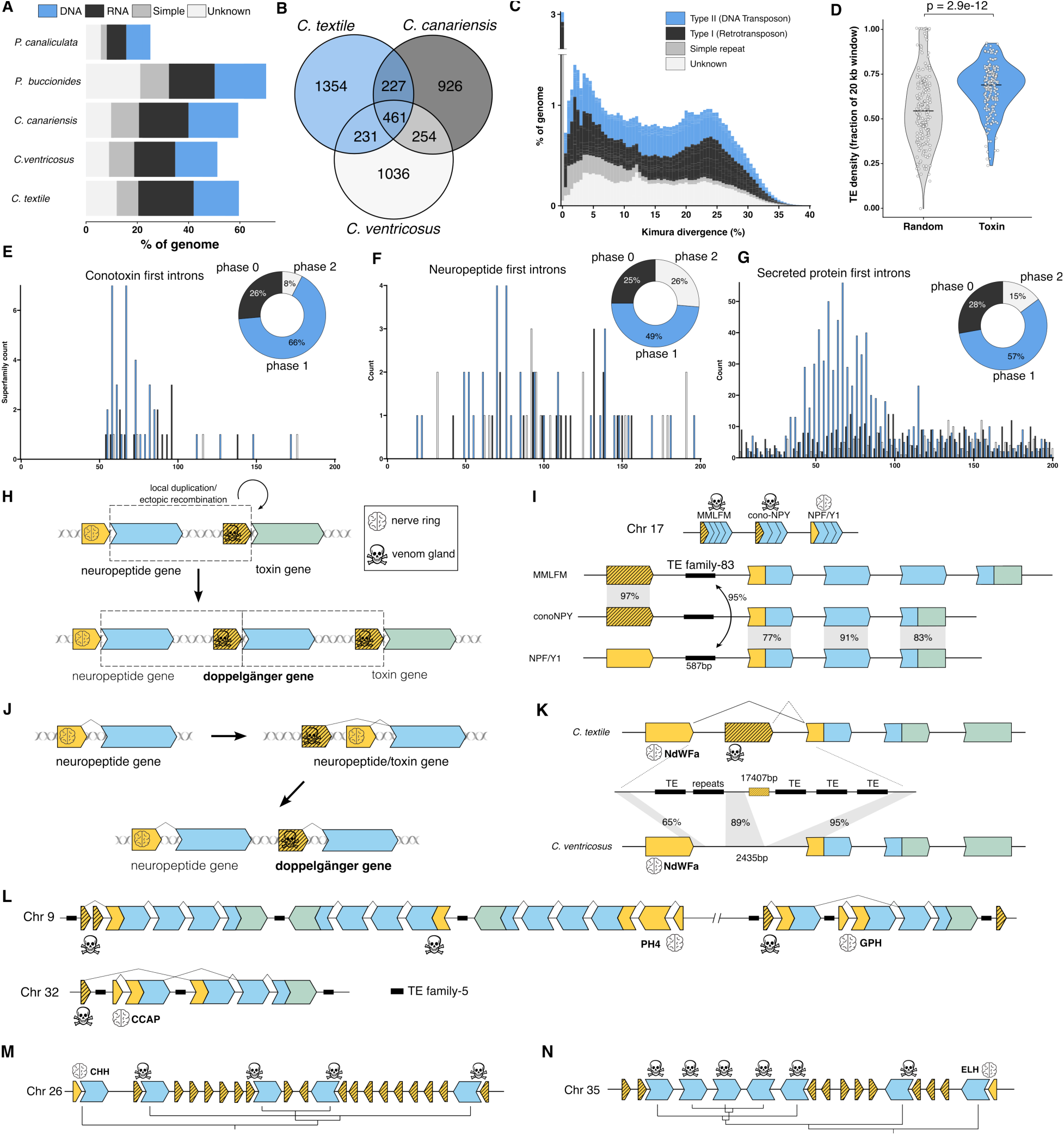
Genomic mechanisms underlying the evolution of doppelgänger toxins. **(A)** Cone snails and other neogastropods possess a higher proportion of transposable elements (TEs) and repeat elements compared to other mollusks (e.g., *P. canaliculata*). (B) ∼ 98 % of *Conus* TEs are unique relative to the conoidean *P. buccionides*, with many being species-specific. (C) The *C. textile* TE landscape reveals a recent expansion of simple repeats and multiple historical TE bursts. (D) Toxin-flanking regions (±10 kb) show significantly higher TE density than random genomic regions (p = 2.9e-12, Wilcoxon rank sum test). (E-G) Secreted proteins, conotoxins, and neuropeptides show a strong phase-1 intron bias 50-100 bp into the coding region. (H) Model of doppelgänger toxin recruitment via ectopic recombination. (I) ConoNPY doppelgänger genes likely arose via exon shuffling between MMLFM toxins and NPF/Y neuropeptides, facilitated by family-83 TEs located in the first intron. (J) Doppelgänger toxins can also be recruited via alternative splicing and subsequent gene duplication as a mode to escape adaptive conflict. (K) The NdWFamide doppelgänger toxin arose via transposition of a new exon into the first intron of the neuropeptide gene, producing an alternative splice variant, likely facilitated by the transposition of TEs in the first intron. (L) PH4, GPH, and CCAP doppelgänger toxins share a 5’UTR. While PH4 doppelgängers exists as separate genes located next to the neuropeptide gene, both GPH and CCAP doppelgängers arise from alternative splicing. (M-N) Local duplications generate clusters of CHH and ELH doppelgängers near their neuropeptide origins. The second exon (blue) contains the complete CDS and 3’UTR.

The TE landscape of *C. textile* reveals both a recent expansion of simple repeats as well as historical bursts of retrotransposons and unclassified elements (**Fig. 5C**). Furthermore, we observe increased TE density in regions surrounding toxin genes—opposite to the pattern seen near other genes (**Fig. 5D, S39-40**), strongly suggesting an important role for TEs in driving toxin evolution. The most common TE family in the *C. textile* genome, the retrotransposon family-5 (186,000 copies), can be found in a wide range of conotoxin transcripts, including nine different superfamilies (A, Conikotikots, Conohyal, I4, L, M, O1, S, and T) (**Fig. S38**). Notably, this same TE is also present in the genomes of another molluscan species, *Littorina saxatilis*, including within annotated gene regions (**Fig. S41**).

As discussed, we observed several cases where exon shuffling occurred with a fusion point within the CDS. Such cases depend on intron phase compatibility among the fusion proteins to result in a functional gene. Furthermore, to retain biological activity, the fusion should preferentially happen such that functional regions (e.g., peptide) of the original proteins are not separated. Conotoxin genes show a strong bias for the first intron to be phase 1 (**Fig. 5E**) and are located 50-100 nucleotides into the CDS, corresponding to the typical length of signal sequences (15-30 amino acids). The same pattern is present in neuropeptide genes (**Fig. 5F**) and secreted proteins in general (**Fig. 5G**), but not in non-secreted proteins (**Fig. S42**). Thus, a strong preference for phase 1 introns located adjacent to signal sequence encoding exons creates opportunities for exon shuffling to happen within the coding region of proteins at functional domain boundaries, as seen in several doppelgänger toxins.

### Mechanisms of doppelgänger toxin recruitment

Doppelgänger toxins appear to have been recruited through several distinct mechanisms across different gene families. Several molecular mechanisms are consistent with the patterns we observe, including cases we can reconstruct in detail—particularly for more recent recruitments—while others remain unresolved due to a combination of sequence divergence and imperfect genomic resolution. In this section, we outline three primary mechanisms by which doppelgänger toxins appear to have evolved: ectopic recombination, alternative splicing, and gene duplication following initial recruitment.

#### Ectopic recombination

Ectopic recombination refers to crossover events between homologous sequences located at non-allelic positions, often facilitated by TEs^28^. We identify at least one clear case in which the origin of a doppelgänger toxin appears to result from such an event, involving TE-mediated recombination (**Fig. 5H, Data S9**). This case involves family-83 TEs (Unclassified), which are highly abundant in the *C. textile* genome (42,794 copies). *C. textile* encodes the neuropeptide NPF/Y in two orthologous loci on chromosomes 17 and 31 (**Fig. 2B**). The doppelgänger conoNPY gene is located ∼2 Mb upstream of the NPF/Y ohnolog on chromosome 17 and shares a 5′ UTR with toxins from the MMLFM superfamily.

Only a single MMLFM toxin gene is present in the *C. textile* genome, situated ∼0.2 Mb upstream of conoNPY (**Fig. 5I**). Sequence comparisons reveal that the first exon and part of intron 1 of the MMLFM gene are nearly identical to those of conoNPY, while the rest of conoNPY matches NPF/Y. Both the MMLFM and NPF/Y loci contain family-83 TE sequences with near-perfect identity around the presumed recombination breakpoint where conoNPY switches from resembling MMLFM to NPF/Y. This suggests that ectopic recombination at TE insertion sites mediated a local duplication and fusion producing the conoNPY doppelgänger gene. This mechanism couples gene duplication and neofunctionalization into a single event (**Fig. 5H**). The similarities between the MMLFM and nonoNPY loci extend ∼400 bp upstream of the 5’UTR (**Fig. S43**).

#### Alternative splicing

In several instances, doppelgänger toxins have emerged through alternative splicing of neuropeptide precursor genes (**Fig. 5J**). One well-resolved example is the NdWFamide doppelgänger family, found only in mollusk-hunting species. In *C. textile*, the NdWFamide gene shows an alternative venom gland-specific transcript that uses a novel exon within a greatly expanded intron (∼17 kb) (**Fig. 5K**). This exon encodes a unique 5′ UTR not found in the nerve ring transcriptome nor in the genome of *C. ventricosus* that lacks NdWFamide doppelgängers and has a much shorter intron (∼2 kb). The expanded intron in *C. textile* is enriched in TEs, strongly implicating TE activity in exon emergence and tissue-specific expression of the doppelgänger variant.

Additional examples include the Prohormone-4, CCAP, and GPH doppelgängers, all of which share similar 5′ UTR-encoding exons also found in some MWIRK conotoxins (**Fig. S44**). Among these, CCAP and GPH appear to be generated by alternative splicing in *C. textile*, whereas Prohormone-4 doppelgängers are encoded by distinct genes (**Fig. 5L**). GPH and Prohormone-4 are located on chromosome 9, and the GPH doppelgänger may have recombined with a 5′ UTR exon from the Prohormone-4 family. TE family-5 is among the most abundant TEs in regions surrounding the GPH, CCAP, and Prohormone-4 genes (excluding simple repeats). The ±10 kb windows around these loci contain 68%, 62%, and 79% TEs, respectively, further supporting the role of TE-driven regulatory and structural changes. Although CCAP doppelgänger transcripts were only detected in mollusk hunters in our analysis, this toxin has also been isolated from a vermivore^30^, suggesting a possible independent recruitment event.

#### Gene duplication following recruitment

For some families, the mechanism of initial recruitment remains unclear. However, gene duplication following recruitment appears to have played a major role in the expansion of several doppelgänger families.

For instance, Prohormone-4 doppelgängers are represented by multiple gene copies (**Fig. 5L**). In contrast, the more recently recruited GPH and CCAP doppelgänger toxins, which share their 5’UTRs with Prohormone-4, are generated through alternative splicing. These examples suggest that tissue-specific isoforms may create evolutionary adaptive conflicts, which sometimes are resolved through gene duplication and subfunctionalization (**Fig. 5J**).

Two additional families, CHH and ELH, illustrate how recombination with a novel 5’UTR-encoding exon (of unknown origin in these case) led to the emergence of doppelgänger toxins from single-copy neuropeptide genes. In both cases, repeated rounds of local duplication have resulted in tandemly arrayed toxin genes (**Figs. 5M-N**), and the 5’UTR-encoding exon itself appears to have undergone rapid expansion, producing redundant copies.

Collectively, these findings reveal that TEs play a central role in facilitating the recruitment of doppelgänger toxins. Through a combination of ectopic recombination, alternative splicing, and local duplication—often mediated by TE activity—cone snails have repeatedly co-opted neuropeptide genes for venom production in a lineage-specific and mechanistically diverse manner, illustrating the dynamic interplay between genome architecture and molecular innovation in venom evolution.

## Discussion

The origin of novel genes and their functional integration into biological systems is a central problem in evolutionary biology. Venoms, due to their dynamic and often accelerated evolution of toxin-encoding genes that confer adaptive advantages in predator-prey interactions, offer exceptional systems for investigating the molecular mechanisms underlying gene recruitment, neofunctionalization, and diversification. The unparalleled diversity of toxins in cone snail venom makes them an ideal system for investigating these phenomena.

In this study, genomic and transcriptomic analyses of *C. textile* revealed a remarkably rich repertoire of doppelgänger toxins that mimic molluscan neuropeptides. Many of these likely evolved as adaptations to a molluscivorous lifestyle that emerged in the early Miocene, ∼18 million years ago^12^. Due to this relatively recent origin, these toxins represent a powerful system to study how genes can be co-opted from endogenous neuropeptide signaling roles to venoms used in prey capture.

We demonstrate that *Conus* venoms contain a far more diverse and abundant array of doppelgänger toxins derived from neuropeptide genes than previously recognized. These toxins have been repeatedly gained, lost, and independently recruited into the venom system across lineages. A key mechanistic insight is that most doppelgänger genes harbor 5’UTRs that are distinct from those of their neuropeptide progenitors but homologous to known conotoxin 5’UTRs. These venom-specific 5’UTRs – termed toxi-drivers – are absent from non-venom tissues, strongly supporting a model in which exon shuffling, specifically recombination between venom-specific regulatory exons and neuropeptide-encoding exons, drives the recruitment of genes into the venom system. Importantly, these recombination events address a longstanding issue in models of gene duplication: how newly duplicated genes escape pseudogenization. By acquiring regulatory elements that confer expression in the venom gland, doppelgänger toxins gain instant expression in the venom, thereby bypassing the need for a gradual accumulation of regulatory changes.

Multiple genomic features facilitate this process. Both conotoxins and neuropeptides show a strong enrichment of phase 1 introns adjacent to their signal peptide-encoding regions, creating intron phase-compatibility for exon fusion. Additionally, the *C. textile* genome shows a substantial expansion of TEs, including families specific to *Conu*s. We find evidence for retrotransposons recombining with conotoxins, potentially providing opportunities for later recombination events, TE enrichment around toxin loci, and TEs flanking the breakpoints of recombination events. In the case of conoNPY, for instance, highly similar TEs between neuropeptide and MLSML conotoxins support TE-mediated ectopic recombination as a recruitment mechanism.

Although exon shuffling and TE-mediated recombination are predominant, not all doppelgänger toxins arise through the same pathway. In several cases, such as for the neuropeptides NdWFamide, CCAP, and GPH, alternative splicing generates functionally distinct splice variants of the same gene expressed in the nervous system and venom gland. These cases may represent intermediate “moonlighting” states, in which a single gene possesses dual functionality prior to duplication and subfunctionalization that resolve the adaptive conflict. Once recruited, several doppelgänger toxins, such as Prohormone-4, CHH, and ELH toxins, undergo repeated rounds of local duplication, which mirrors what is seen in many other conotoxin families.

Our findings add to the growing evidence that cone snail venom diversity arises through multiple, complex mechanisms. However, these are not the only mechanisms involved in creating toxin diversity in cone snails. Exonization of previously non-coding loci has happened in the SSRP doppelgänger from *C. geographus*^31^. And further duplications of doppelgänger genes provide opportunities for continued evolutionary innovation. For example, members of the SSRP doppelgänger family neofunctionalized in some cases to target entirely new receptors, such as contulakin-G, a neurotensin receptor agonist from *C. geographus*, and αC-PrXA, a muscle nicotinic acetylcholine receptor (nAChR) antagonist from *Conus parius* (**Fig. S45**).

Further, it is important to note that although these appear to be exceptions in *Conus*, not all doppelgänger toxins evolved from a preexisting neuropeptide precursor. For instance, the abovementioned contulakin-G, a neurotensin doppelgänger, evolved from the SSRP doppelgänger family. Likewise, conorfamides, several of which possess a C-terminal RFamide motif, likely did not evolve from the RFamide neuropeptide precursor.

The striking expansion of doppelgänger toxins in molluscivorous cone snails raises the question: why this lineage? Our genome-wide comparisons found no major structural differences between molluscivores and vermivores. We propose that the key factor driving this expansion is phylogenetic relatedness to prey. Molluscan prey, being closely related to cone snails, possess neuropeptide receptors that recognize cone snail neuropeptides. In some cases, cone snails even prey on other cone snails (**Movie S3-4**), meaning their targets can be genetically identical or nearly so. This likely receptor-ligand cross-reactivity would make neuropeptide-derived toxins functionally effective, offering an immediate selective advantage and facilitating evolutionary retention. This hypothesis aligns with the observed inverse relationship between the number of doppelgänger toxins and the phylogenetic distance to prey, with fish-hunters possessing the fewest doppelgänger toxins. An alternative or additional hypothesis is that the successful capture of molluscan prey requires unique toxins—including doppelgängers—as we have previously demonstrated for several non-doppelgänger toxins^11^.

Together, our findings support a generalizable model of toxin evolution in which the modular architecture of genes, in combination with recombination, splicing events, and TE activity, enables the rapid evolution of new genes. This framework aligns with the “genes in pieces” model proposed by Gilbert^32^ and reinforces the idea that gene architecture itself facilitates the evolution of complex traits. We propose that the strategies of doppelgänger toxin emergence represent broadly applicable mechanisms of evolutionary innovation that will likely be uncovered across other venomous lineages as genomic resources continue to expand.

## Materials and Methods

### Sample collection

The *Conus textile* specimen was collected by SCUBA. The snail was dissected, and tissues were stored in RNAlater for RNAseq and HiC and flash-frozen in liquid nitrogen for PacBio sequencing.

### Transcriptome sequencing

RNA was extracted from the venom gland, circumoesophageal nerve ring, hepatopancreas, venom bulb, foot, and salivary gland using Direct-zol RNA extraction kit (Zyme Research, Irvine, CA, USA) following the manufacturer’s protocol. Extracted RNA was submitted to the Huntsman Cancer Institute’s High Throughput Genomics Core Facility, where library preparation with the Illumina TruSeq Stranded mRNA Library Preparation kit with polyA selection was performed. Sequencing libraries were chemically denatured in preparation for sequencing. Following transfer of the denatured samples to an Illumina NovaSeq X instrument, a 151 x 151 cycle paired end sequence run was performed using a NovaSeq X Series 10B Reagent Kit.

For assembly, adapter trimming of de-multiplexed raw reads was performed using fqtrim^33^, followed by quality trimming and filtering using prinseq-lite^34^. Error correction was performed using the BBnorm ecc tool, part of the BBtools package. Trimmed and error-corrected reads were assembled using Trinity (version 2.2.1)^35^ with a k-mer length of 31 and minimum k-mer coverage of ten. Assembled transcripts were annotated using a blastx^36^ search (E-value setting of 1e-3) against a combined database derived from UniProt and conoserver^37^. Transcripts per million (TPM) counts were generated using the Trinity RSEM^38^ plugin (align_and_estimate_abundance) and expression data analysed using the trinity utilities abundance_estimates_to_matrix and contig_ExN50_statistic.

### DNA isolation and shearing for PacBio sequencing

PacBio HiFi sequencing proved to be challenging for this species. To increase sequencing yields, three different libraries were generated following the PacBio protocol for whole-genome HiFi library preparation with the SMRTbell® prep kit 3.0.

Flash-frozen tissue from foot and columnar muscle was cryogenically powdered with a cryoPREP® (Covaris) in batches of 115-250 mg. An initial DNA isolation was performed using the Nanobind® tissue kit (PacBio), following the “Extracting HMW DNA from black mystery snail tissue using Nanobind® kits” Procedure & checklist. The isolated DNA had low purity, indicating probable polysaccharide contamination, and was not processed further. DNA isolation was therefore repeated using an in-house protocol consisting of CTAB buffer tissue lysis followed by DNA purification with Genomic-tip 100/G gravity-flow columns (QIAGEN). This DNA was of high purity and integrity and was processed into PacBio HiFi library 1.

One additional DNA isolation was performed following the in-house CTAB protocol but with the addition of sorbitol-washes of the powdered tissue in order to remove possible long-read sequencing inhibitors^39^. This DNA was processed into PacBio HiFi libraries 2 and 3, and the detailed protocol is available on https://www.protocols.io/private/5E72D9E3558211F0AD2A0A58A9FEAC02.

Prior to shearing, DNA was treated with SMRTbell® cleanup beads (PacBio, library 1) or with the Megaruptor® 3 DNAFluid+ kit (Diagenode, libraries 2 and 3). DNA was sheared into an average fragment size of 12-28 kb using the Megaruptor® 3 Shearing kit (Diagenode).

Quality control of DNA yields, purity, and integrity was performed using the Qubit BR and HS DNA quantification assay kits (Thermo Fisher), NanoDrop (Thermo Fisher), and Fragment Analyzer (DNA HS 50kb large fragment kit, Agilent Technologies).

### PacBio HiFi library preparation and sequencing

HiFi libraries were size-selected on High Pass PlusTM gel cassettes on the BluePippinTM instrument (Sage Science). The final library size was determined on the Fragment Analyzer (DNA HS 50kb large fragment kit, Agilent Technologies). Sequencing was performed by the Norwegian Sequencing Centre on the PacBio Revio instrument using the Revio™ polymerase kit and one 25M SMRT cell for each library. Circular consensus sequence (CCS) HiFi reads were generated using the CCS pipeline (SMRT Link version 10.2.0.133434). Due to sub-optimal yields, sequencing was performed on three libraries with variable parameters for library preparation (**Table S8**).

The first library had an average insert size of 19 kb after a standard BluePippinTM size-selection cutoff of 10 kb, and sequencing yielded 24.7 Gb of HiFi reads. For non-model species, outputs of >60Gb HiFi reads are expected for successful runs (Wellcome Sanger Institute, personal communication). Furthermore, short polymerase read length (**Table S8**) and base yield density distribution (data not shown) indicate poor polymerase performance and early polymerase termination events.

To remove possible polymerase inhibiting contaminants, DNA was isolated again, including the sorbitol wash step described above. HiFi library 2 was prepared as described for library 1, with an additional cleanup of the sheared DNA using the DNeasy PowerClean Pro Cleanup Kit (QIAGEN). The final library had an average insert size of 26 kb and yielded 15.6 Gb of HiFi reads, with shorter polymerase read length and increased polymerase termination events. The poorer performance of library 2 compared to library 1 suggests that the low sequencing output may not be caused by the presence of sequencing-inhibiting contaminants. Furthermore, none of the DNA quality controls indicate the presence of impurities or DNA damage that could explain poor sequencing performance.

To generate higher sequencing output despite sub-optimal polymerase performance, we repeated the library preparation as described for library 2, with a shorter insert size. The DNA was sheared with higher speed code settings on the Megaruptor® 3, and the cutoff for size-selection on the BluePippinTM was set at 6 kb; the final library had an average insert size of 13 kb. While the polymerase read length and base yield density only marginally improved compared to library 1, library 3 yielded more than twice the amount of HiFi reads, for a total of 38.4 Gb HiFi yield.

### HiC library preparation, and sequencing

Approximately 80 mg of tissue was used for the construction of a library for Hi-C sequencing employing the Arima High coverage Hi-C kit (L/N2309050008; part no. A160162 v01; Arima Genomics, CA, USA). This library was constructed using cross-linked and proximally-ligated DNA, fragmented to 550–660 bp using an ultra-sonicator (M220; Covaris, USA), then assessed for quality using the TapeStation (as described above) and paired-end sequenced in one lane of a Nova Seq X (10B, 300 cycles; Illumina, USA), with short reads being stored in the FASTQ format.

### Genome assembly and scaffolding

Cutadapt version 4.4^40^ was used to remove remaining adapter sequences from HiFi reads before the trimmed HiFi reads were assembled using hifiasm version 0.19^41^ with default settings, incorporating Hi-C. Haplotigs and overlaps were purged with Purge_Dups v1.2.6^42^ based on coverage of HiFi reads mapped with minimap2^43^ using the maphifi option and self-aligned contigs aligned with minimap2 using the xasm5 option. This yielded a primary assembly consisting of 35102 contigs with an N50 of 187.7 kb and a total length of 3.5 Gb. For scaffolding the assembly, Hi-C reads were mapped to the primary contigs with chromap v0.2.6 (r486)^44^, converted to bam format with SAMtools v.1.17^45^, and scaffolded with YAHS v2.1^46^. Hi-C reads were then mapped to the scaffolded assembly using chromap and a contact map generated with Paired REad TEXTure Mapper (PretextMAP) v0.1.9 (https://github.com/wtsi-hpag/PretextMap) and visualized with PretextView v0.2.5 (https://github.com/sanger-tol/PretextView). Due to the low contiguity of the scaffolded assembly (13,831 scaffolds; N50 1,508,245; total length 3.5 Gb) we aligned and further scaffolded it against *Conus canariensis* (NCBI accession JAMYXO000000000.1;^20^) with the scaffolding function in RagTag v2.1.0^47^. We then mapped H-C reads to the resulting RagTag-scaffolded assembly with Chromap to correct any erroneous scaffolding with YAHS, and used PretextMAP and PretextView to further manually inspect and correct the scaffolded assembly. Contiguity was calculated using Quast version 5.2.0^48^ while completeness was assessed by comparing against universally conserved single-copy orthologs from Mollusca (mollusca_odb10) and metazoa (metazoan_odb10) using BUSCO version 5.0.0^49^.

### Annotation of repeats and transposable elements

RepeatModeler v2.0.5^50^ and RepBase were used to contruct a *de novo* repeat library for *C. textile* (and other genomes reanalyzed here). This database was filtered for *bone fide* genes (sometimes RepeatModeler will interpret an expanded gene family as a repetitive element) using the predicted proteome of other cone snails^20–22^, as well as an expansive set of conotoxin sequences^11^. We used DIAMOND v2.1.9^51^ with an e-value of 1e-5 to identify sequences in the *de novo* repeat library with significant similarity to protein-coding genes. 876 repeat elements had significant hits, which were manually inspected and removed if they indeed encoded proteins. The filtered repeat database was used to annotate and soft mask the genome assembly with RepeatMasker v4.1.9^52^. All repeat families were further annotated using Terrier v.0.2.0^53^.

The same TE annotation approach was used for *Conus ventricosus, C. canariensis, Pomacea canaliculata,* and *Phymorhynchus buccinoides*. To identify overlapping TE families, we used mmseqs2^54^ easy-cluster with settings -min-seq-id 0.5 -c 0.8 --cov-mode 1. The output was parsed, and each cluster was characterized by the species represented.

The Kimura TE landscape was generated by parsing the output from RepeatMasker for *C. textile* and calculating the histograms of sequence divergence for each of the different TE families. The classifications were based on the annotation generated from Terrier.

### TEs surrounding conotoxins

To calculate the TE density surrounding toxins, we used bedtools v2.31.0 to extract a window spanning 10k bp on either side of the ToxCodAn-genome toxin annotations. The TE coverage of these regions was calculated with bedtools. We further used bedtools shuffle to randomly position the toxins across the genome and calculated the TE density surrounding these randomly shuffled toxins for comparison. A similar analysis was performed with the braker annotated genes.

### Gene and conotoxin annotation

We mapped all generated RNA-seq data to the soft-masked genome using minimap2 v2.24.0^43^ and converted to bam files using samtools v.1.16.0 ^45^. Additionally, a database of predicted *Conus* proteins, conotoxins, and UniProtKB entries (Uniprot Consortium, 2024; downloaded January 2025) was used in Braker v.3.0.8 in ETP mode^55^. The completeness of the gene set was assessed using BUSCO^49^ in ‘protein mode’ (‘mollusca’ and ‘metazoa’ lineage) and OMark^56^. Genes and inferred proteins were annotated using Eggnog-mapper v.2.1.9^57^.

We identified toxin-encoding transcripts from the assembled venom gland transcriptome using an in-house database of known toxin families as queries for blastn searches. Additionally, we identified toxin-encoding genes in the genome assembly using ToxCodAn-genome^58^ using a custom set of conotoxin CDS from^11^.

### Macrosynteny analyses

Proteomes and gff-files from *P. canaliculata, C. ventricosus,* and *C. canariensis* were downloaded from relevant repositories. Reciprocal best blast hits were identified between organism pairs: *P. canaliculata – C. ventricosus, C. ventricosus – C. canariensis,* and *C. canariensis – C. textile* as implemented in rbhxpress (https://github.com/SamiLhll/rbhXpress). The combined set of reciprocal best blastp hits was combined and macrosynteny was visualized using the macrosyntR library in R^59^.

### Neogastropod species tree

The proteomes of *C. textile, P. canaliculate, C. ventricosus,* and *C. canariensis* were used as inputs to OrtoFinder v.2.5.4^60^ and the inferred orthologs were used for species tree inference using STAG^61^. The time was calibrated using makeChronosCalib as implemented in the R library ape v.5.8.1. The divergence times were obtained from TimeTree^62^.

### Neuropeptide annotation

To identify neuropeptide precursor sequences from the *Conus* circumoesophageal nerve ring transcriptomes, we performed a relaxed tBLASTn v.2.14.1^36^ search with e-value 1e-3. As queries, we used a large reference database of neuropeptide precursors from^63–66^. The hits were visually inspected to confirm they possessed the characteristics of the query, such as putative cleavage sites and conserved residues in the peptide-encoding regions. The identified precursors were analyzed for the presence of signal sequences using SignalP v6^67^. The neuropeptide precursor sequences can be found in **Data S1**.

### Homolog Chromosome Identification

To identify homologous chromosomes resulting from whole-genome duplication (WGD), we first identified ohnologs by performing a self-query of the predicted proteomes. We used a modified version of rbhXpress (https://github.com/SamiLhll/rbhXpress) to detect reciprocal best hits: for each query protein, the top two DIAMOND hits were retrieved, excluding the query itself. This yielded an all-against-all ‘homolgy matrix’ retaining up to the best two matches per protein. Macrosynteny relationships were then visualized using the macrosyntR package ^59^ and the circlize library in R.

### Ohnolog retention in *C. textile*

Orthogroups shared between *Conus textile*, *Pomacea canaliculata*, *Conus ventricosus*, and *Conus canariensis* were identified using OrthoFinder, as described above. To assess ohnolog retention in *C. textile*, we extracted orthogroups containing at least one gene from both *C. textile* and *P. canaliculata*. Within this subset, we calculated the proportion of orthogroups in which *C. textile* possesses two or more genes, interpreted as retained duplicates.

Although this approach does not distinguish between true ohnologs and lineage-specific gene expansions, it provides a useful approximation of the degree of gene duplication retained in the *C. textile* genome following whole genome duplication.

### Doppelgänger peptide identification

We searched for instances of neuropeptide recruitment to the venom glands to form doppelgänger toxins. We assembled venom gland transcriptomes from 45 phylogenetically diverse cone snails (see **Table S7**) as described for the RNAseq assembly of *C. textile* tissues described above. The identified neuropeptide precursors were used as queries in tBLASTn searches against the assembled venom gland transcriptomes using an e-value of 1e-5.

The identified doppelgänger toxins in *C. textile, C. ventricosus,* and *C. canariensis* were mapped to their genomic location using a nucleotide sequence homology search (BLASTn).

### Mass spectrometry

Publicly available proteomics data (PRIDE PXD038993) was analyzed using Xcalibur software (Thermo Scientific) and Byonic (Protein Metrics) to search for new doppelgänger peptides.

### *Conus* gene trees

We used a combination of published species trees from ^11,68,69^ for species tree in Figure 3.

Individual toxin gene families were calculated similarly. The sequences were aligned using mafft v.7.526^70^ and the trees was constructed using IQ-TREE v. 2.2.2.3^71^ with UF bootstrap calculation for 1000 replicates^72^. The best model according to the Bayesian Information Criterion was calculated to be GTR+F+I+R4 with ModelFinder^73^. The consensus tree was visualized using FigTree. The determined models of evolution can be found in the relevant supplementary figures.

### Gene structure analyses

We mapped assembled transcripts to the *C. textile* (or other) genomes using exonerate v. 2.76.2^74^ using both the p2g and e3g methods. The intron phases and positions were determined by this output file. In all cases, an alternative assembly (GCA_049309315.1) was compared. For the

Prohormone-4, GPHb, and CCAP families, this assembly was used, as it provided better resolution.

For non-neuropeptide/-conotoxin genes, we calculated intron phases by pairing the GFF file using a custom Python script. The subset of the secreted proteome was determined using SignalP6^67^ on the Braker predicted proteome, and the intron phases of this subset were calculated independently.

Large gene loci alignments were performed using Mauve snapshot 2015-02-25^75^ using progressiveMauve with standard settings and the HOXD scoring matrix.

### Untranslated region comparisons

We used a custom Python script to extract the 3’ and 5’ untranslated regions, as well as the coding regions from neuropeptide and doppelgänger toxin transcripts. Using the individual gene trees as references, we matched the doppelgänger toxin transcript with the most closely associated neuropeptide transcript and calculated the percentage identity for each region. Only UTRs longer than 15 nucleotides were included in the calculations.

### Conotoxin 5’ UTR comparisons

We extracted all 5’ UTRs longer than 15 nucleotides from the annotated conotoxin transcripts of 45 cone snail species, as well as the identified doppelgänger transcripts. These nucleotide sequences were clustered using a sequence-based clustering, CLANS^76,77^, performing all-against-all blastn searches between the extracted 5’ UTRs. The clustering was run for 10,000 rounds, at which point the clustering had converged.

## Acknowledgments

The PacBio sequencing service was provided by the Norwegian Sequencing Centre (www.sequencing.uio.no), a national technology platform hosted by the University of Oslo and supported by the “Functional Genomics” and “Infrastructure” programs of the Research Council of Norway and the Southeastern Regional Health Authorities. Assembly was performed on resources provided by Sigma2—the National Infrastructure for High Performance Computing and Data Storage in Norway. The National Genomics Infrastructure at SciLifeLab Uppsala (Sweden) provided access to their PacBio Revio instrument for part of the sequencing. The support and resources from the Center for High Performance Computing at the University of Utah are gratefully acknowledged. The support and resources from the Mass Spectrometry and Proteomics Core at the University of Utah are gratefully acknowledged. Research reported in this publication utilized the High-Throughput Genomics and Cancer Bioinformatics Shared Resource at Huntsman Cancer Institute at the University of Utah and was supported by the National Cancer Institute of the National Institutes of Health under Award Number P30CA042014. The content is solely the responsibility of the authors and does not necessarily represent the official views of the NIH. We would like to thank Dylan Taylor for the videos showcasing *C. textile*.

## Author contributions

Conceptualization: TLK, NDY, BMO, EABU, HSH

Methodology: TLK, GF, SBS, PFS, MW, KC, ATK, AOL, AY, NDY, EABU

Investigation: TLK, NDY, EABU, HSH

Visualization: TLK

Funding acquisition: TLK, NDY, BMO, EABU, HSH

Project administration: HSH

Supervision: NDY, BMO, EABU, HSH

Writing – original draft: TLK

Writing – review & editing: All authors

## Data and materials availability

The *C. textile* assemblies generated by this study is achieved under National Center for Biotechnology Information (NCBI) GenBank (accession number pending). RNAseq data from *C. textile* and *C. cordigera* is deposited at National Center for Biotechnology Information (NCBI) GenBank. (accession numbers pending). All additional data are available in the main text or the supplementary materials.

