## Supplementary figures for "From Neuropeptides to Toxins: Illuminating the Origins of Venom Complexity in Cone Snails"

**The PDF file includes:**

Figs. S1 to S45

Tables S1 to S8

References

**Other Supplementary Materials for this manuscript include the following:**

Movies S1 to S4

Data S1 to S9

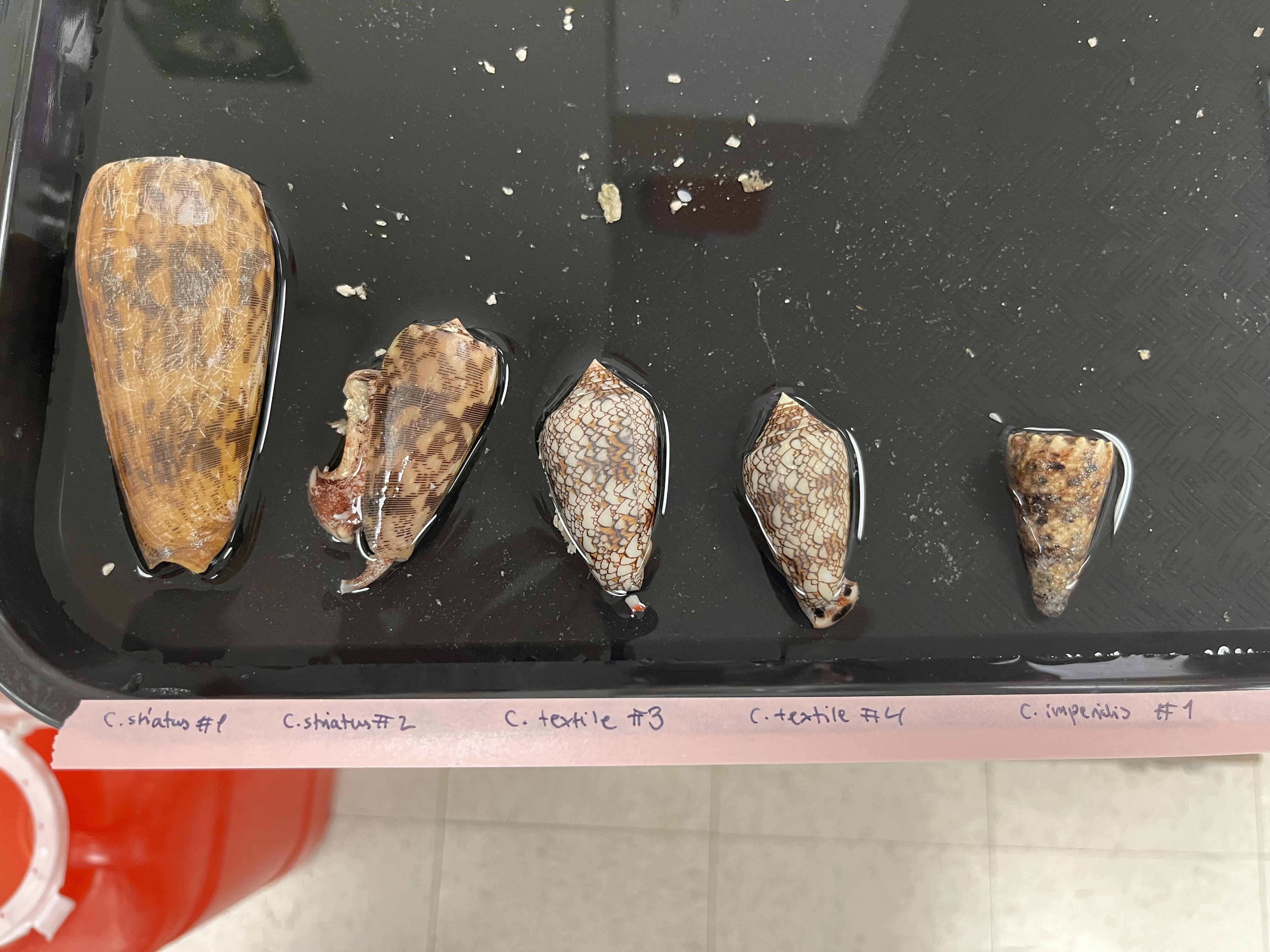

**Fig. S1. *Conus textile* specimen used for sequencing.**

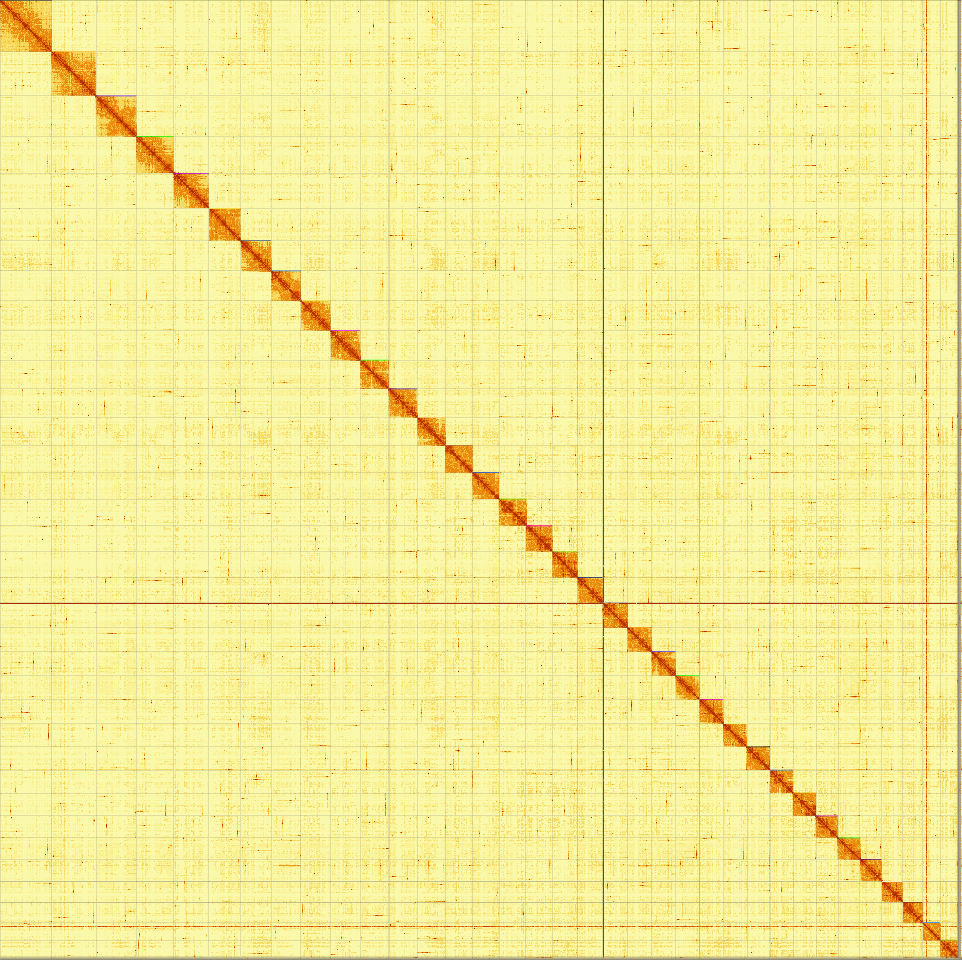

Fig. S2. Hi-C contact map. Hi-C contact map for the nuclear haplotypic genome of *Conus textile.* The contact map displays the chromosomal interactions within the genome at a resolution of 100 kb. The color corresponds with the frequency of interactions between genomic regions. Darker colors indicate a higher frequency of interactions

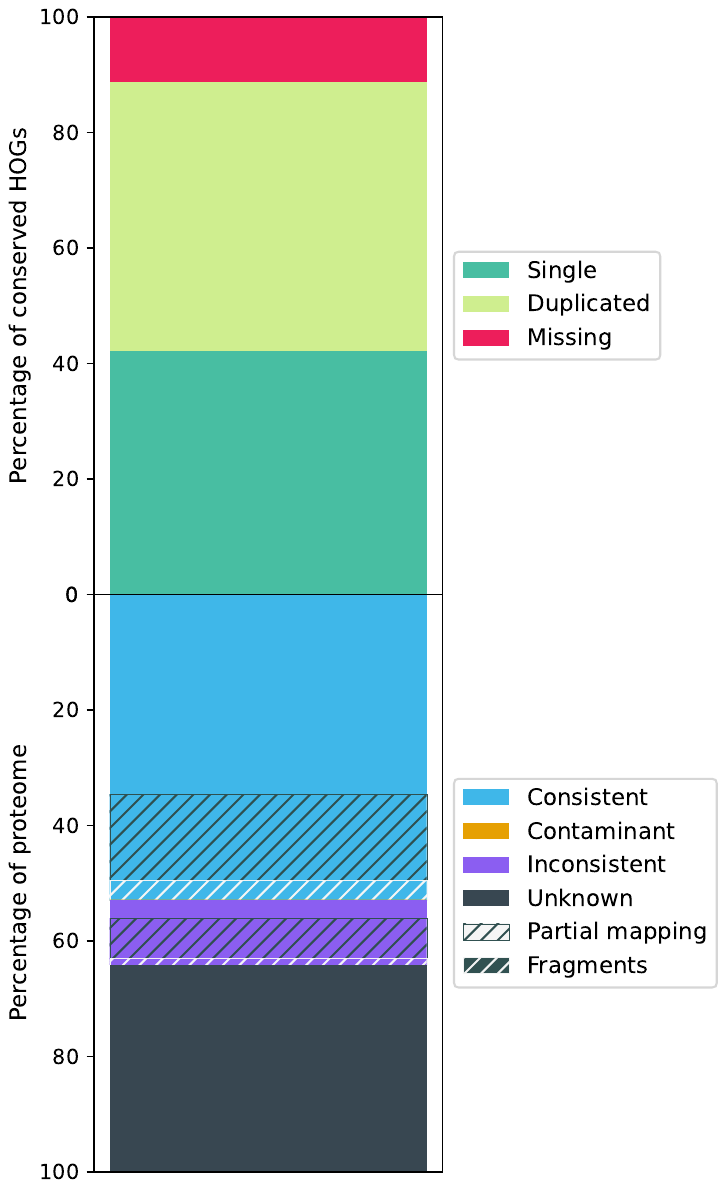

Fig. S3. OMArk completeness analysis. Schematic overview of the OMArk analysis. The top of the stacked bar represents the completeness assessment: Single copy (42.14%), Duplicity (46.61%), and Missing genes (11.25%). The lower part represents the consistency assessment: Consistent (52.85%), Inconsistent (11.29%), Unknown (35.86%), Contaminant (0 %).

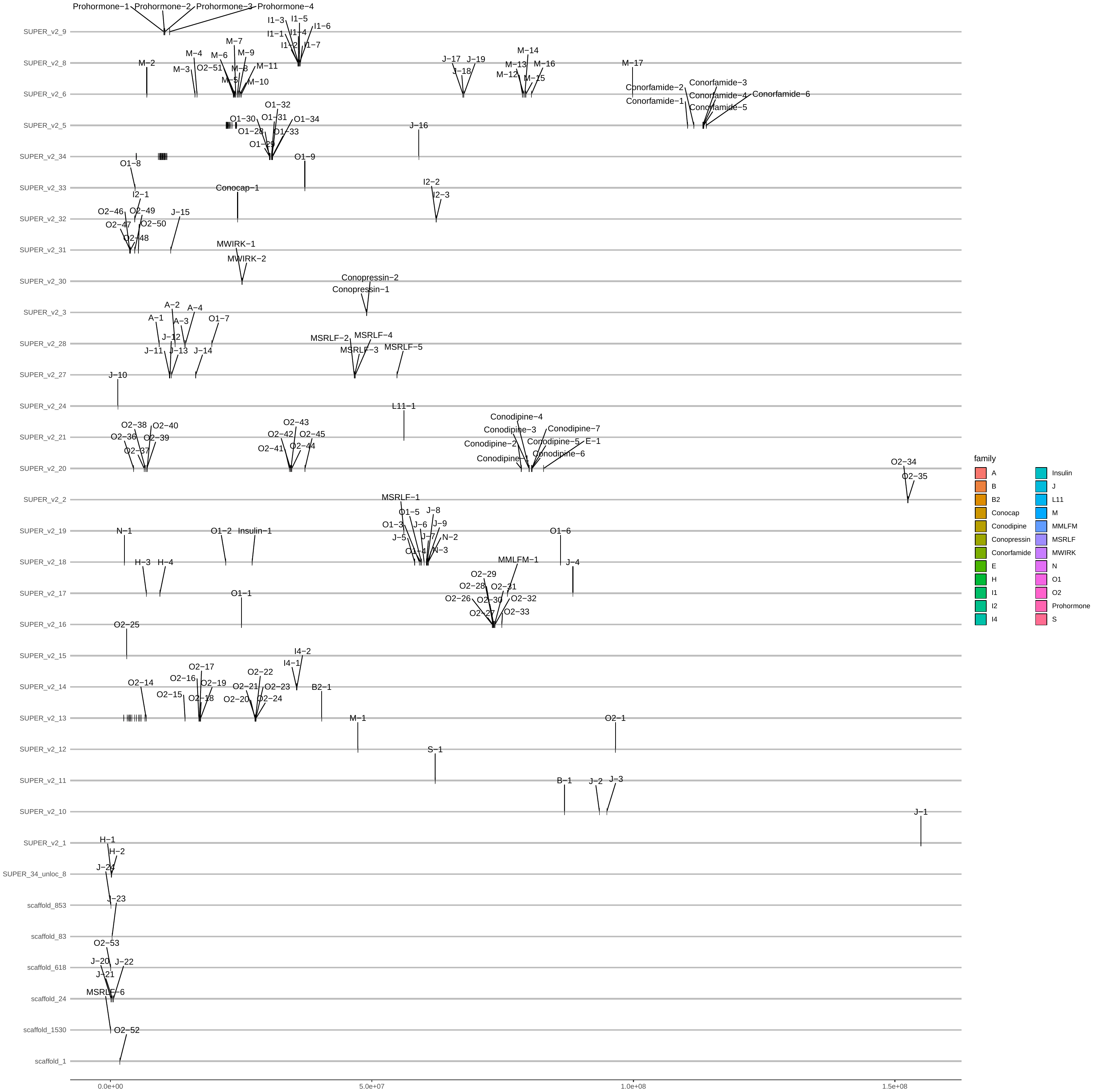

Fig. S4. Loci of full-length toxin genes in *C. textile* genome*.* ToxCodAn-genome annotations of full-length toxin genes in the *C. textile genome.* The toxins are annotated according to conotoxin gene superfamilies.

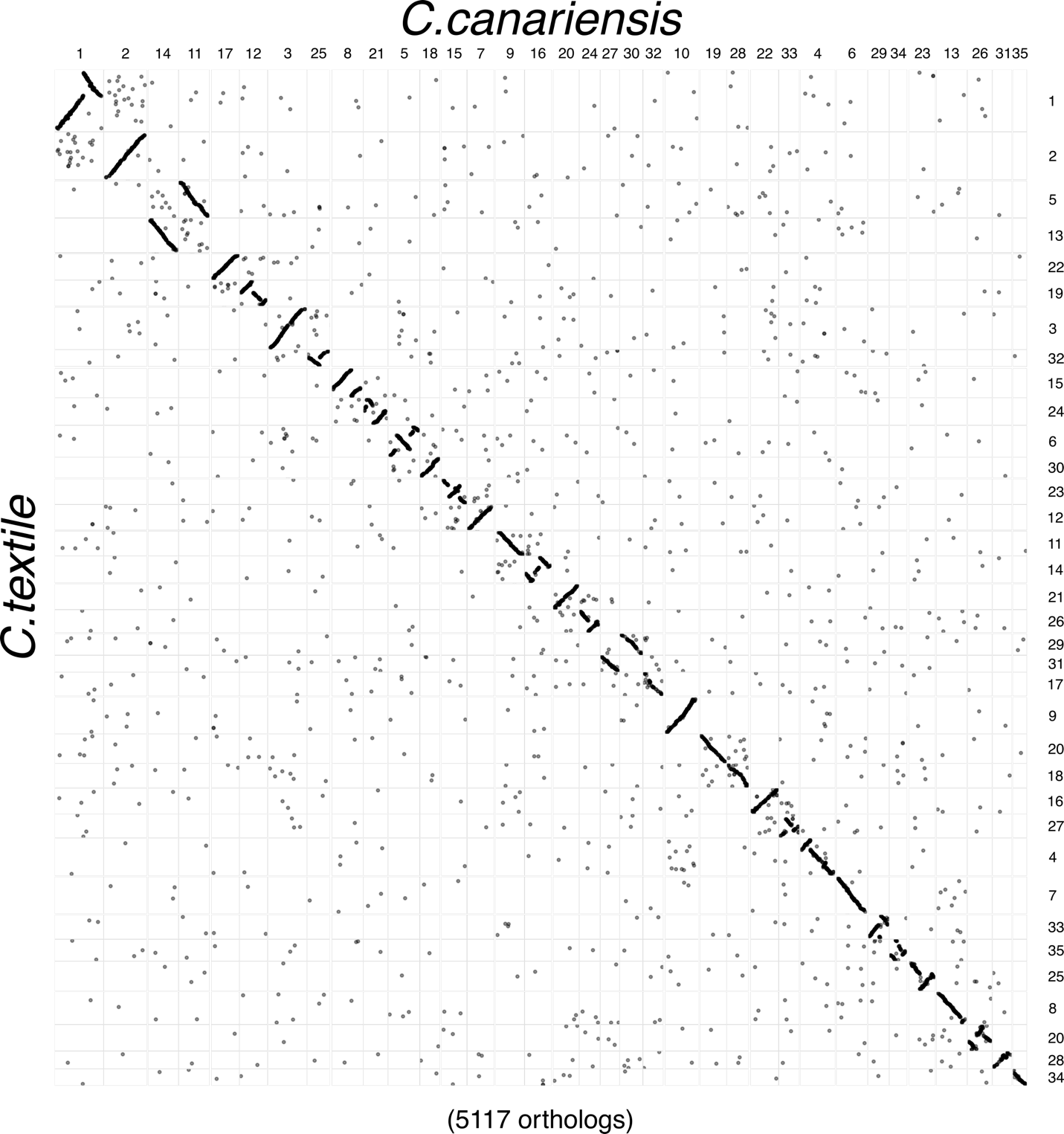

Fig. S5. Oxford dot-plot between *Conus canariensis* and *Conus textile.* The dot-plot shows the ordinal location of orthologous genes in *C. canariensis* and *C. textile* determined by reciprocal best blast hits. The colored dots represent genes with consistently statistically significant conserved chromosomal synteny; genes of variable synteny are shown in grey.

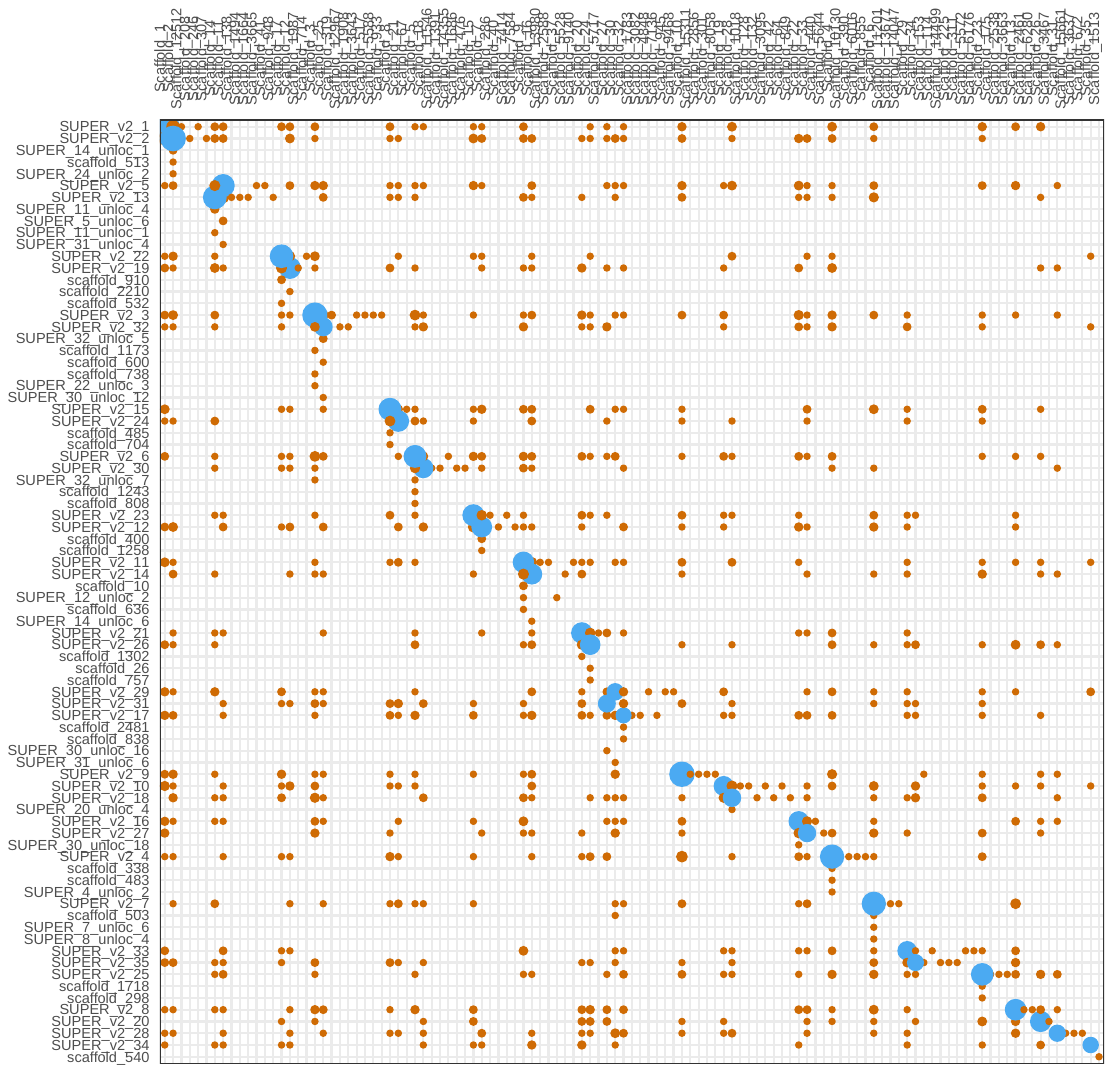

Fig. S6. Significant linkage groups between *Conus canariensis* and *Conus textile.* Significant chromosome-chromosome synteny calculated by Fisher’s exact test as described in^1^. Blue circles represent significant macrosynteny.

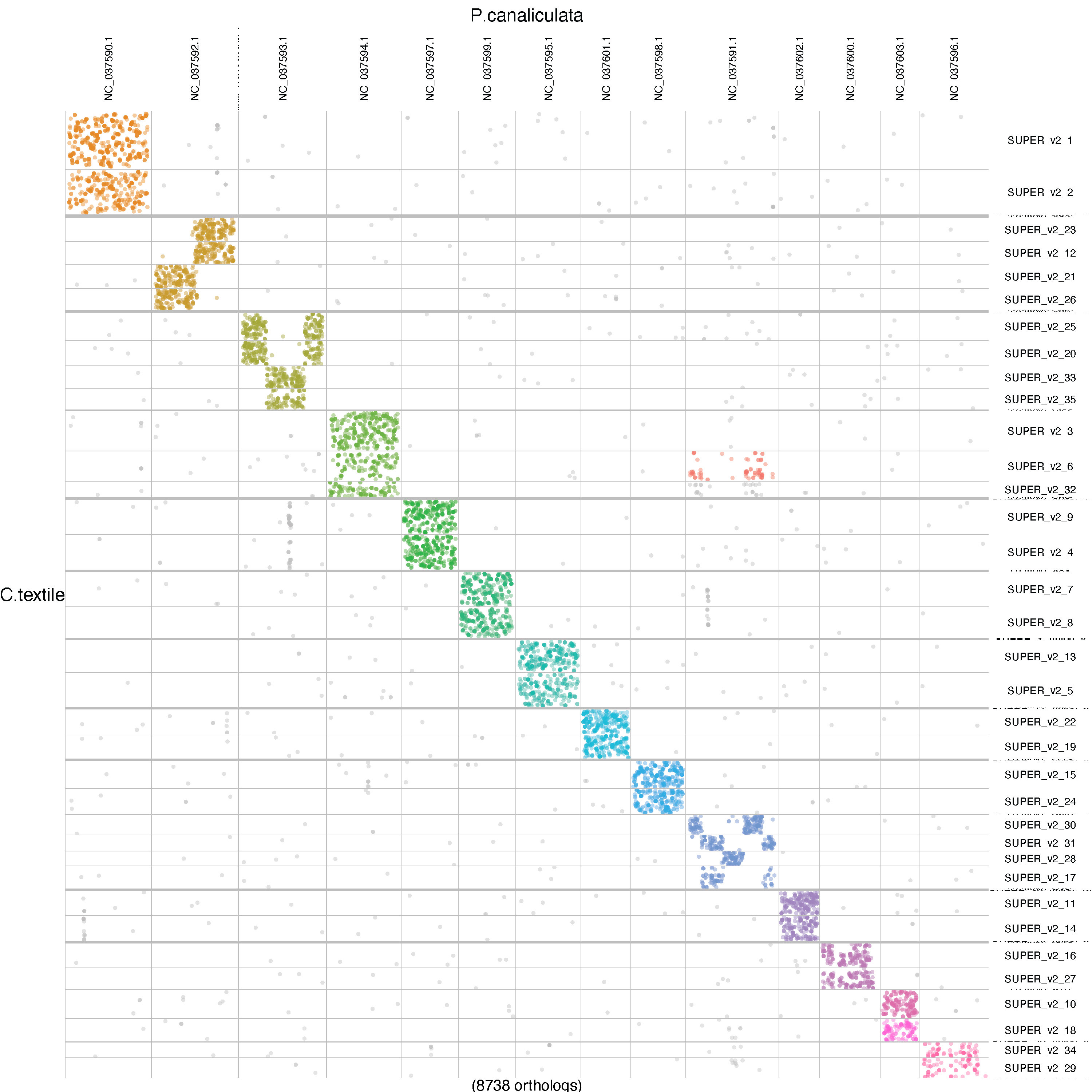

Fig. S7. Oxford dot-plot between *Pomacea canaliculata* and *Conus textile.* The dot-plot shows the ordinal location of orthologous genes in *P. canaliculata* and *C. textile* determined by reciprocal best blast hits. The colored dots represent genes with consistently statistically significant conserved chromosomal synteny; genes of variable synteny are shown in grey.

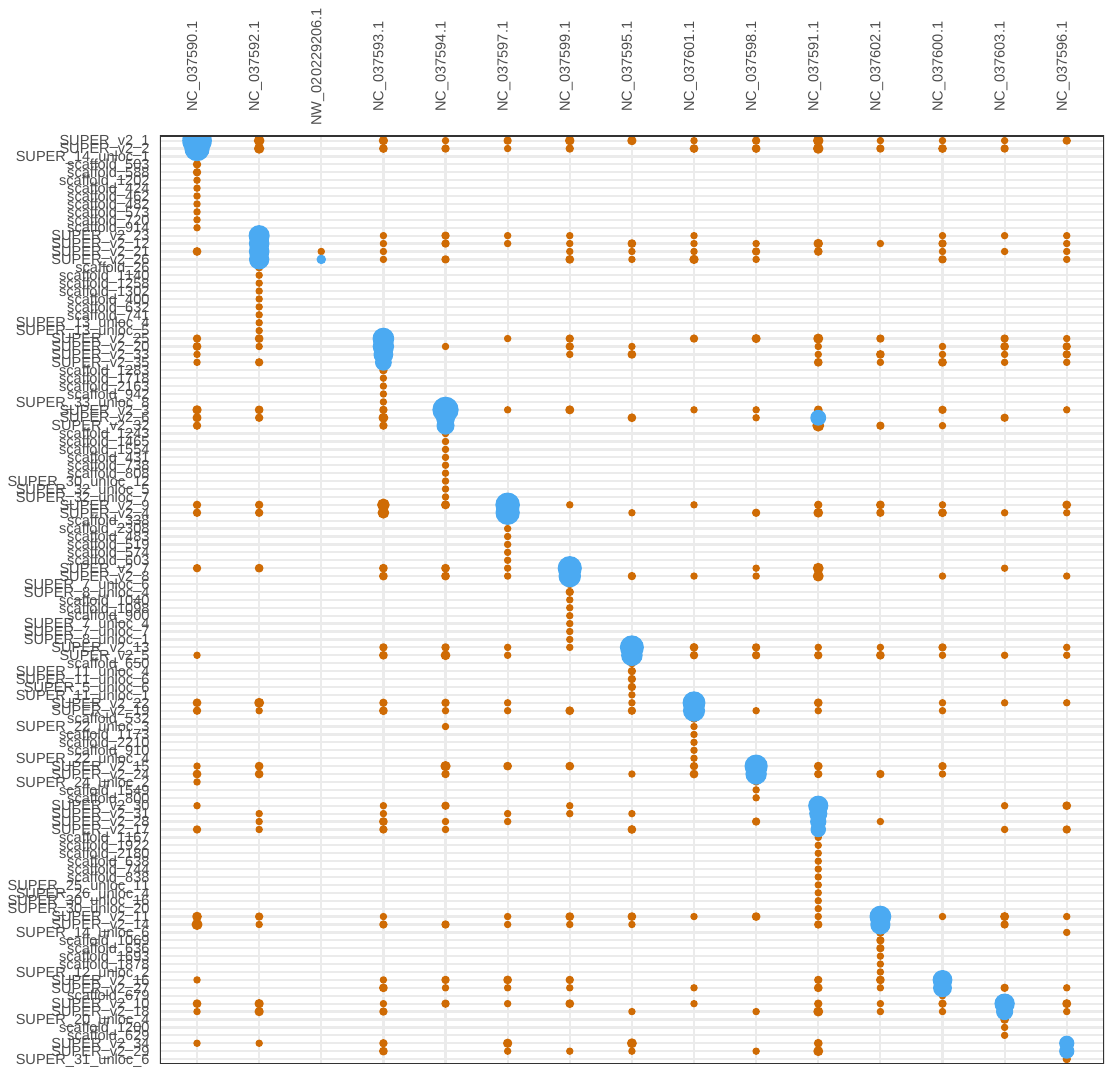

Fig. S8. Significant linkage groups between *Pomacea canaliculata* and *Conus textile.* Significant chromosome-chromosome synteny calculated by Fisher’s exact test as described in^1^. Blue circles represent significant macrosynteny.

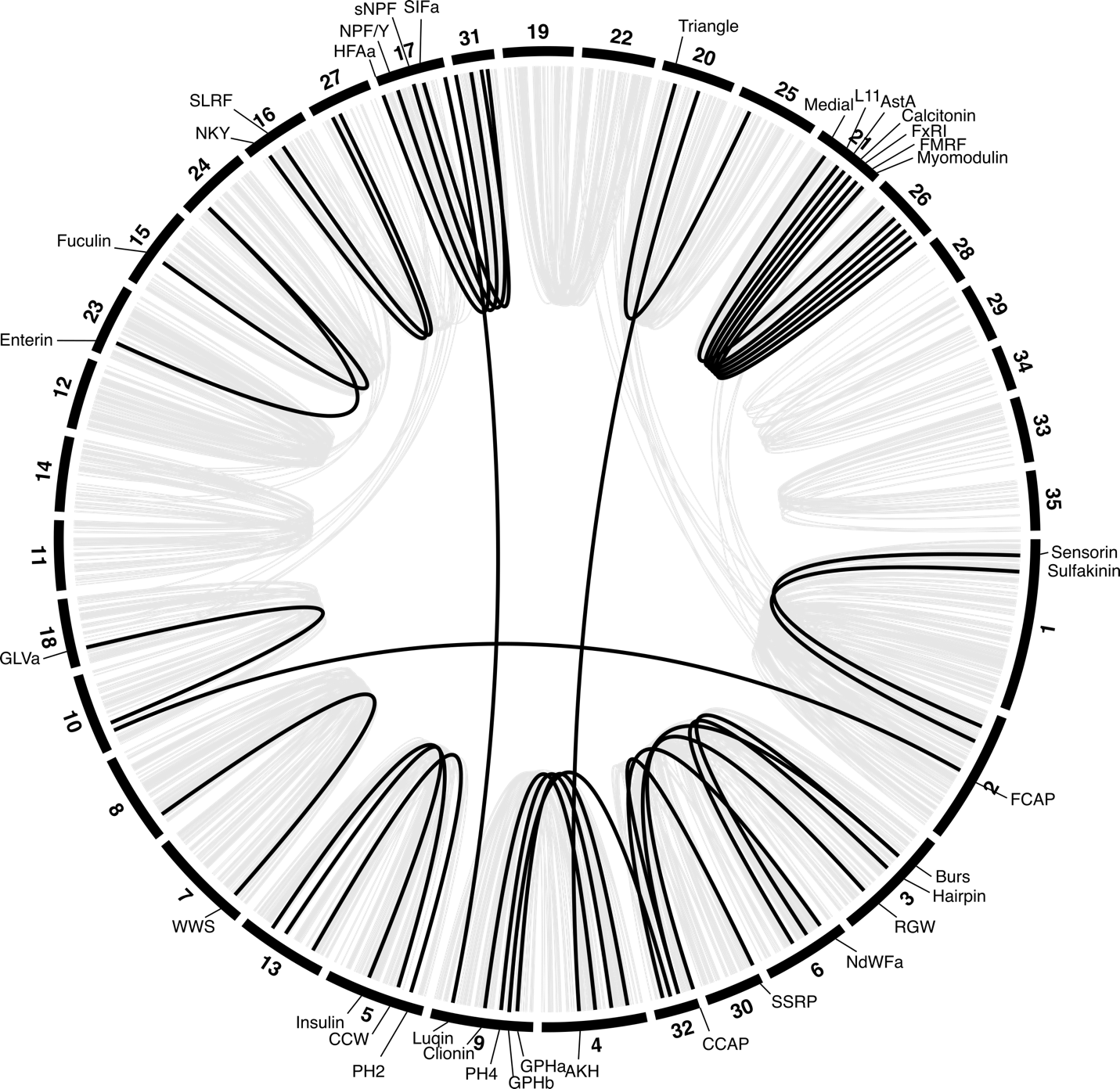

Fig. S9. Neuropeptide ohnologs are typically located on homologous chromosomes in *C. textile*. The figure matches Fig2B but shows additional multi-gene neuropeptide families and their location in the *C. textile* genomes. The consistent location on homologous chromosomes supports their designation as ohnologs. Some neuropeptide genes identified transcriptomically were only found as partial genes or missing.

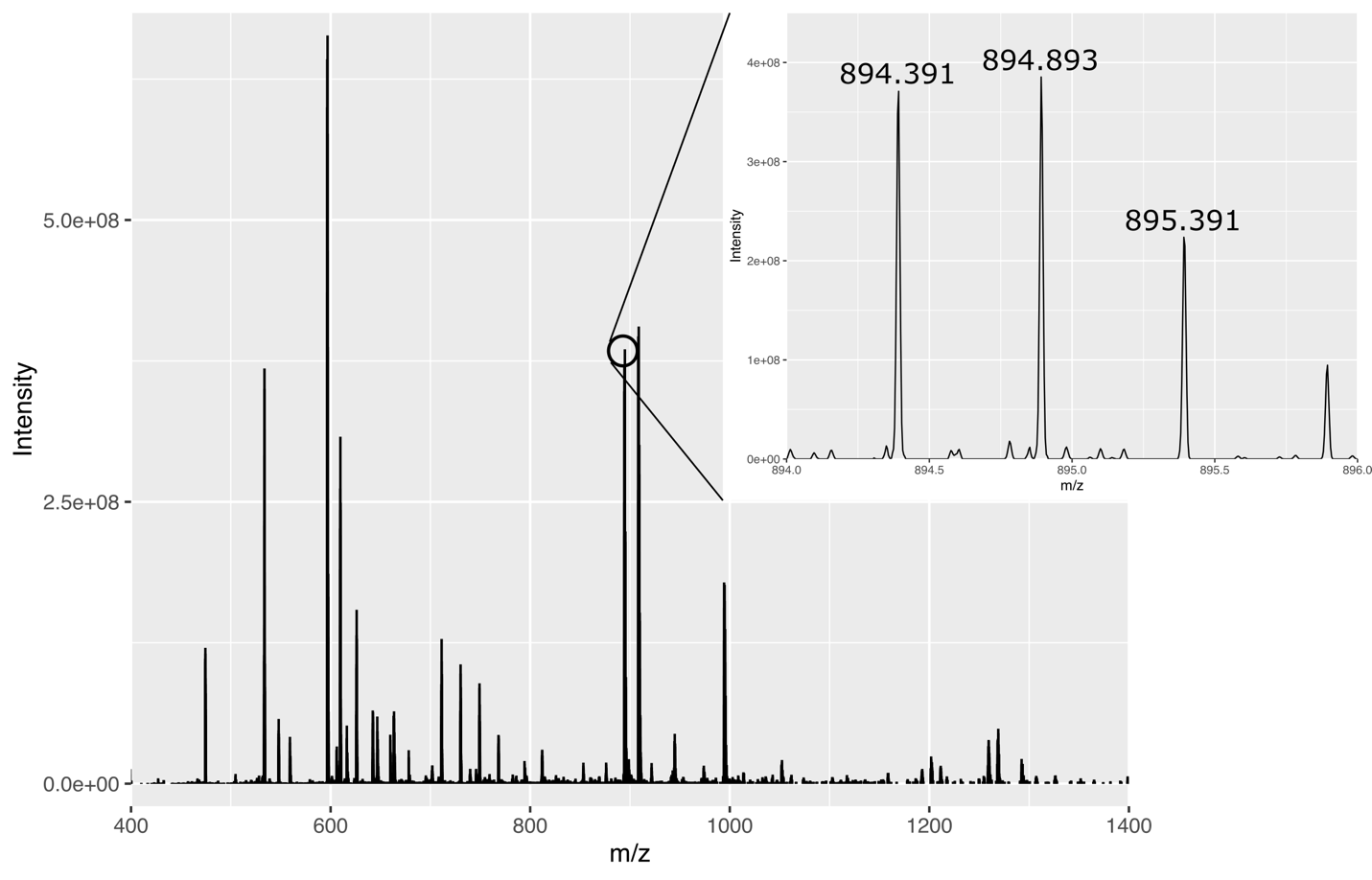

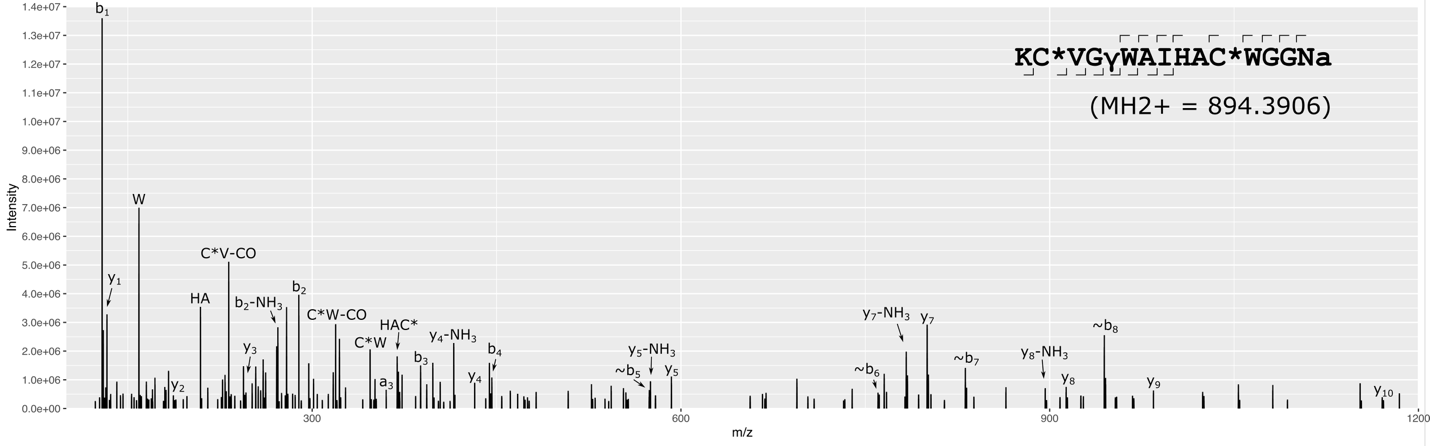
**Fig. S10. Mass spectrometric identification of GGNamide.** MS of precursor ion (upper) and MS/MS of fragment ion (lower) spectra of GGNamide doppelgänger toxin in *C. textile*.

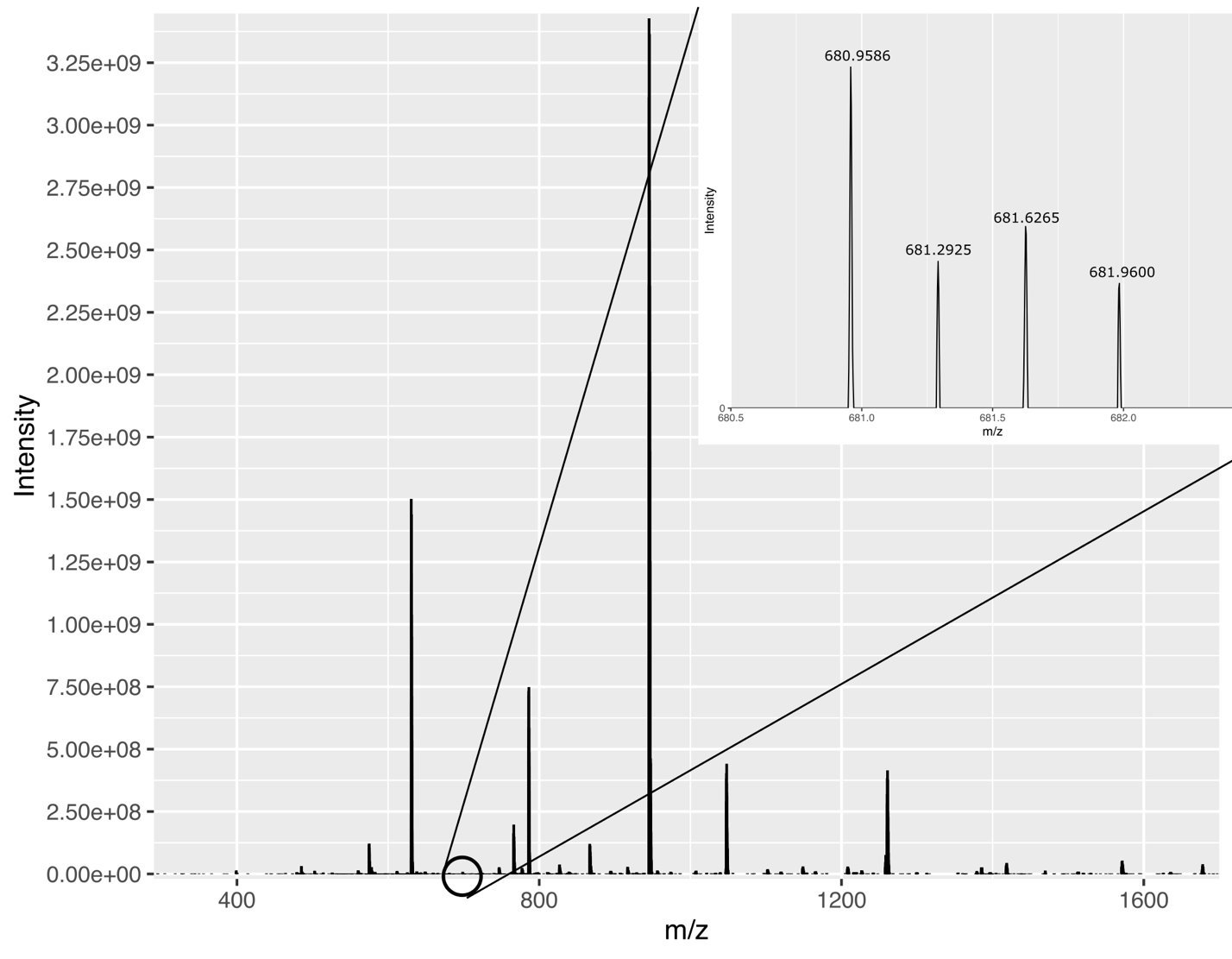

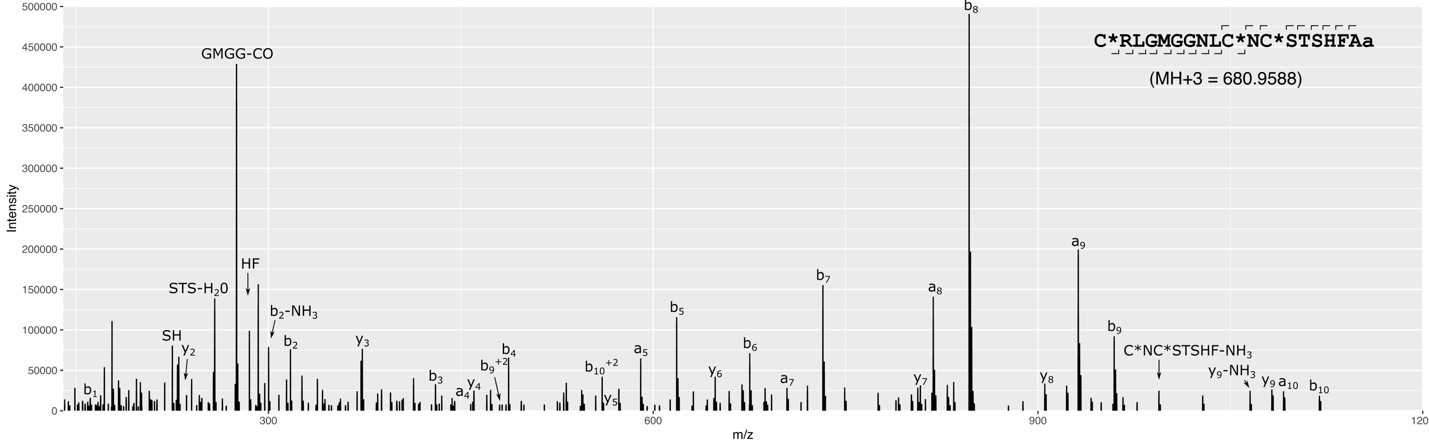
**Fig. S11. Mass spectrometric identification of HFAamide.** MS of precursor ion (upper) and MS/MS of fragment ion (lower) spectra of HFAamide doppelgänger toxin in *C. textile*.

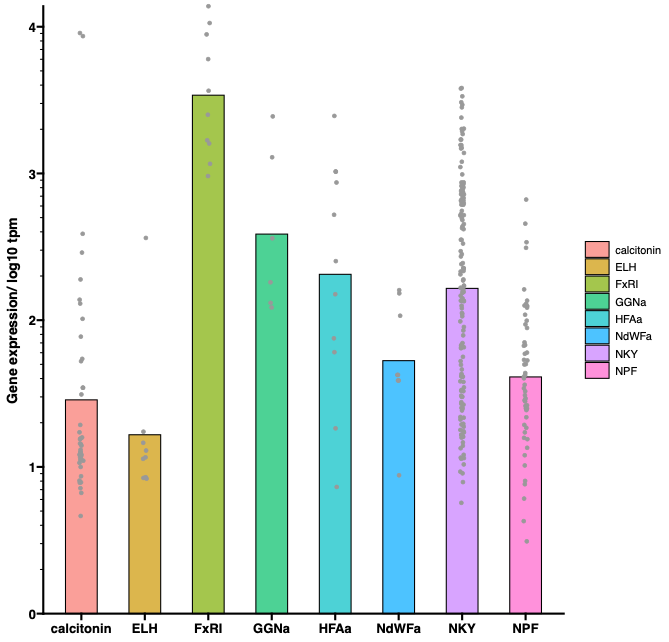
**Fig. S12. Expression of new doppelgänger toxin families.** Expression levels, measured by transcripts per million (tpm), are shown for identified transcripts belonging to the different new doppelgänger toxin families. The NPF-doppelgänger genes include only transcripts belonging to the large lineage of doppelgängers found in *Conus miliaris* and its sister species.

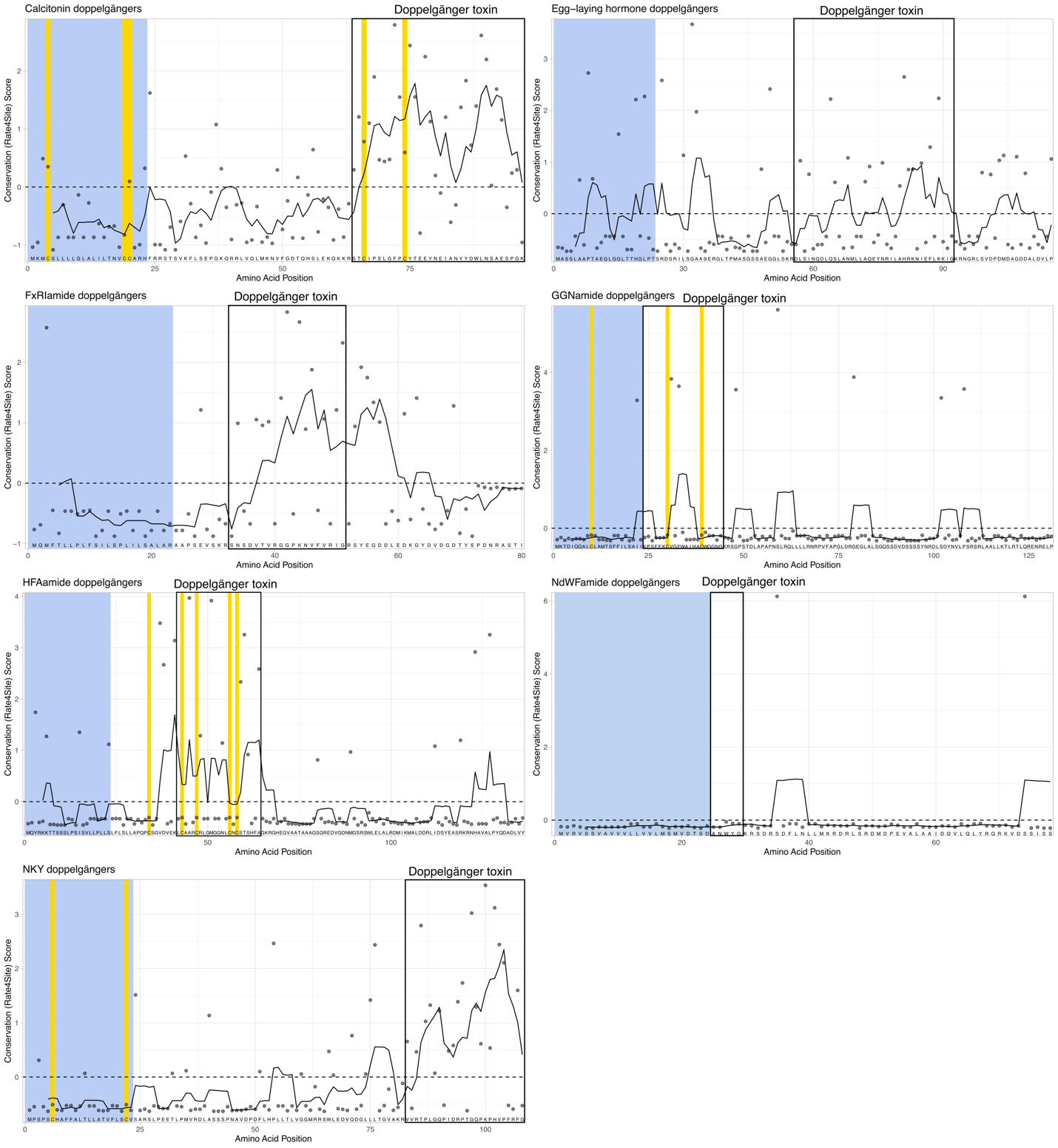

**Fig. S13. Evolutionary trace analyses of new doppelgänger toxin families.** Four of the newly identified doppelgänger toxins (Calcitonin, FxRIamide, HFAamide, and NKY) exhibit the characteristic pattern of more rapidly evolving toxin-encoding regions compared to their precursor sequences. As discussed in the main manuscript, the divergence patterns of doppelgänger toxin conservation generally correspond to cases of doppelgänger toxins have evolved through alternative splicing and represent more recently evolved toxins. The predicted mature peptides are boxed.

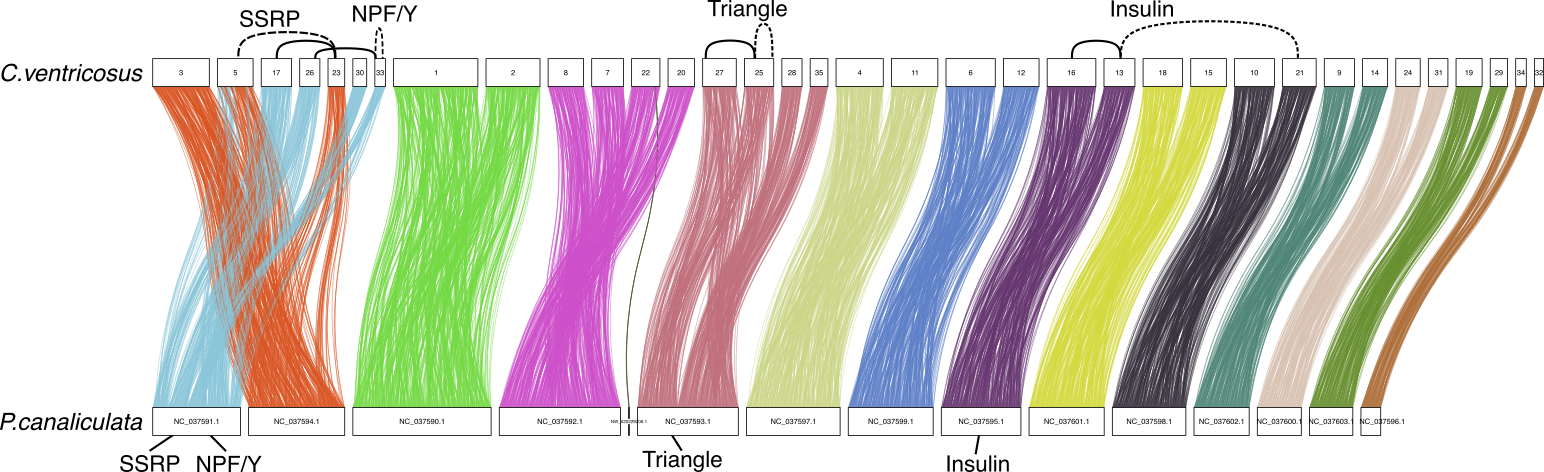

Fig. S14. Doppelgänger toxins and corresponding neuropeptide genes in *C. ventricosus*. Full lines mark the locations of neuropeptide ohnologs and dashed lines show the location of the corresponding doppelgänger toxins. The orthologous neuropeptide genes in *P. canaliculata* are shown.

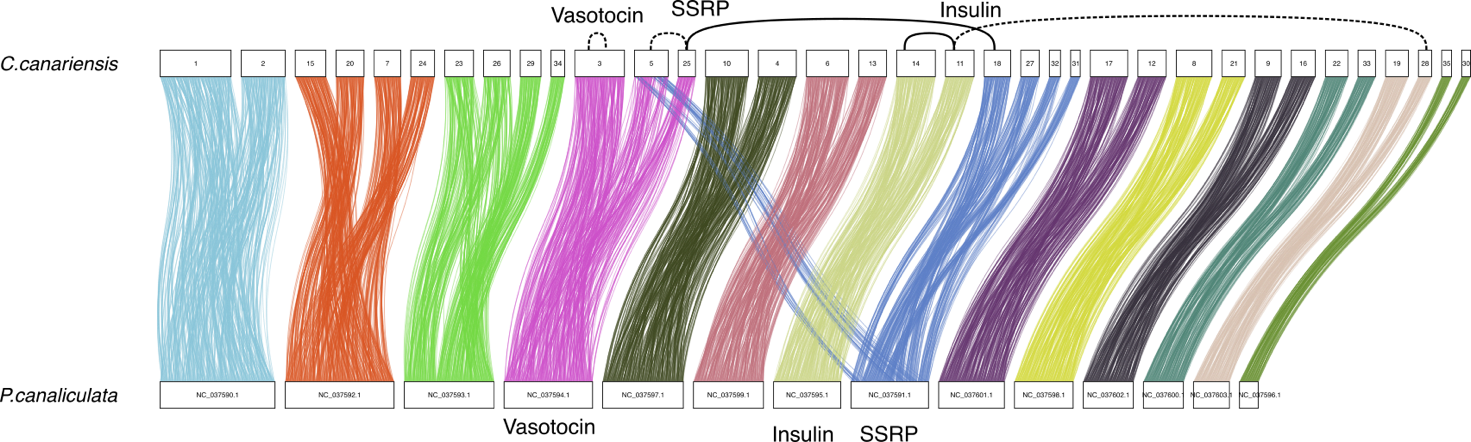

Fig. S15. Doppelgänger toxins and corresponding neuropeptide genes in *C. canariensis* genomes. Full lines mark the locations of neuropeptide ohnologs and dashed lines show the location of the corresponding doppelgänger toxins. The orthologous neuropeptide genes in *P. canaliculata* are shown.

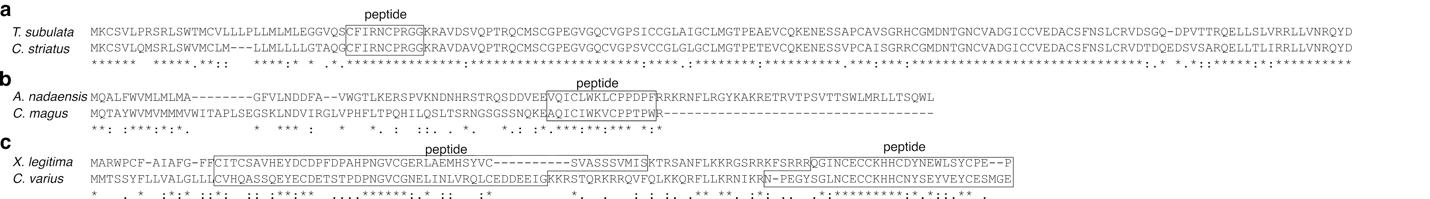
**Fig. S16. Presence of conopressin, consomatin, and con-insulins in non-*Conus* Conoidia. (A)** Sequence alignment of *Terebra subulata* conopressin identified in the venom gland transcriptome (SRR2060989) with *Conus striatus* conopressin. (B) Sequence alignment of *Annulaturris nadaensis* consomatin identified in the venom gland transcriptome (SRR26897527) with *Conus magus* consomatin. (C) Sequence alignment of *Xenuroturrus legitima* conopressin identified in the venom gland transcriptome (SRR26897521) with *Conus varius* con-insulin.

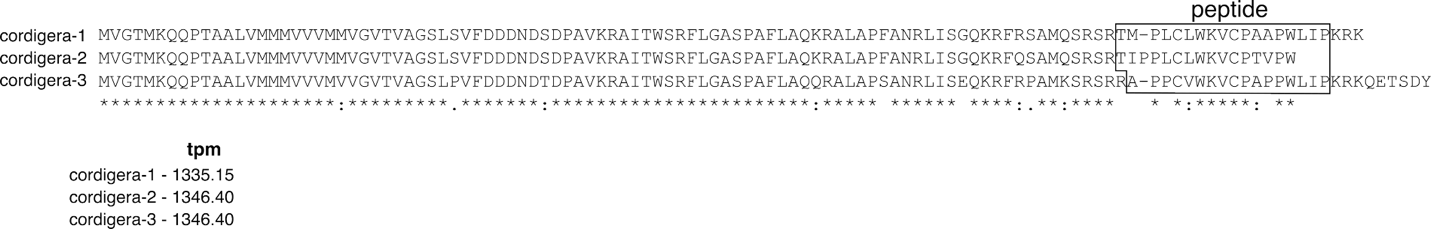

**Fig. S17. *Conus cordigera* SSRP doppelgänger toxins.** Sequence alignment of the three unique *C. cordigera* SSRP doppelgänger toxins, highlighting the mature peptide/toxin region where the primary sequence occurs. Expression levels (TPM) for each transcript are indicated below the alignment.

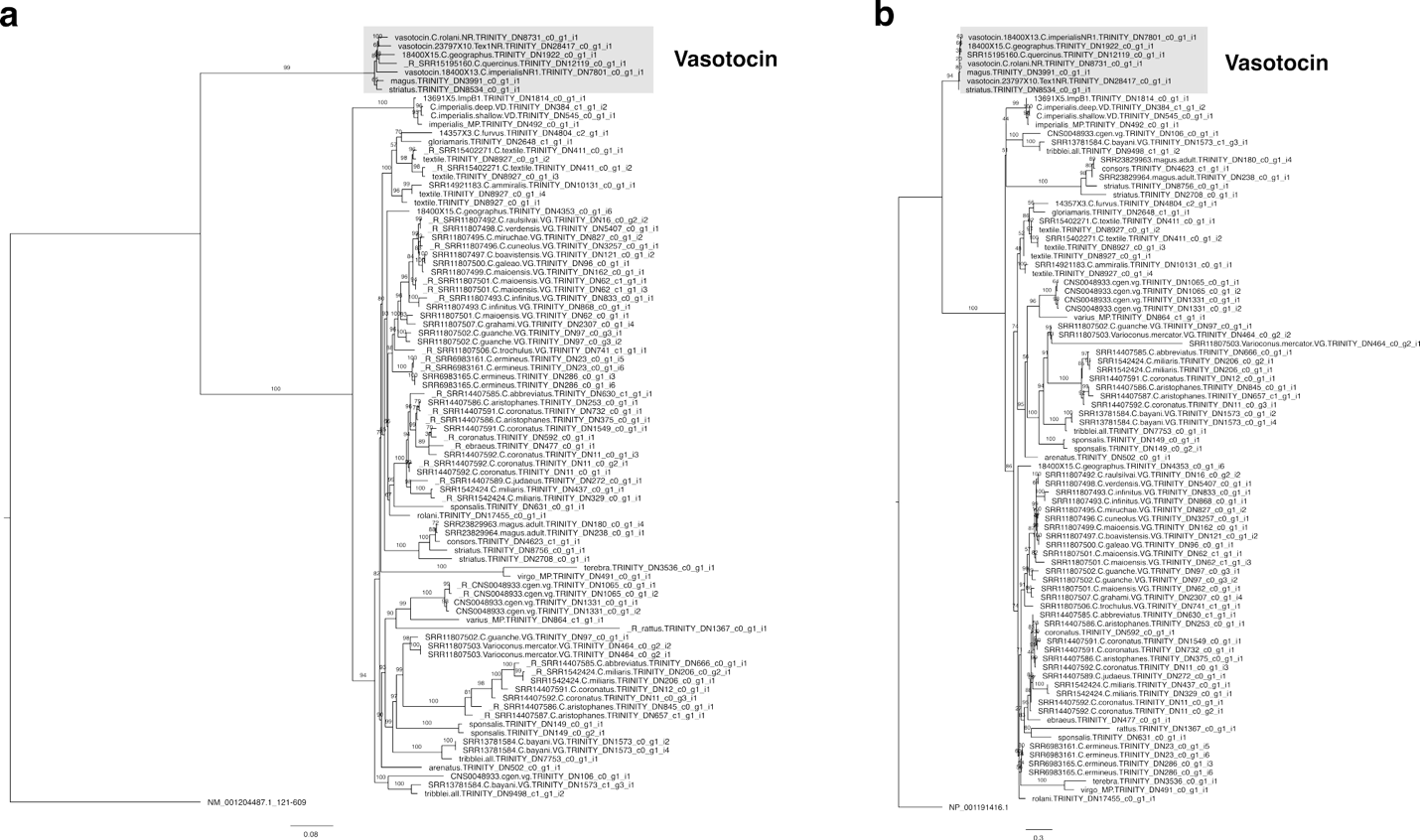

**Fig. S18. Vasotocin doppelgänger evolution. (A)** Maximum likelihood tree reconstruction of nucleotide alignment of vastocin neuropeptides and conopressin doppelgänger toxins. Using ModelSelector, the HKY+F+G4 model of evolution was chosen based on the Bayesian Information Criterion. The tree was outgrouped with *Aplysia californica* vasotocin. (B) Maximum likelihood tree reconstruction of protein alignment of vastocin neuropeptides and conopressin doppelgänger toxins. Using ModelSelector, the JTTDCMut+G4 model of evolution was chosen based on the Bayesian Information Criterion. The tree was outgrouped with *A. californica* vasotocin.

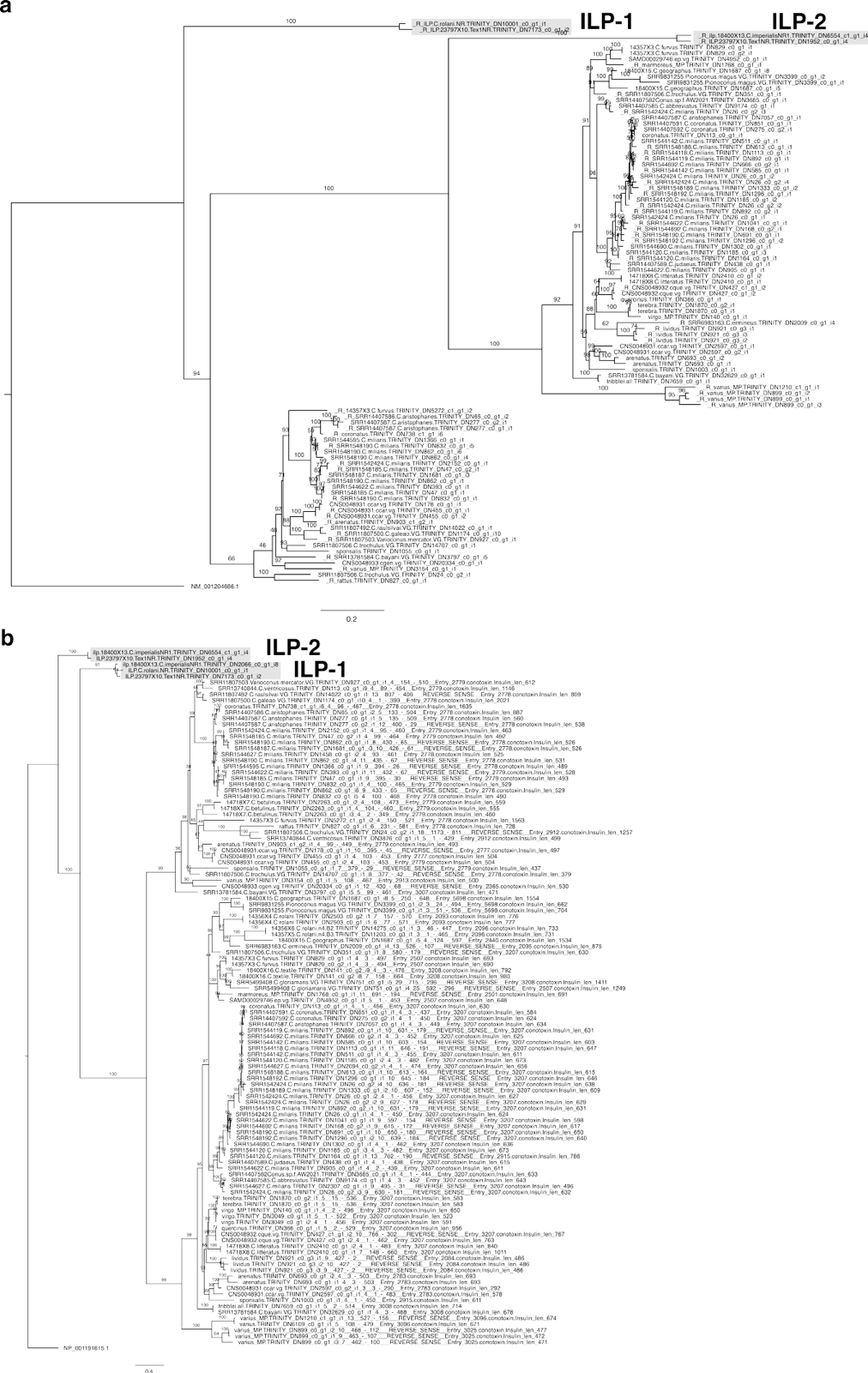

**Fig. S19. Insulin doppelgänger evolution. (A)** Maximum likelihood tree reconstruction of nucleotide alignment of insulin neuropeptides and doppelgänger toxins. Using ModelSelector, the K2P+R2 model of evolution was chosen based on the Bayesian Information Criterion. The tree was outgrouped with *A. californica* insulin. (B) Maximum likelihood tree reconstruction of protein alignment of insulin neuropeptides and doppelgänger toxins. Using ModelSelector, the JTT+G4 model of evolution was chosen based on the Bayesian Information Criterion. The tree was outgrouped with *A. californica* insulin.

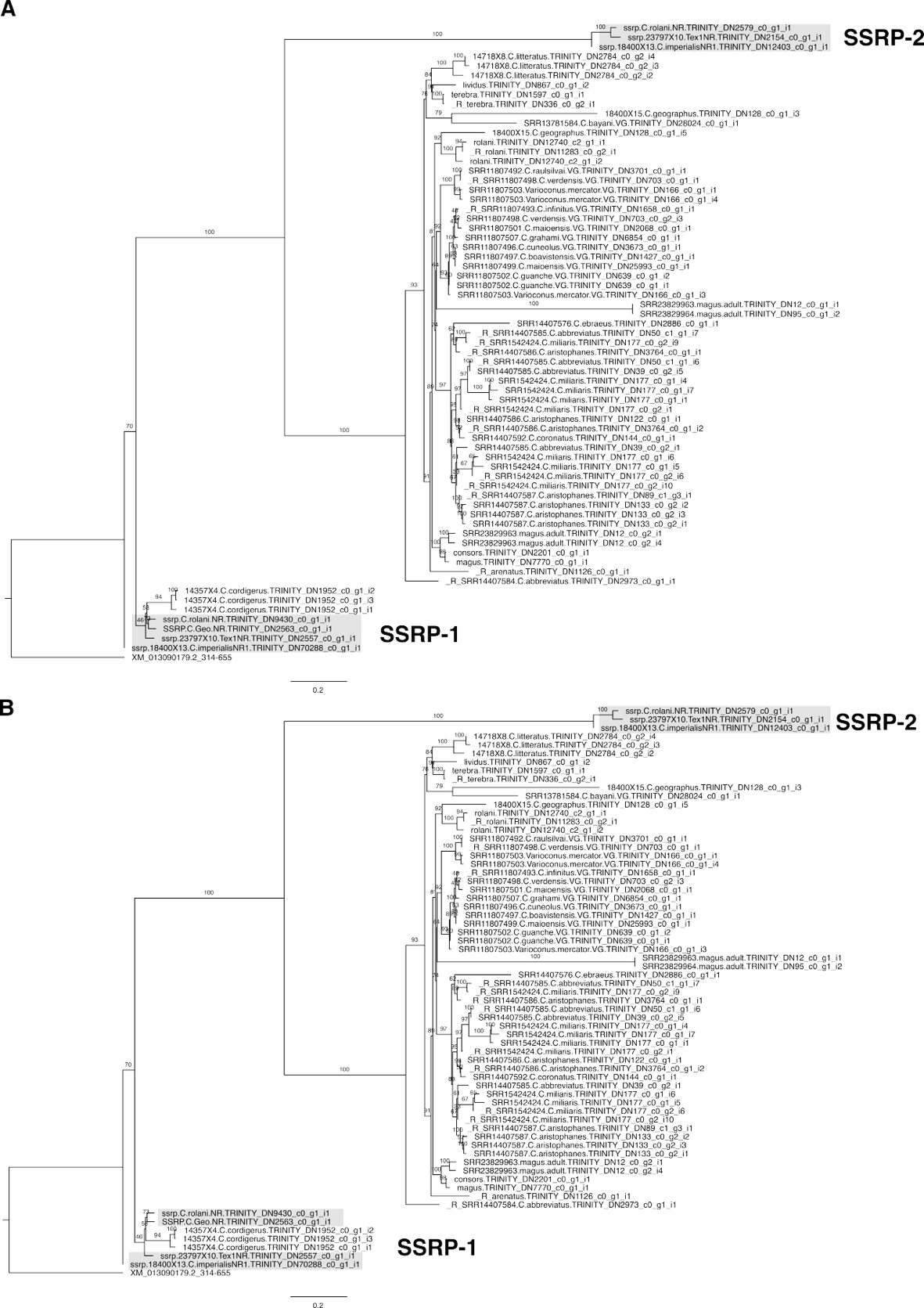

**Fig. S20. SSRP doppelgänger evolution. (A)** Maximum likelihood tree reconstruction of nucleotide alignment of SSRP neuropeptides and consomatin doppelgänger toxins. Using ModelSelector, the TN+F+R3 model of evolution was chosen based on the Bayesian Information Criterion. The tree was outgrouped with *A. californica* SSRP. (B) Maximum likelihood tree reconstruction of protein alignment of SSRP neuropeptides and consomatin doppelgänger toxins. Using ModelSelector, the Q.mammal+G4 model of evolution was chosen based on the Bayesian Information Criterion. The tree was outgrouped with *A. californica* SSRP.

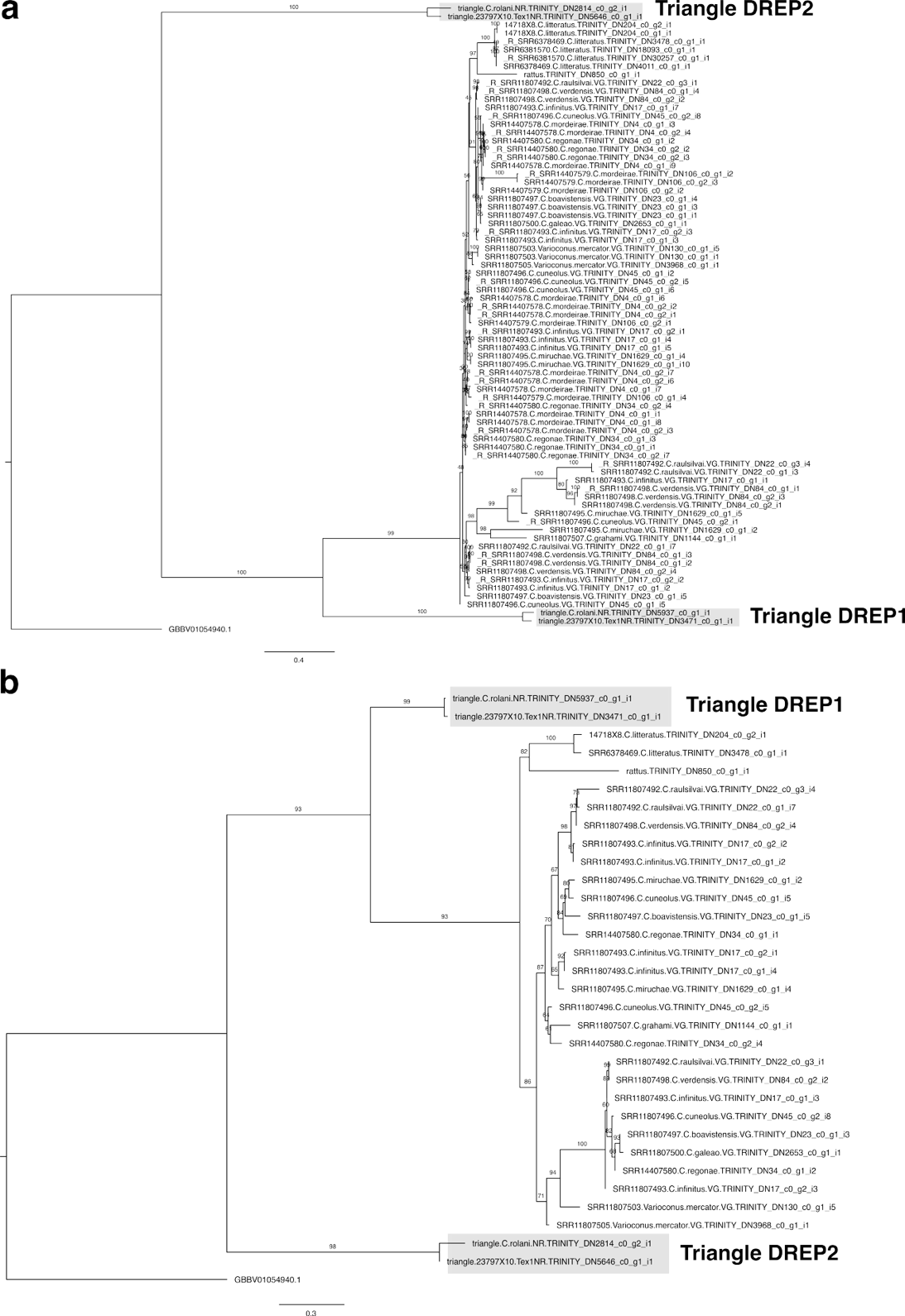

**Fig. S21. Triangle doppelgänger evolution. (A)** Maximum likelihood tree reconstruction of nucleotide alignment of Triangle neuropeptides and doppelgänger toxins. Using ModelSelector, the TPM2+I+R3 model of evolution was chosen based on the Bayesian Information Criterion. The tree was outgrouped with *A. californica* Triangle. (B) Maximum likelihood tree reconstruction of protein alignment of Triangle neuropeptides and doppelgänger toxins. Using ModelSelector, the JTTDCMut+I+R2 model of evolution was chosen based on the Bayesian Information Criterion. The tree was outgrouped with *A. californica* Triangle.

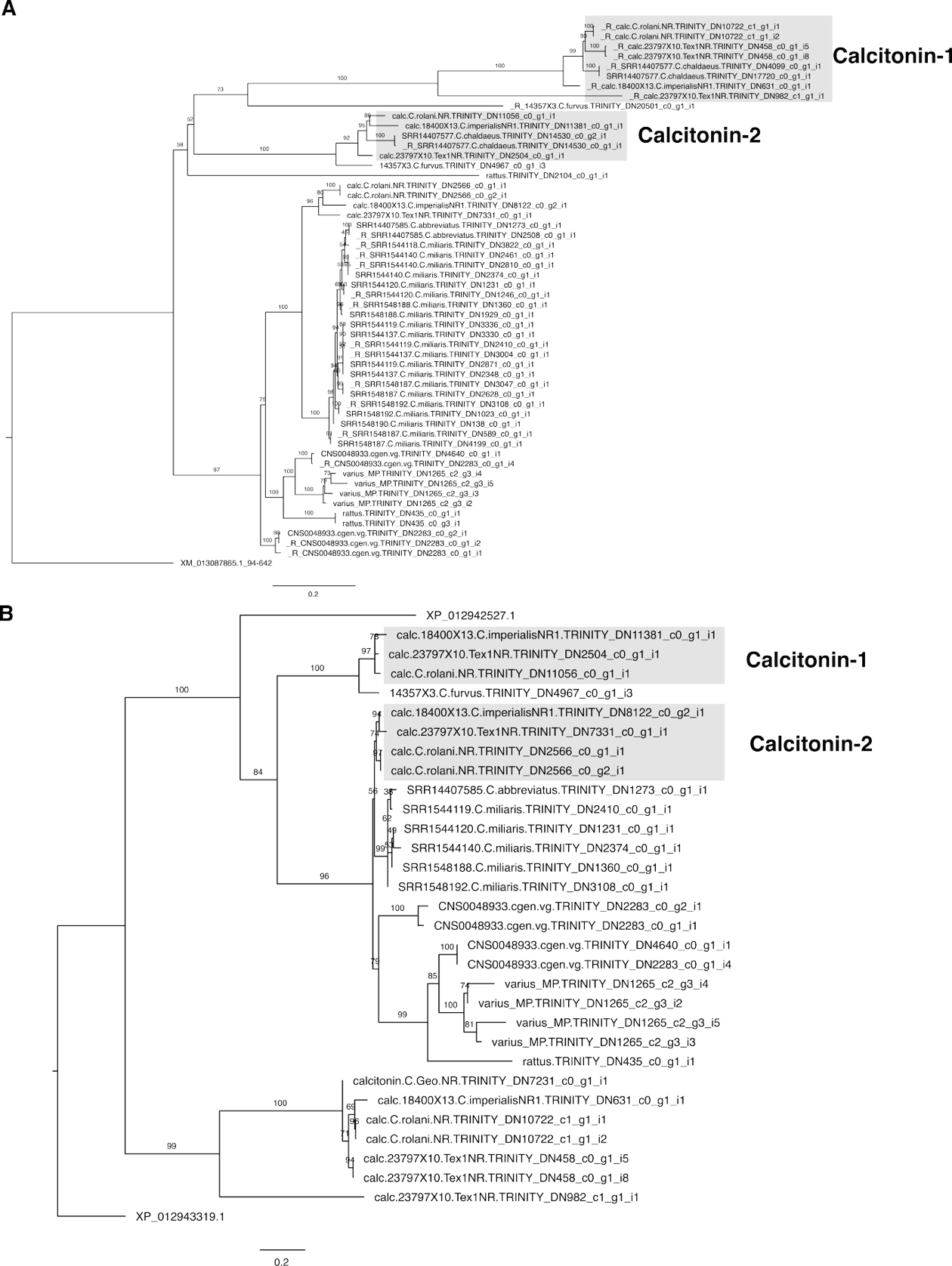

**Fig. S22. Calcitonin doppelgänger evolution. (A)** Maximum likelihood tree reconstruction of nucleotide alignment of Calcitonin neuropeptides and doppelgänger toxins. Using ModelSelector, the K2P+G4 model of evolution was chosen based on the Bayesian Information Criterion. The tree was outgrouped with *A. californica* Calcitonin. (B) Maximum likelihood tree reconstruction of protein alignment of calcitonin neuropeptides and conopressin doppelgänger toxins. Using ModelSelector, the VT+G4model of evolution was chosen based on the Bayesian Information Criterion. The tree was outgrouped with *A. californica* Calcitonin.

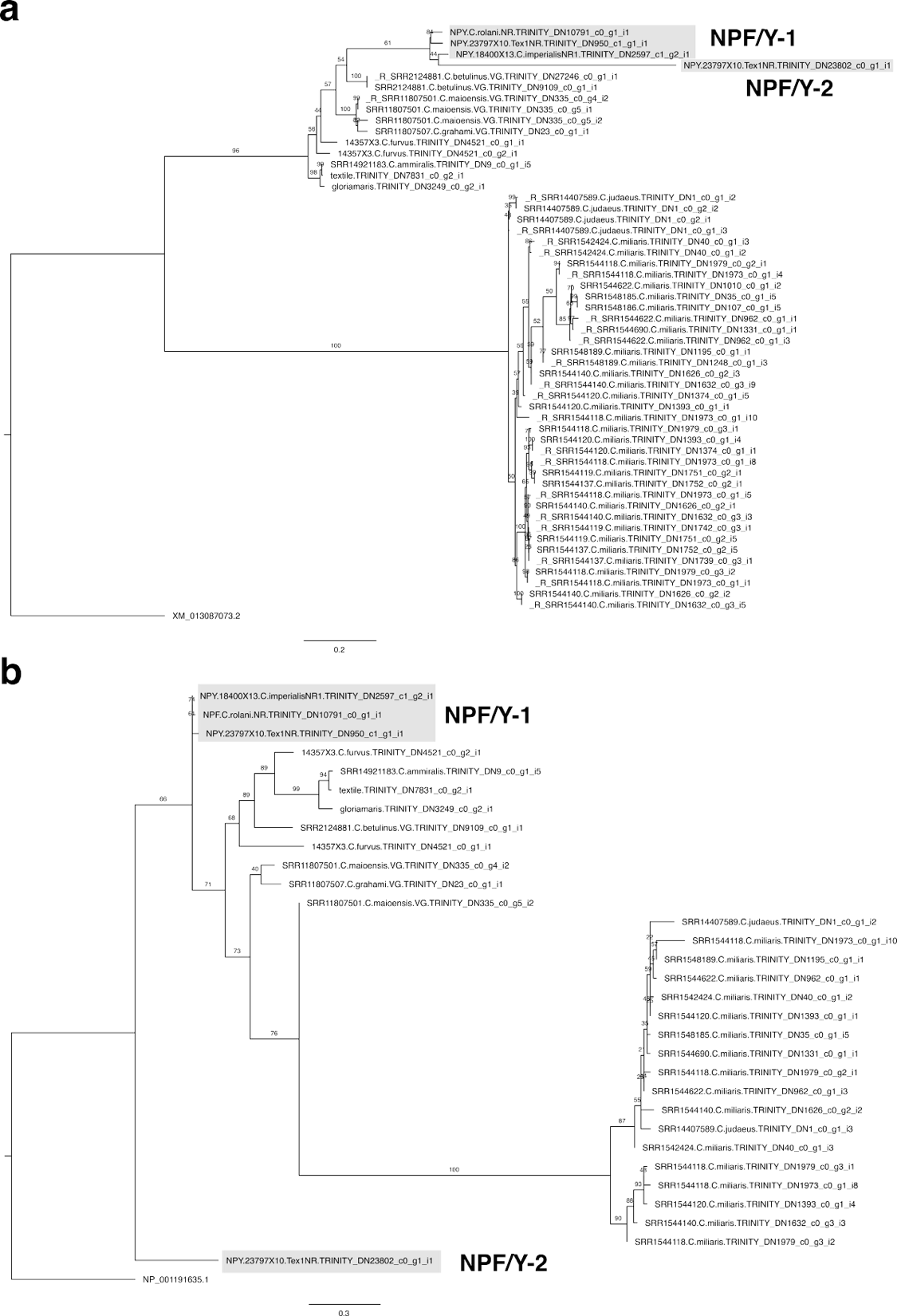

**Fig. S23. NPF/Y doppelgänger evolution. (A)** Maximum likelihood tree reconstruction of nucleotide alignment of NPF/Y neuropeptides and conoNPY doppelgänger toxins. Using ModelSelector, the HKY+F+R2 model of evolution was chosen based on the Bayesian Information Criterion. The tree was outgrouped with *A. californica* NPF. (B) Maximum likelihood tree reconstruction of protein alignment of NPF/Y neuropeptides and conoNPY doppelgänger toxins. Using ModelSelector, the WAG+R2 model of evolution was chosen based on the Bayesian Information Criterion. The tree was outgrouped with *A. californica* NPF.

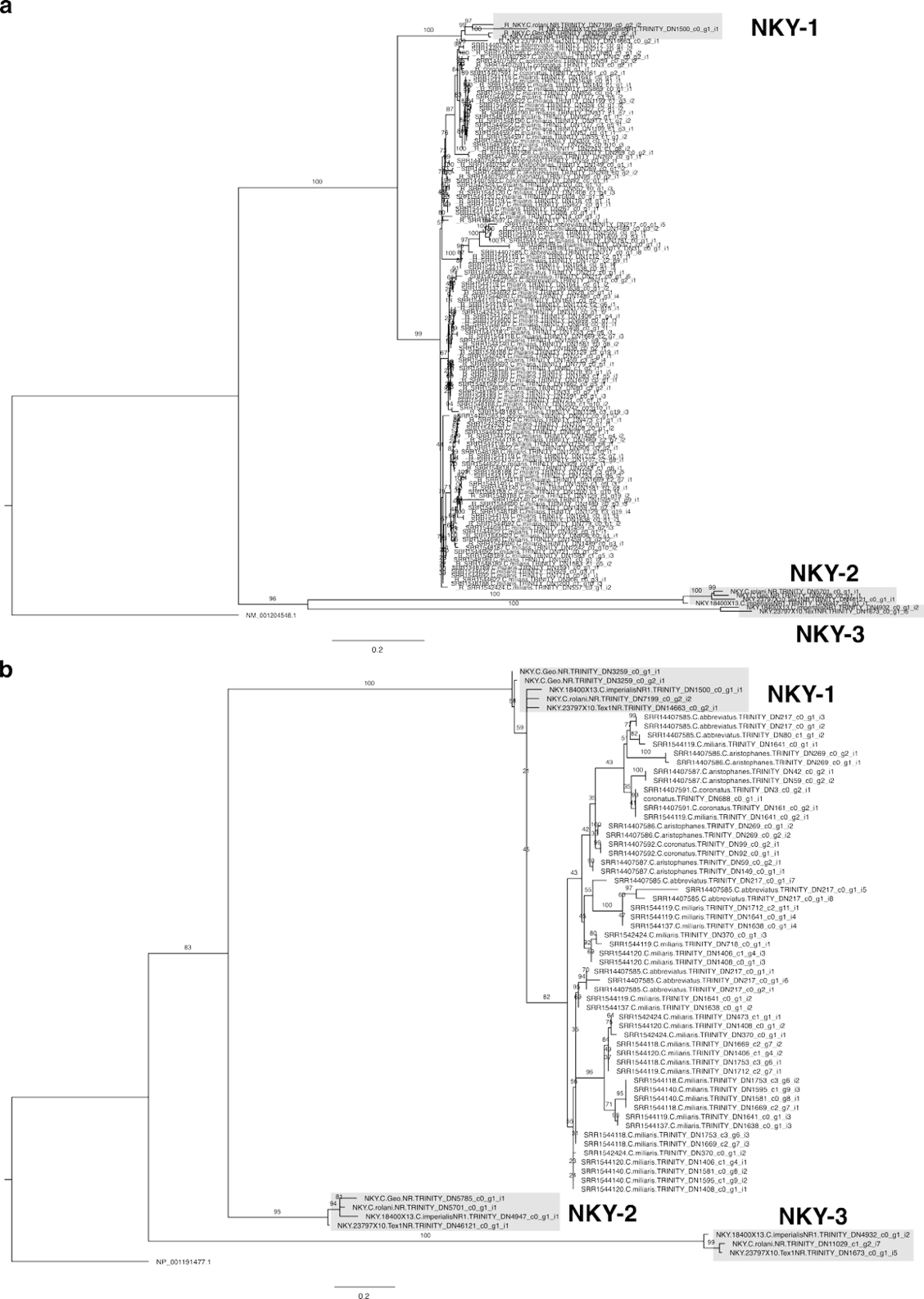

**Fig. S24. NKY doppelgänger evolution. (A)** Maximum likelihood tree reconstruction of nucleotide alignment of NKY neuropeptides and doppelgänger toxins. Using ModelSelector, the model of evolution was chosen to be K2P+R2 based on the Bayesian Information Criterion. The tree was outgrouped with *A. californica* NKY. (B) Maximum likelihood tree reconstruction of protein alignment of NKY neuropeptides and doppelgänger toxins. Using ModelSelector, the Q.mammal+G4 model of evolution was chosen based on the Bayesian Information Criterion. The tree was outgrouped with *A. californica* NKY.

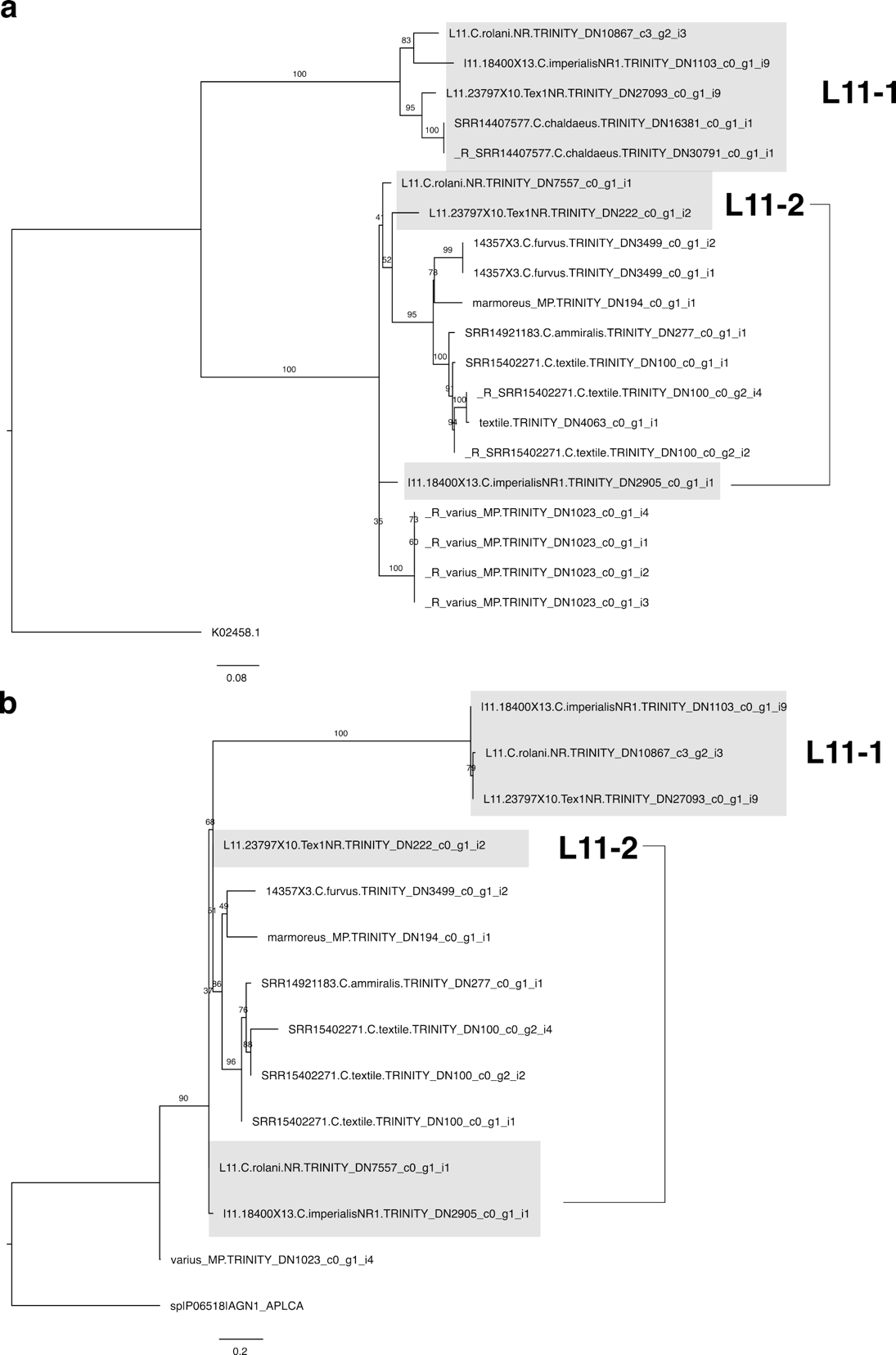
 **Fig. S25. L11 doppelgänger evolution. (A)** Maximum likelihood tree reconstruction of nucleotide alignment of L11 neuropeptides and doppelgänger toxins. Using ModelSelector, the K3Pu+F+G4 model of evolution was chosen based on the Bayesian Information Criterion. Tree was outgrouped with *A. californica* L11. (b) Maximum likelihood tree reconstruction of protein alignment of L11 neuropeptides and doppelgänger toxins. Using ModelSelector, the Q.mammal+G4 model of evolution was chosen based on the Bayesian Information Criterion. The tree was outgrouped with *A. californica* L11.

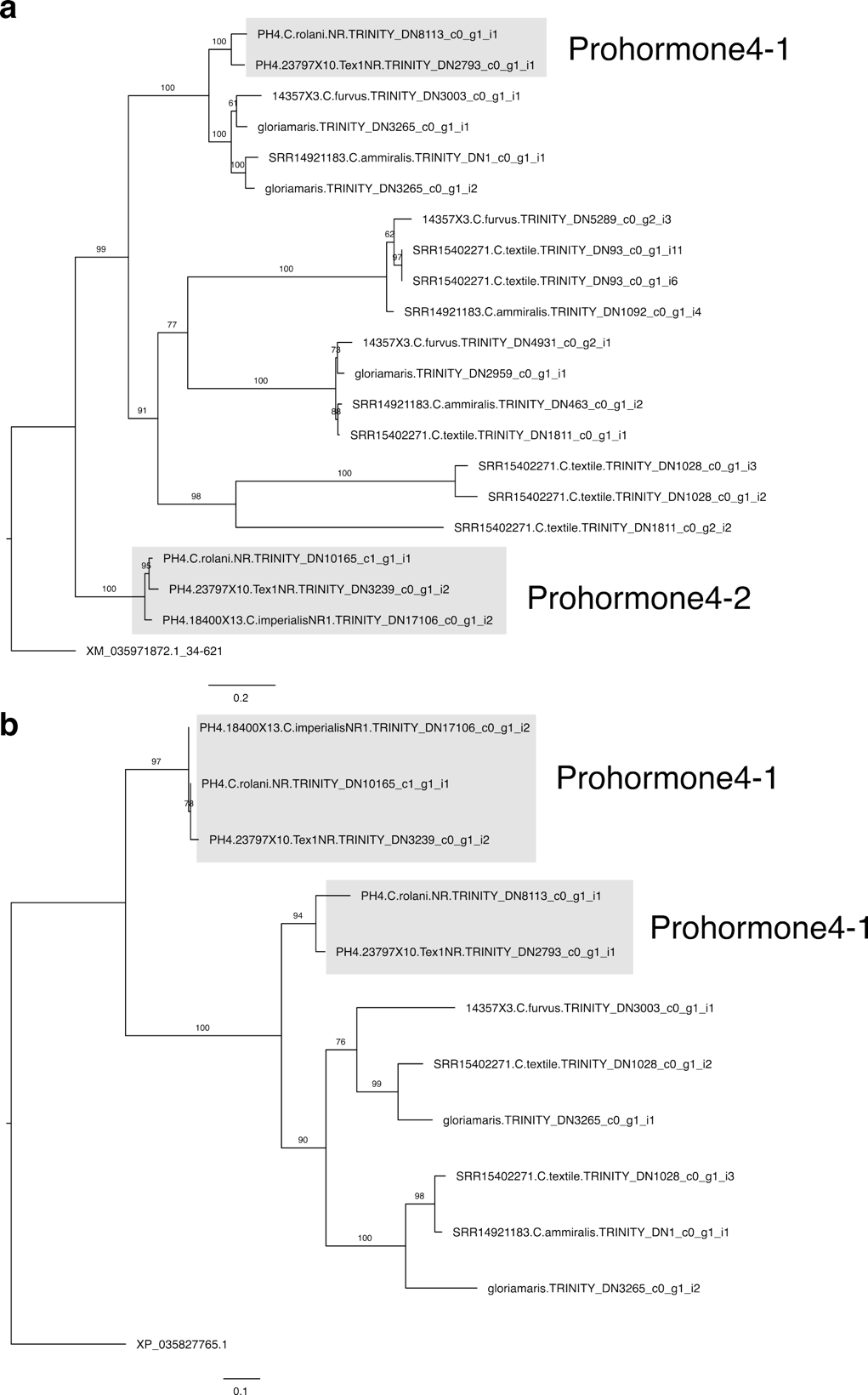

**Fig. S26. Prohormone-4 doppelgänger evolution.** Maximum likelihood tree reconstruction of nucleotide alignment of Prohormone-4 neuropeptides and doppelgänger toxins. Using ModelSelector, the TPM3+G4 model of evolution was chosen based on the Bayesian Information Criterion. The tree was outgrouped with *A. californica* Prohormone-4. (b) Maximum likelihood tree reconstruction of protein alignment of Prohormone-4 neuropeptides and doppelgänger toxins. Using ModelSelector, the JTTDCMut+G4 model of evolution was chosen based on the Bayesian Information Criterion. The tree was outgrouped with *A. californica* Prohormone-4.

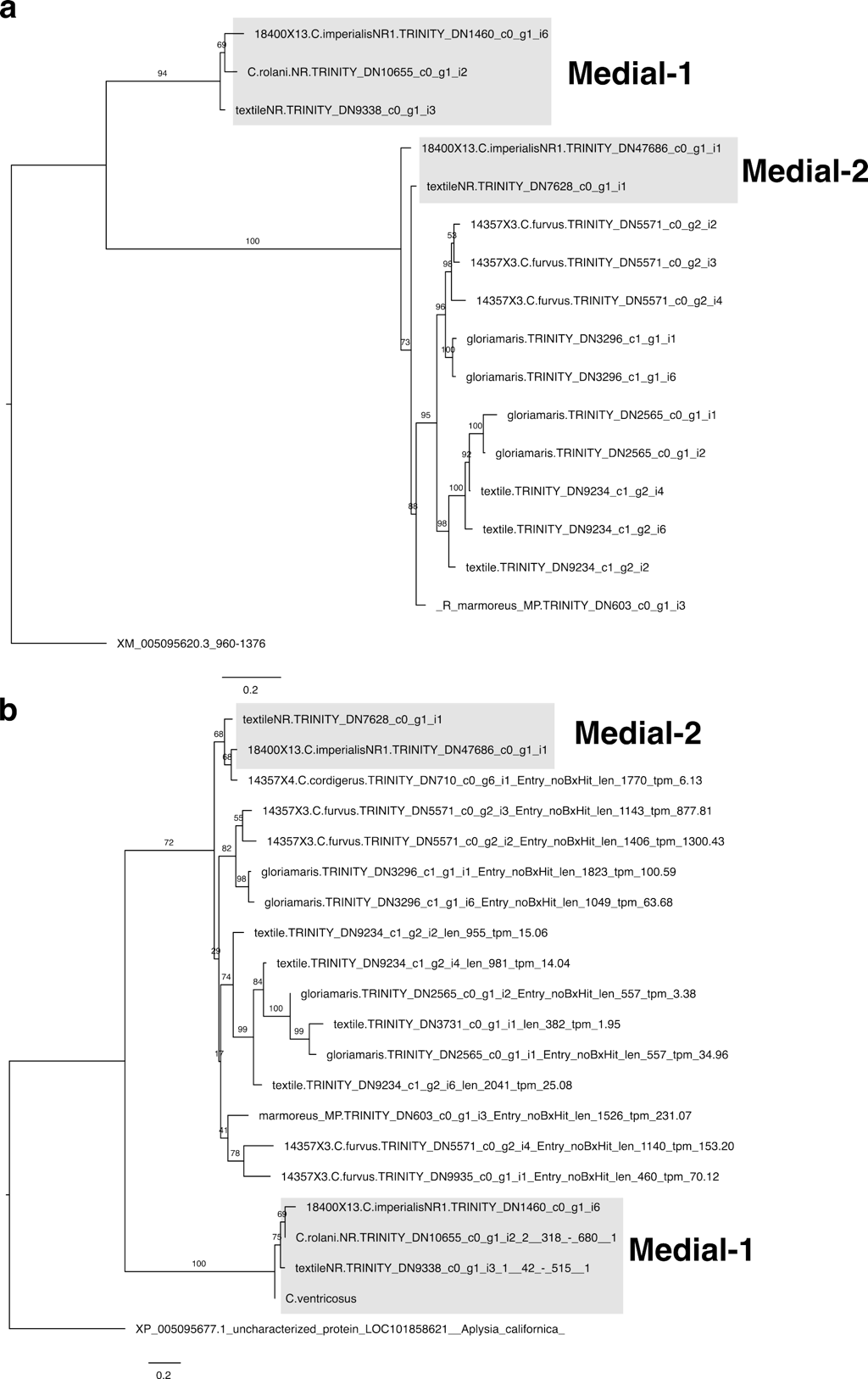

**Fig. S27. Medial doppelgänger evolution. (A)** Maximum likelihood tree reconstruction of nucleotide alignment of Medial DREP neuropeptides and doppelgänger toxins. Using ModelSelector, the HKY+F+R2 model of evolution was chosen based on the Bayesian Information Criterion. The tree was outgrouped with *A. californica* Medial DREP. (B) Maximum likelihood tree reconstruction of protein alignment of Medial DREP neuropeptides and doppelgänger toxins. Using ModelSelector, the JTT+G4 model of evolution was chosen based on the Bayesian Information Criterion. The tree was outgrouped with *A. californica* Medial DREP.

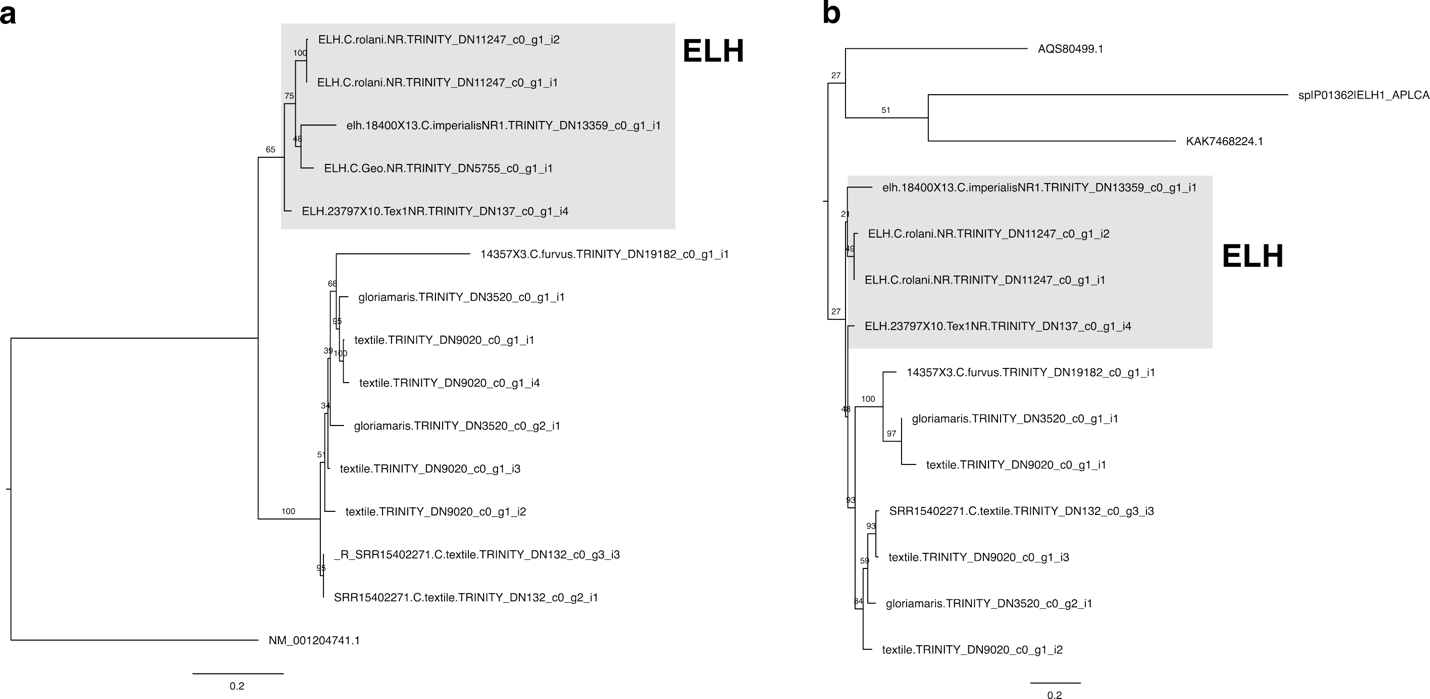

**Fig. S28. Egg-laying hormone (ELH) doppelgänger evolution. (A)** Maximum likelihood tree reconstruction of nucleotide alignment of ELH neuropeptides and doppelgänger toxins. Using ModelSelector, the Q.mammal+G4 model of evolution was chosen based on the Bayesian Information Criterion. The tree was outgrouped with *A. californica* ELH. (B) Maximum likelihood tree reconstruction of protein alignment of ELH neuropeptides and doppelgänger toxins. Using ModelSelector, the HKY+F+R3 model of evolution was chosen based on the Bayesian Information Criterion. The tree was outgrouped with several non-*Conus* molluscan ELH, including *A. californica* ELH.

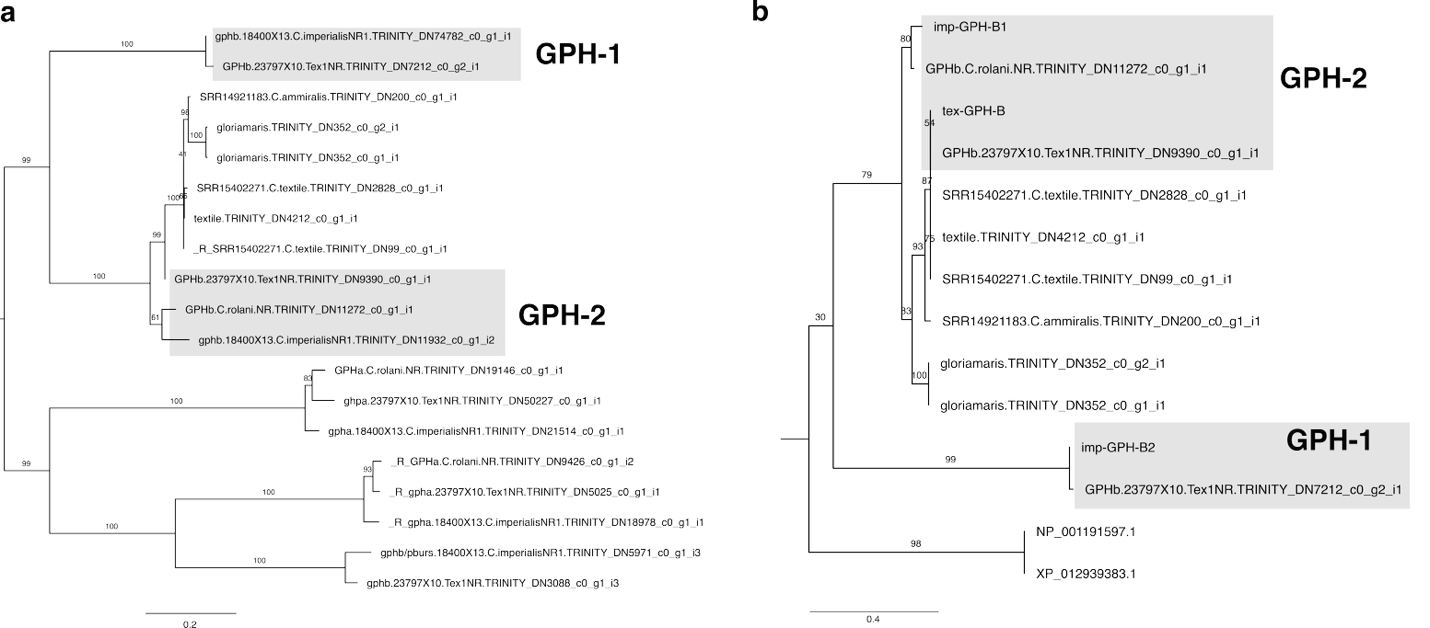

**Fig. S29. Glycoprotein hormone doppelgänger evolution. (A)** Maximum likelihood tree reconstruction of nucleotide alignment of GPH neuropeptides and doppelgänger toxins. Using ModelSelector, the K2P+G4 model of evolution was chosen based on the Bayesian Information Criterion. The tree was outgrouped with *Conus* GPH proteins. (B) Maximum likelihood tree reconstruction of protein alignment of GPH neuropeptides and doppelgänger toxins. Using ModelSelector, the Q.mammal+G4 model of evolution was chosen based on the Bayesian Information Criterion. The tree was outgrouped with several non-*Conus* molluscan GPH, including *A. californica* GPH.

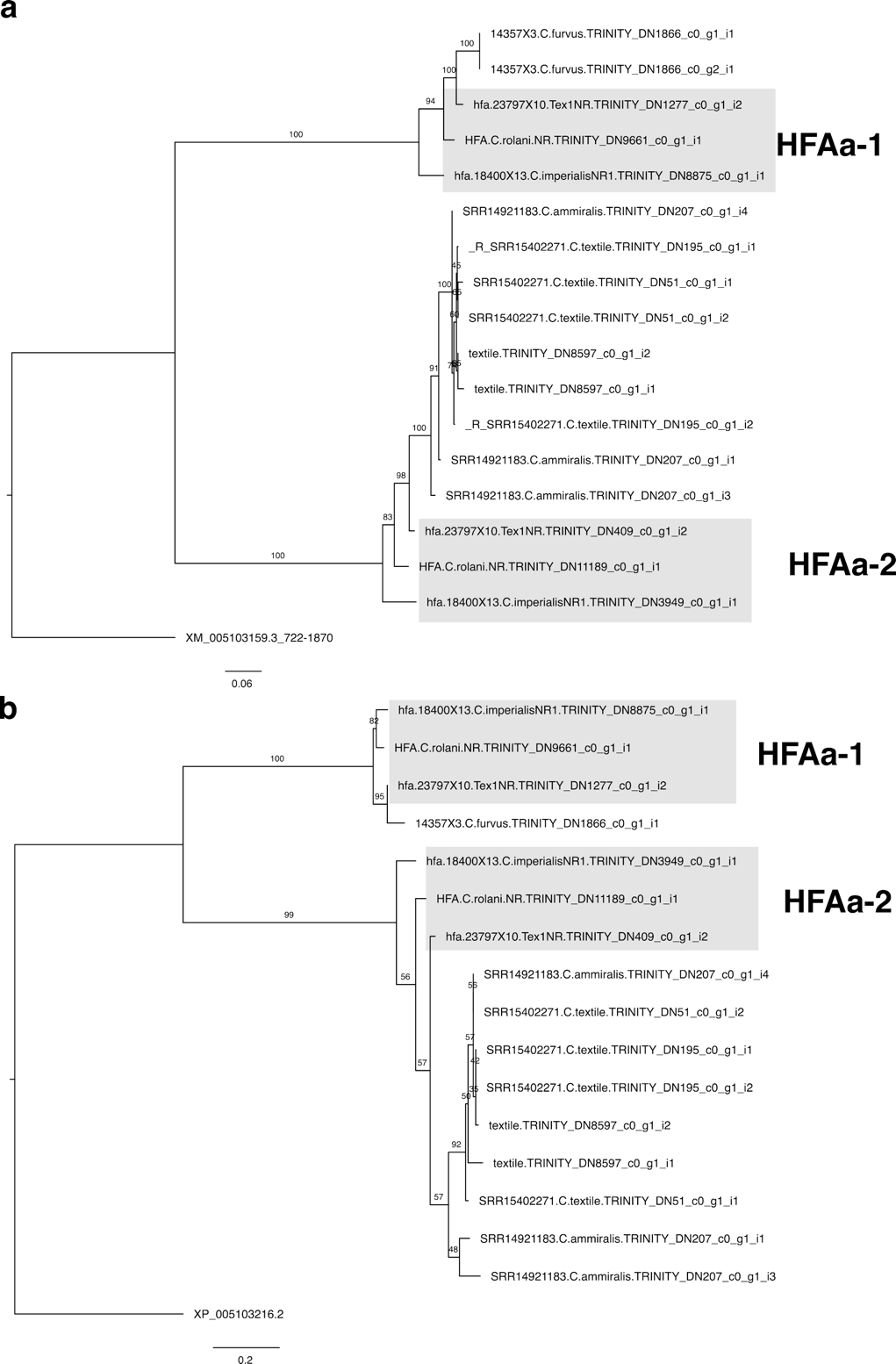

**Fig. S30. HFAamide doppelgänger evolution. (A)** Maximum likelihood tree reconstruction of nucleotide alignment of HFAamide neuropeptides and doppelgänger toxins. Using ModelSelector, the HKY+F model of evolution was chosen based on the Bayesian Information Criterion. The tree was outgrouped with *A. californica* HFAamide. (B) Maximum likelihood tree reconstruction of protein alignment of HFAamide neuropeptides and doppelgänger toxins. Using ModelSelector, the JTT+I model of evolution was chosen based on the Bayesian Information Criterion. The tree was outgrouped with *A. californica* HFAamide.

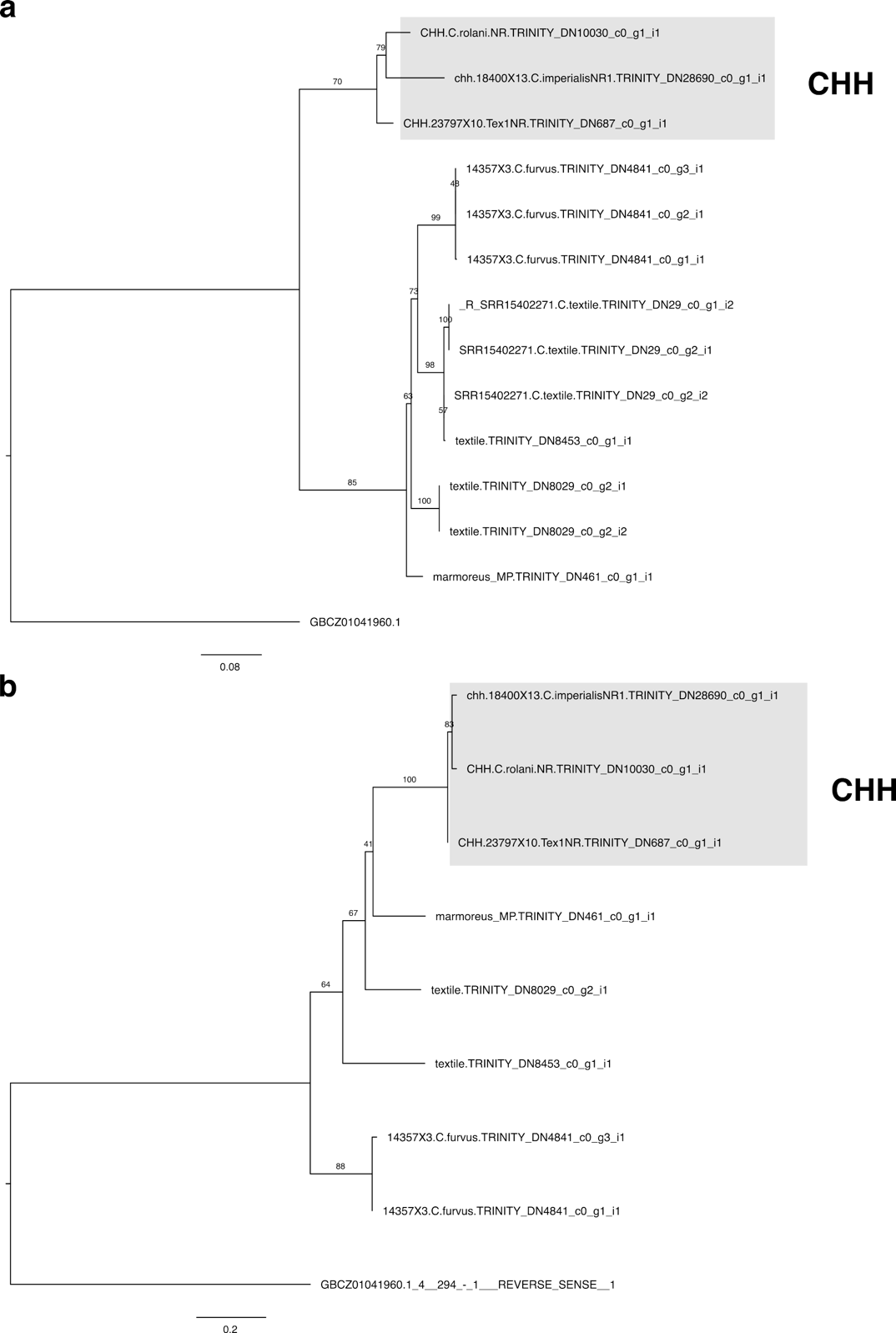

**Fig. S31. Crustacean Hypoglycemic Hormone doppelgänger evolution. (A)** Maximum likelihood tree reconstruction of nucleotide alignment of CHH neuropeptides and doppelgänger toxins. Using ModelSelector, the TPM3+I model of evolution was chosen based on the Bayesian Information Criterion. The tree was outgrouped with *A. californica* CHHe. B) Maximum likelihood tree reconstruction of protein alignment of CHH neuropeptides and doppelgänger toxins. Using ModelSelector, the JTTDCMut+G4 model of evolution was chosen based on the Bayesian Information Criterion. The tree was outgrouped with *A. californica* CHH.

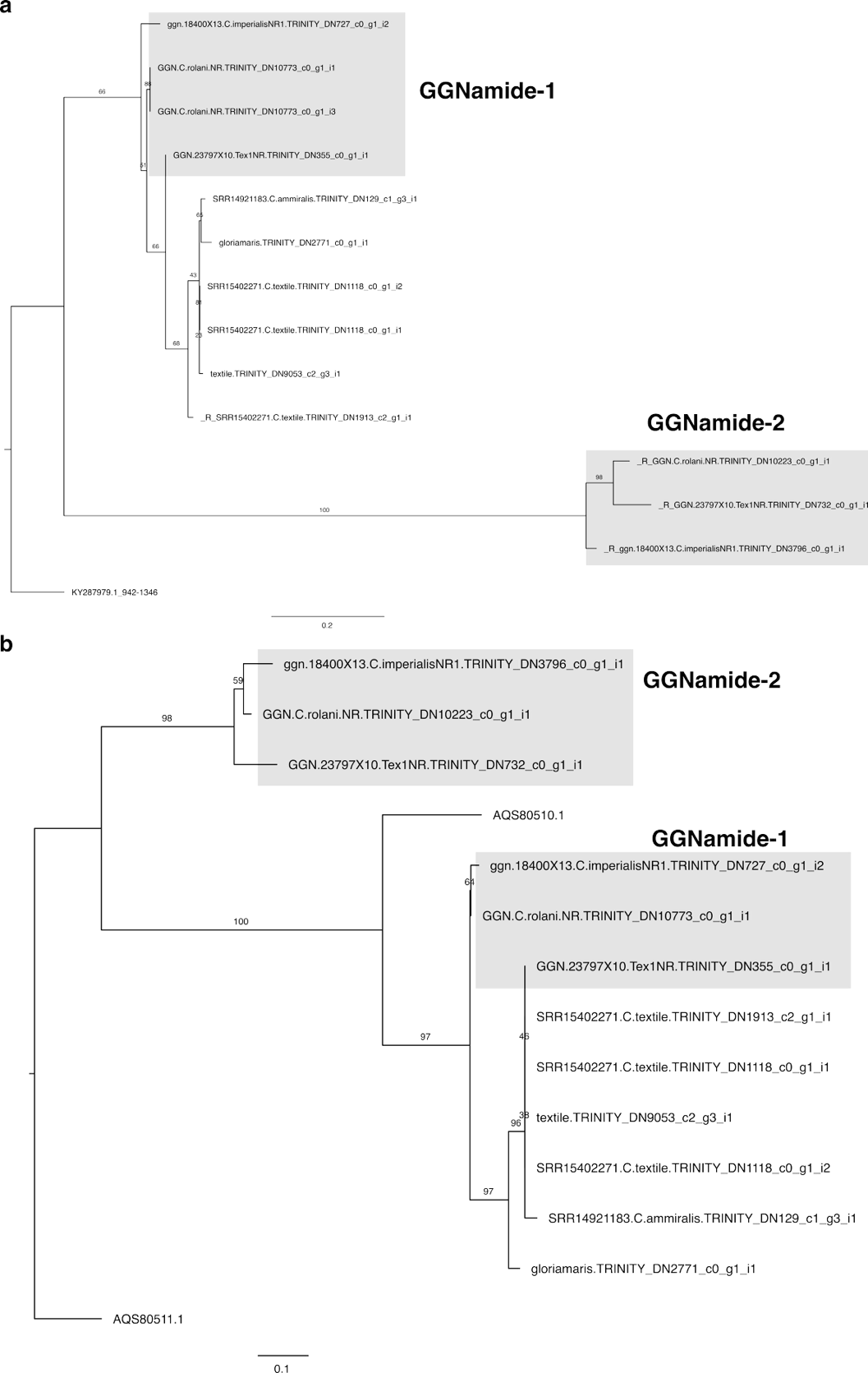

**Fig. S32. GGNamide doppelgänger evolution. (A)** Maximum likelihood tree reconstruction of nucleotide alignment of GGNamide neuropeptides and doppelgänger toxins. Using ModelSelector, the K2P+I model of evolution was chosen based on the Bayesian Information Criterion. The tree was outgrouped with *A. californica* GGNamide. (B) Maximum likelihood tree reconstruction of protein alignment of GGNamide neuropeptides and doppelgänger toxins. Using ModelSelector, the Q.mammal+G4 model of evolution was chosen based on the Bayesian Information Criterion. The tree was outgrouped with *A. californica* GGNamide.

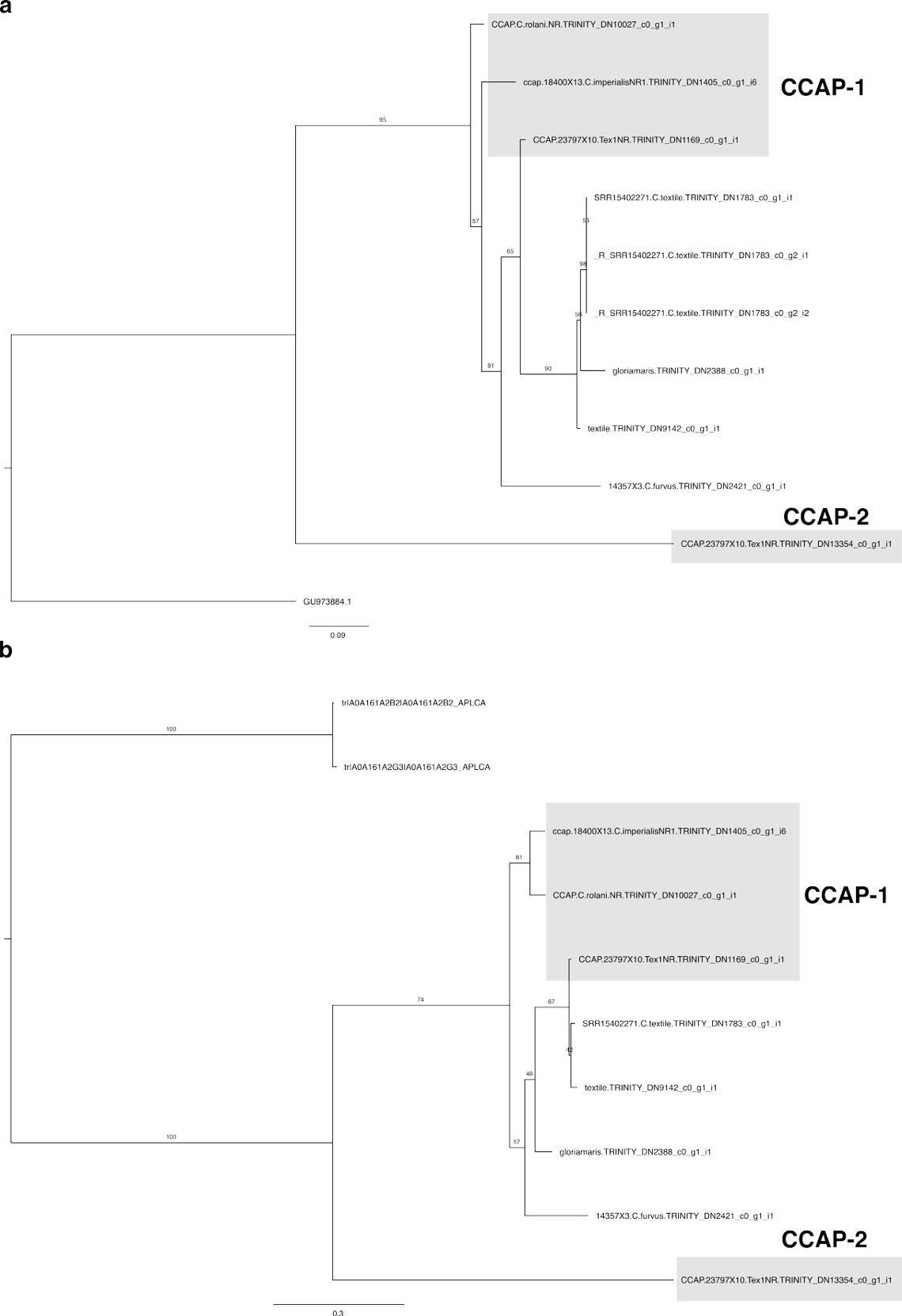

**Fig. S33. CCAP doppelgänger evolution. (A)** Maximum likelihood tree reconstruction of nucleotide alignment of GGNamide neuropeptides and conoCAP doppelgänger toxins. Using ModelSelector, the K2P+G4 model of evolution was chosen based on the Bayesian Information Criterion. The tree was outgrouped with *A. californica* CCAP. (B) Maximum likelihood tree reconstruction of protein alignment of GGNamide neuropeptides and conoCAP doppelgänger toxins. Using ModelSelector, the JTT+G4 model of evolution was chosen based on the Bayesian Information Criterion. The tree was outgrouped with *A. californica* CCAP.

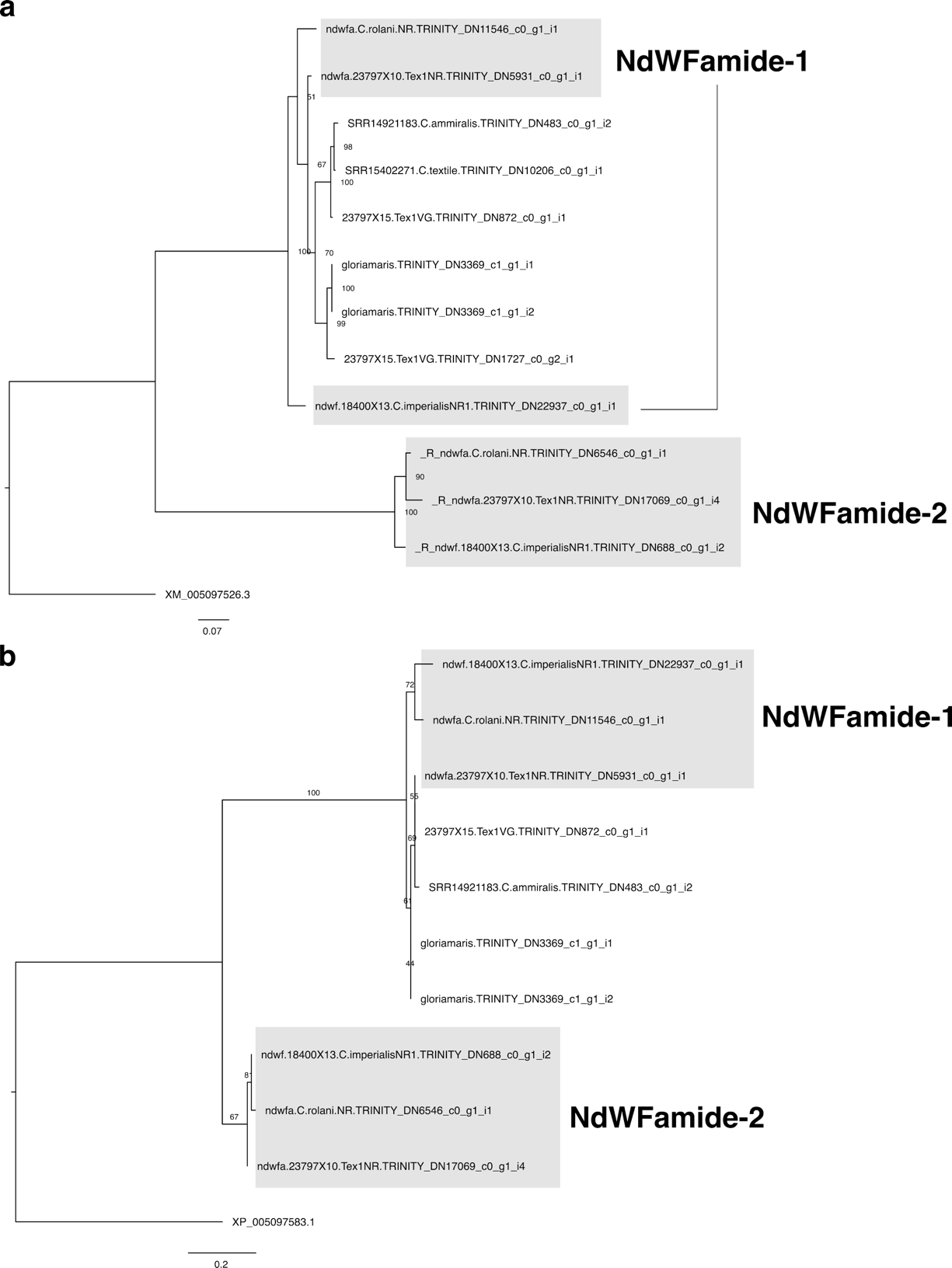

**Fig. S34. NdWFamide doppelgänger evolution. (A)** Maximum likelihood tree reconstruction of nucleotide alignment of NdWFamide neuropeptides and doppelgänger toxins. Using ModelSelector, the HKY+F model of evolution was chosen based on the Bayesian Information Criterion. The tree was outgrouped with *A. californica* NdWFamide. (B) Maximum likelihood tree reconstruction of protein alignment of NdWFamide neuropeptides and doppelgänger toxins. Using ModelSelector, the Q.plant+G4 model of evolution was chosen based on the Bayesian Information Criterion. The tree was outgrouped with *A. californica* NdWFamide.

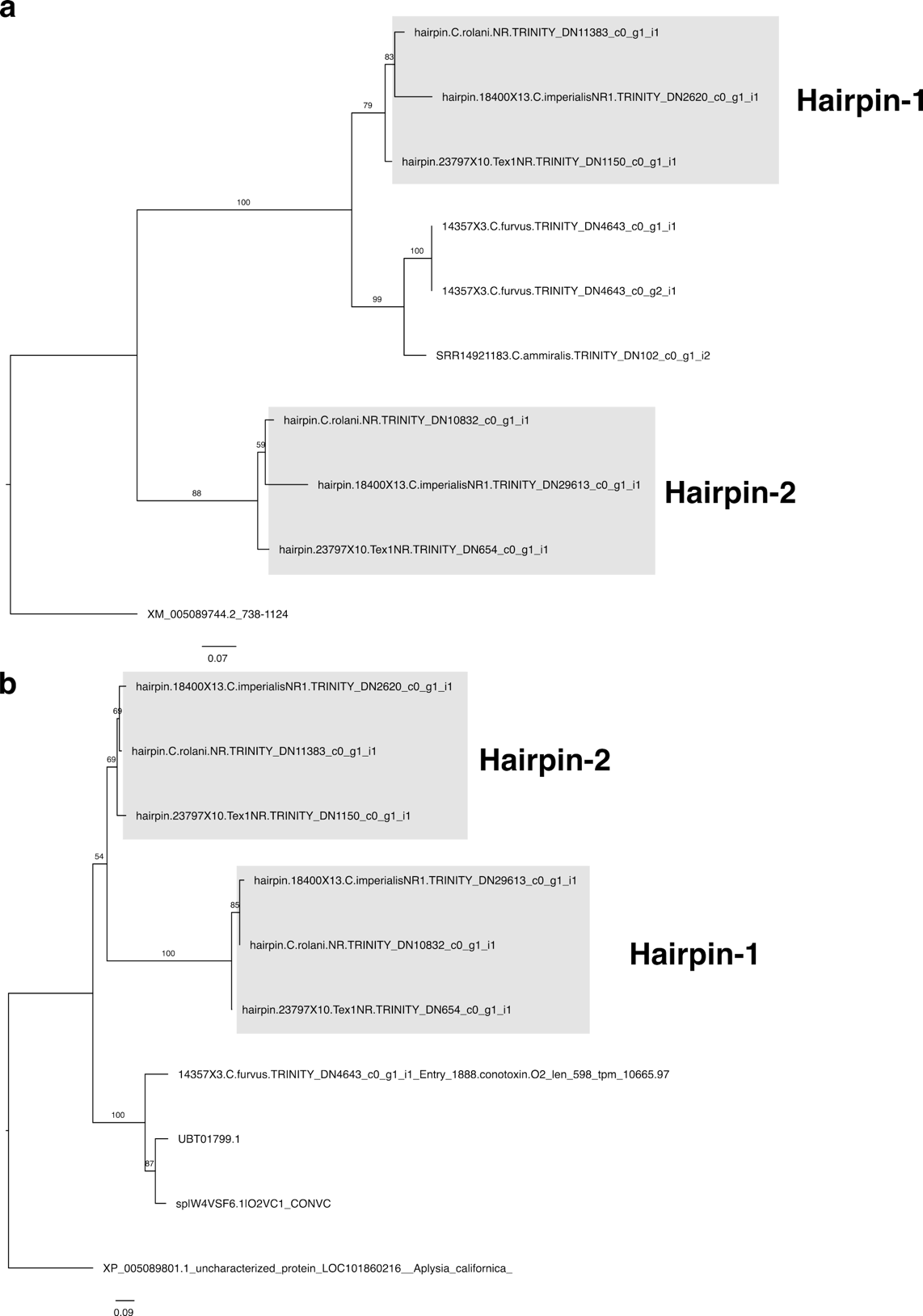

**Fig. S35. Hairpin doppelgänger evolution. (A)** Maximum likelihood tree reconstruction of nucleotide alignment of Hairpin neuropeptides and doppelgänger toxins. Using ModelSelector, the K2P+I model of evolution was chosen based on the Bayesian Information Criterion. Tree was outgrouped with *A. californica* Hairpin. (B) Maximum likelihood tree reconstruction of protein alignment of Hairpin neuropeptides and doppelgänger toxins. Using ModelSelector, the WAG model of evolution was chosen based on the Bayesian Information Criterion. The tree was outgrouped with *A. californica* Hairpin.

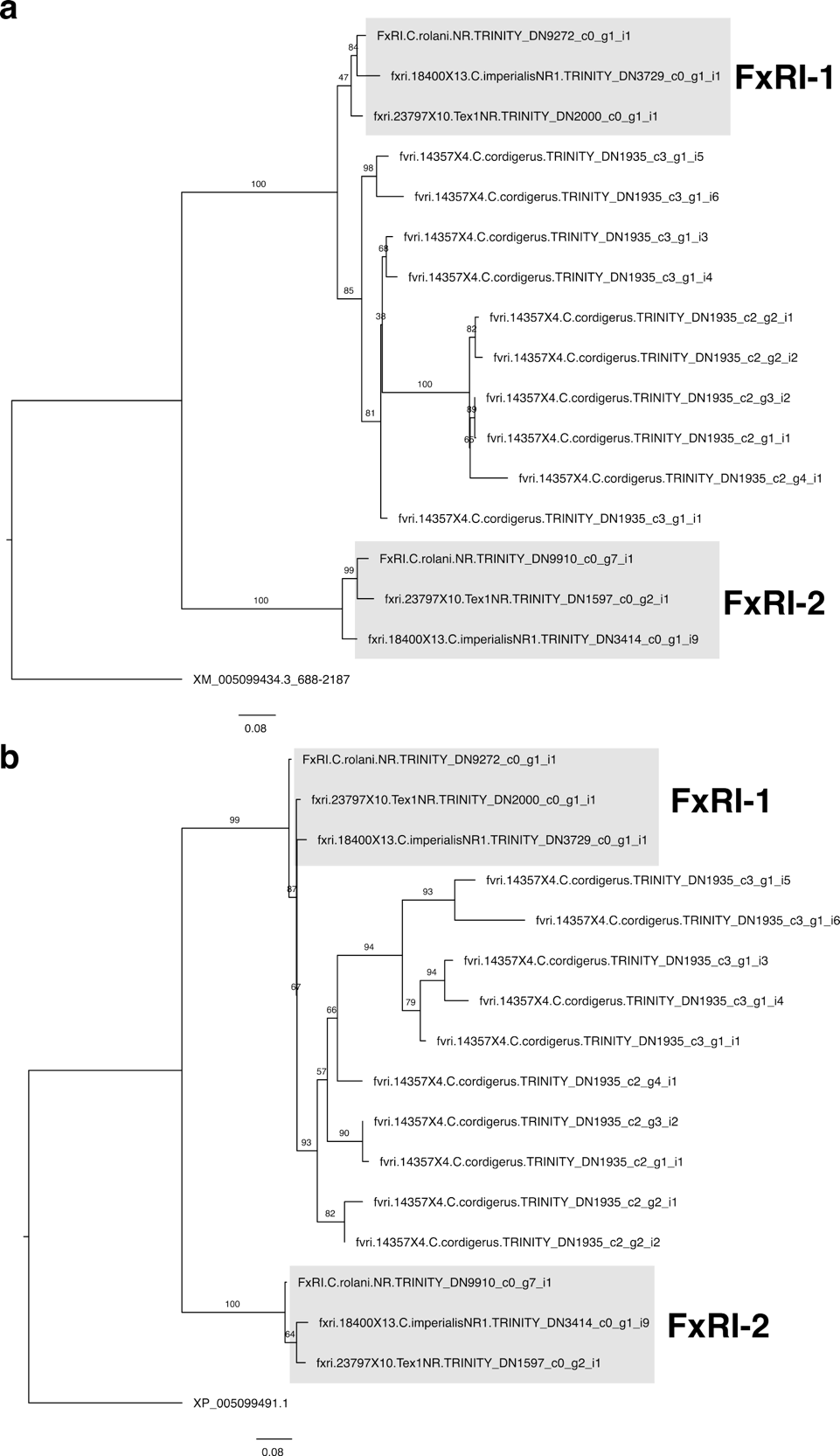

**Fig. S36. FxRIamide doppelgänger evolution. (A)** Maximum likelihood tree reconstruction of nucleotide alignment of FxRIamide neuropeptides and doppelgänger toxins. Using ModelSelector, the K2P+G4 model of evolution was chosen based on the Bayesian Information Criterion. The tree was outgrouped with *A. californica* FxRIamide. (B) Maximum likelihood tree reconstruction of protein alignment of FxRIamide neuropeptides and doppelgänger toxins. Using ModelSelector, the JTT+F+G4 model of evolution was chosen based on the Bayesian Information Criterion. The tree was outgrouped with *A. californica* FxRIamide.

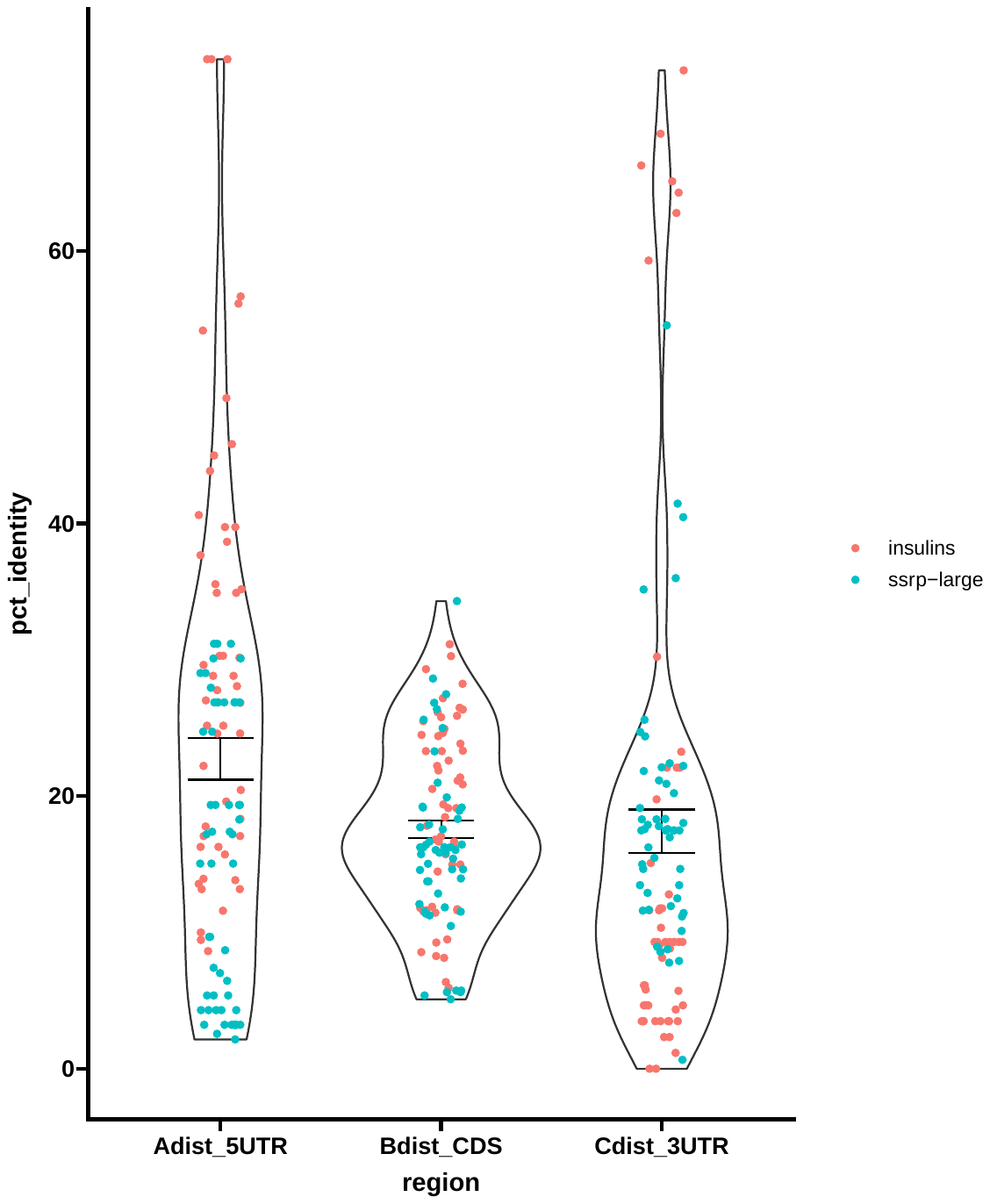

**Fig. S37. Similarity of 5’ UTR, CDS, and 3’ UTR transcript regions between insulin and SSRP doppelgängers with the most similar neuropeptide transcript.** Unlike more recently evolved doppelgänger toxins, con-insulins and consomatins show little difference between their transcript regions.

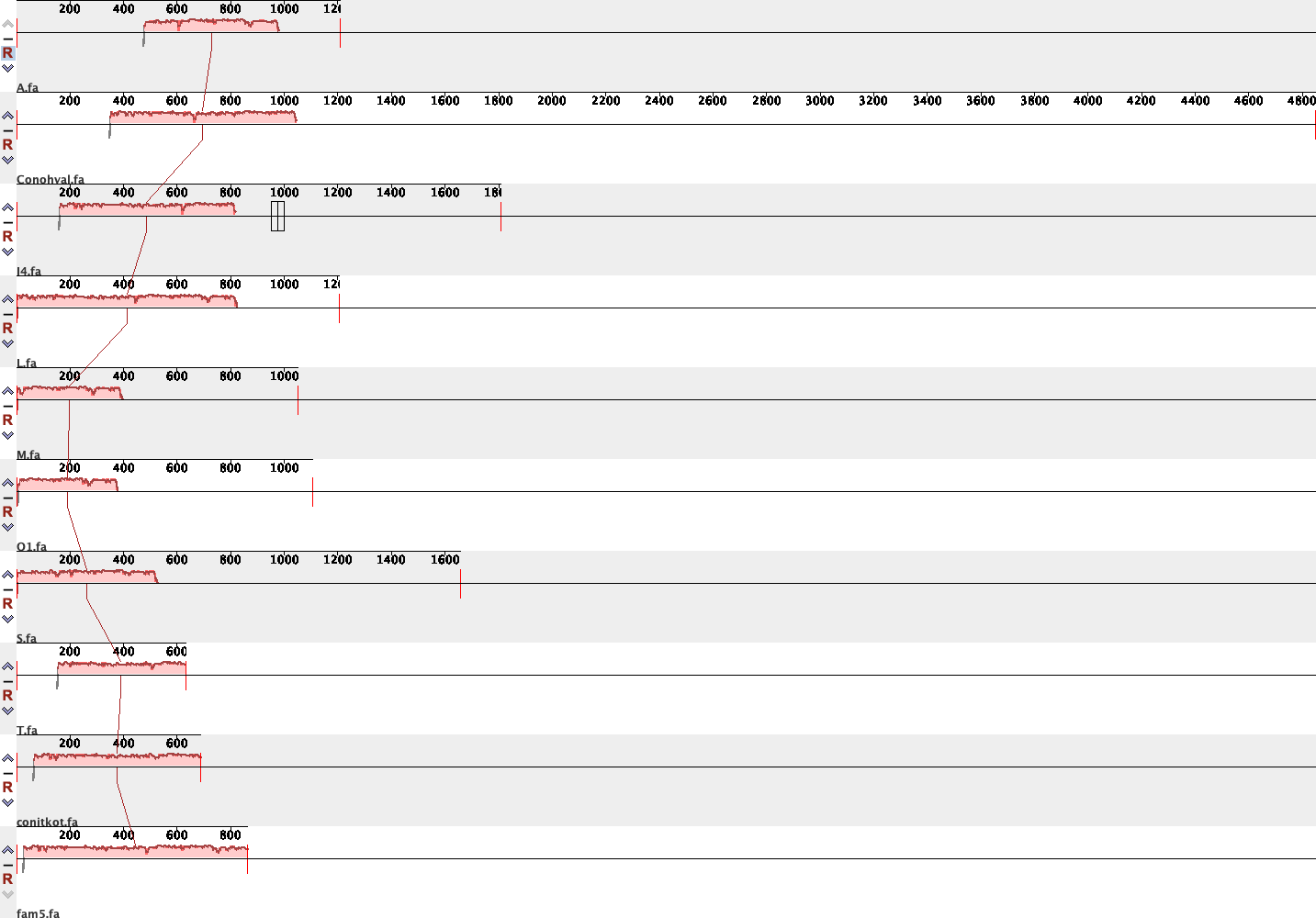

**Fig. S38. Expressed retrotransposon-containing conotoxin 5’ UTRs.** The Family-5 transposable element is found across members of nine different conotoxin superfamilies.

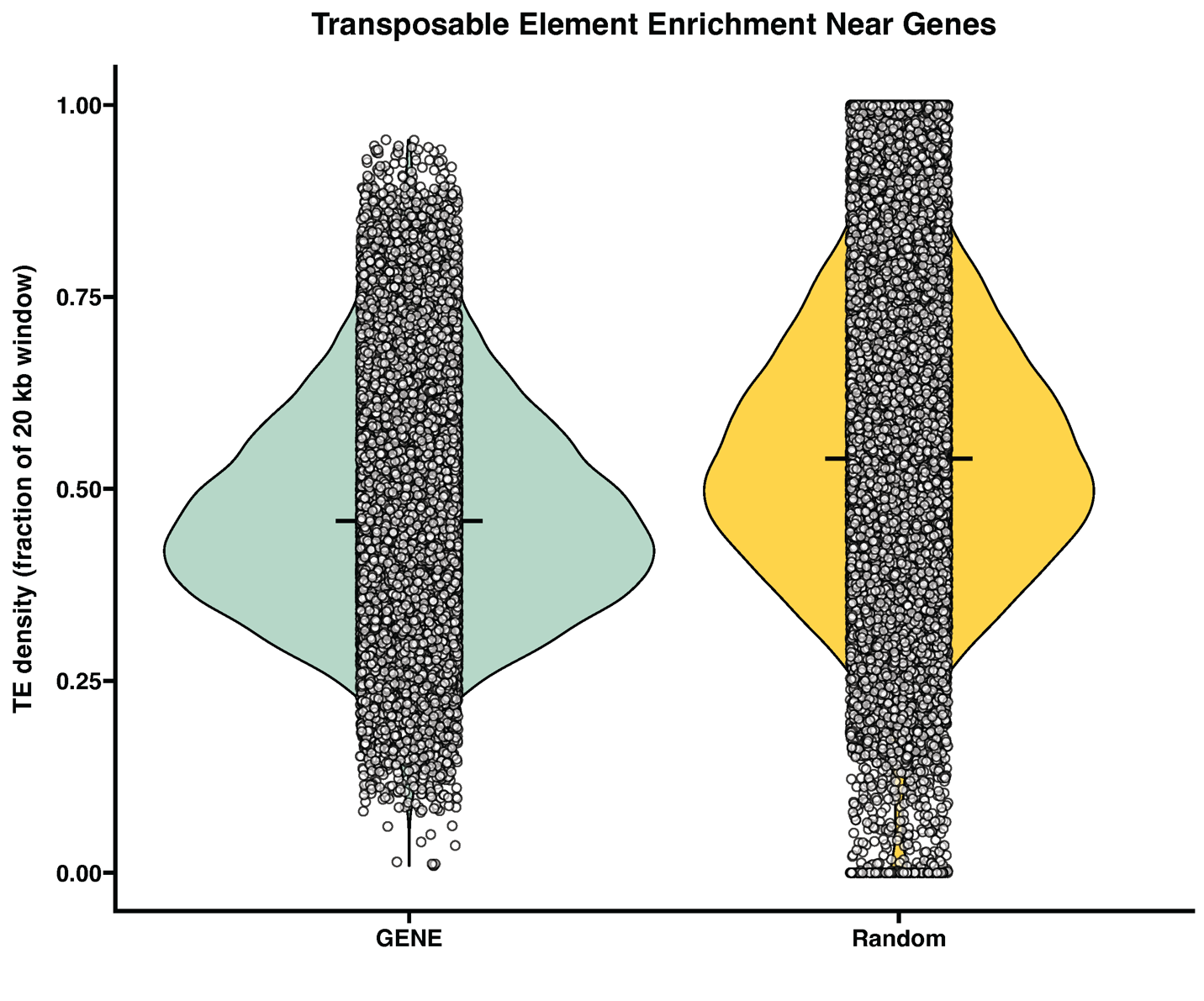

**Fig. S39. TE density surrounding braker annotated genes in *C. textile* and randomly shuffled gene locations.** There is a lower density of TEs surrounding annotated genes compared to random regions in the genome (p = 2.2e-16, Wilcoxon rank sum test).

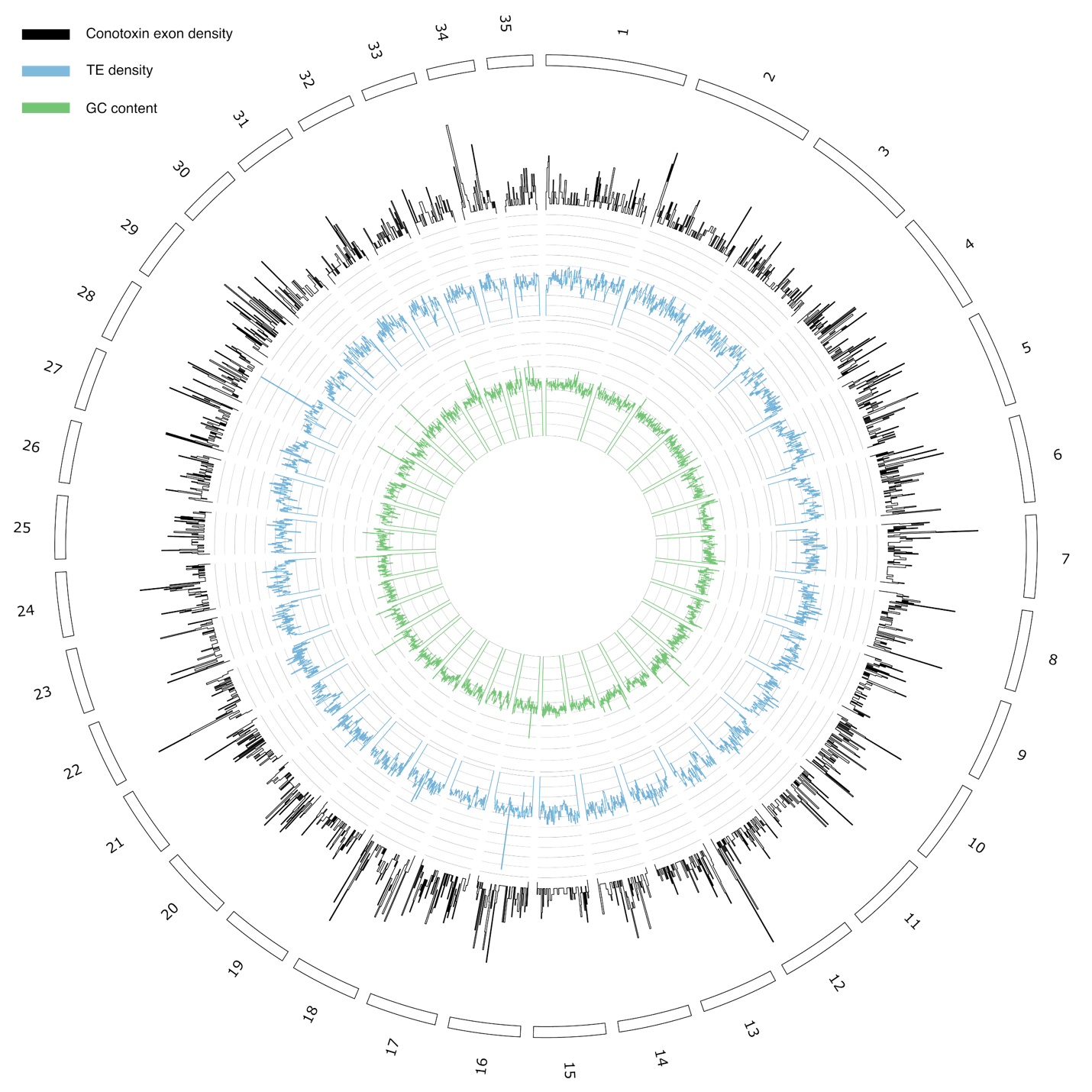
**Fig. S40. Conotoxin exon density, TE density, and GC content in *C. textile* genome.** The conotoxin and TE density, as well as the GC content were calculated in 1Mb subsets of the genome and visualized using Circos.

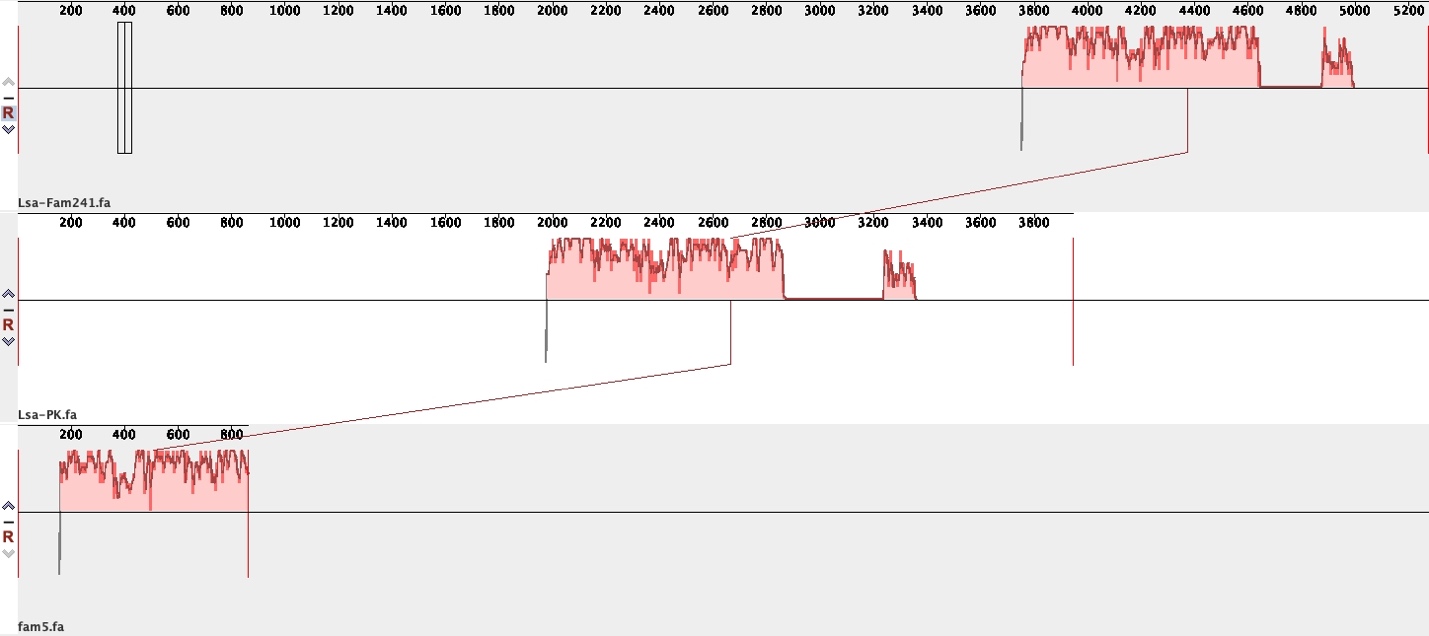

**Fig. S41. Alignment of *C. textile* TE family-5 with mRNA from *Littorina saxatilis* proteins.** The *L. saxatilis* mRNA encoding divergent protein kinase domain 2A-like (NCBI: XM_070356896.1) and protein FAM241B-like (NCBI: XM_070330134.1) are predicted to contain family-5 like TEs.

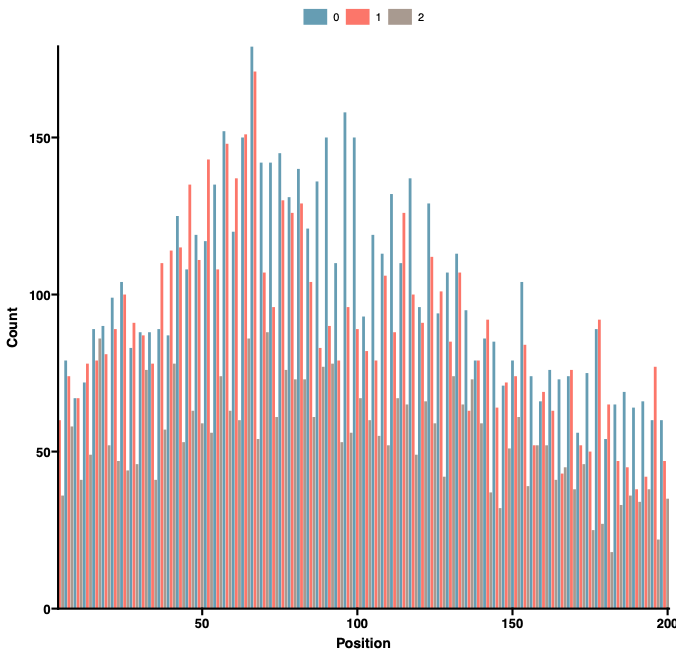

**Fig. S42. Intron phases in non-secreted proteins.** Phase and locations of first introns in non-secreted *C. textile* proteins. The proportions are Phase-0: 41%, Phase-1: 37%, and Phase-2: 22%.

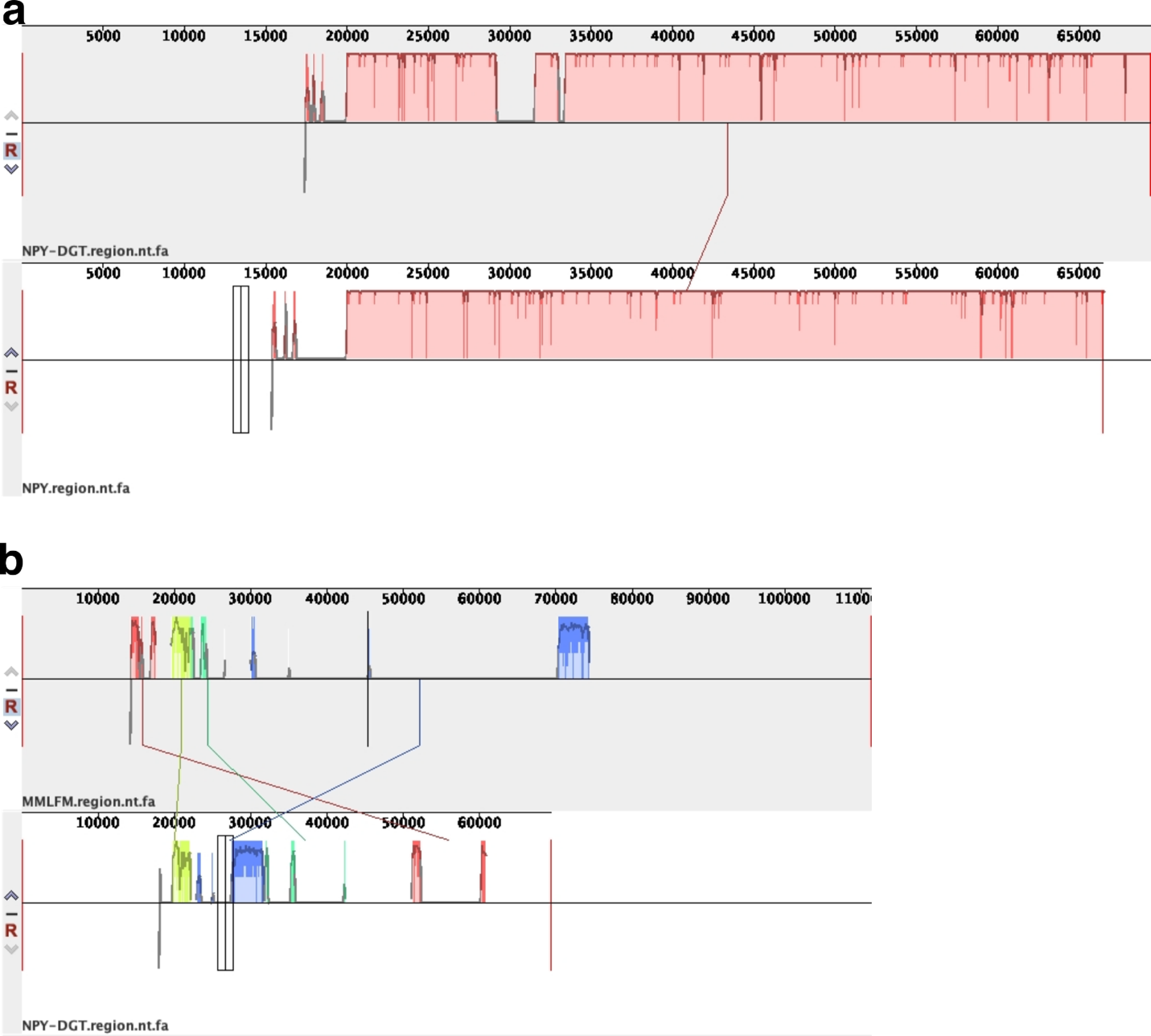

**Fig. S43. Alignment of MMLFM, conoNPY, and NPF/Y loci in *C. textile.*** All genes begin 20kbp into the alignment and extend through the end of the alignment. (A) Alignment between conoNPY (upper) and NPF/Y (lower) loci in the genome. (B) Alignment between MMLFM toxin loci (upper) and conoNPY (lower) loci in the genome.

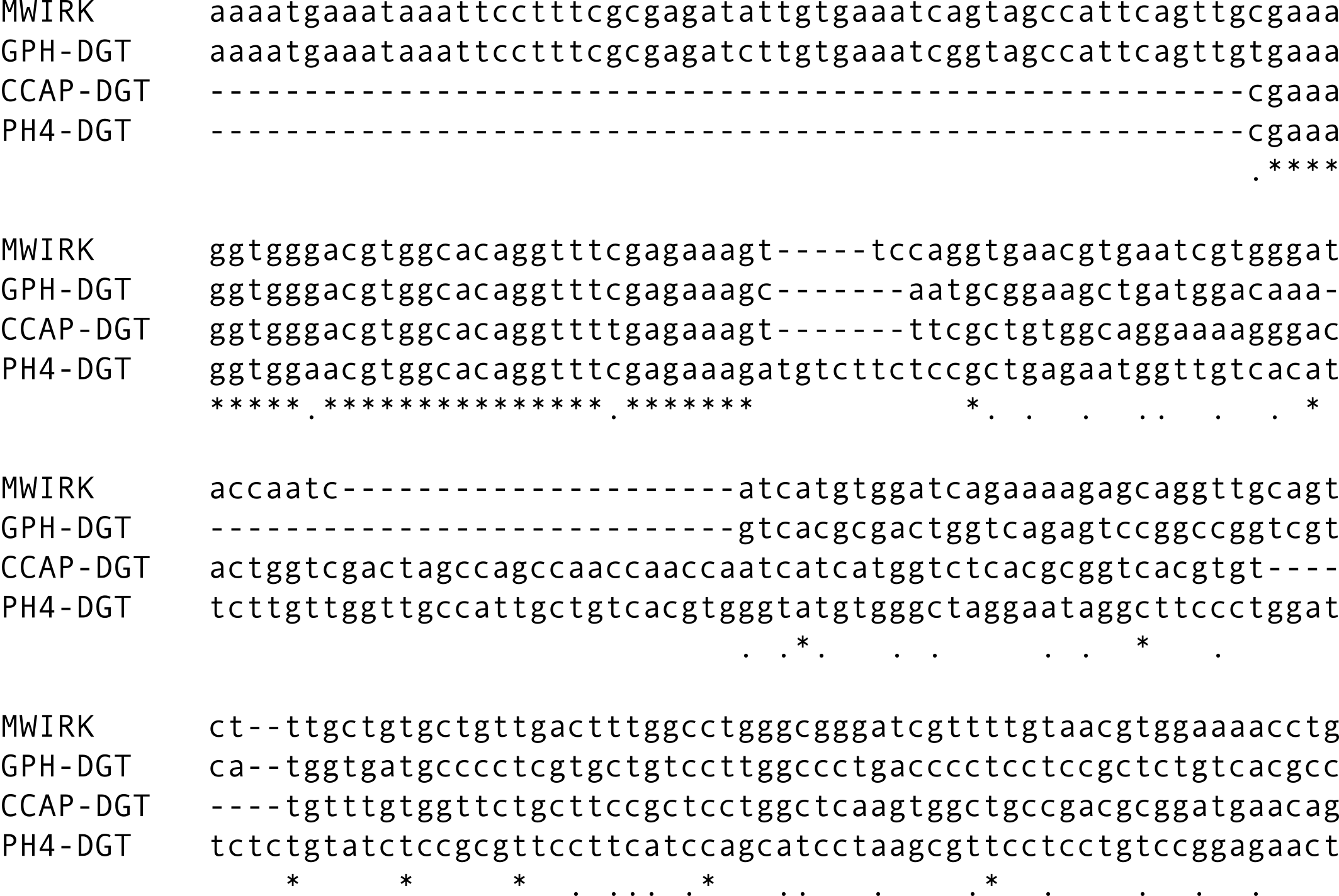

**Fig. S44. Multiple sequence alignment of *C. furvus* MWIRK conotoxin and *C. textile* GPH, CCAP, and Prohormone-4 doppelgänger toxin transcripts.** All sequences share a common 5’ UTR (marked with stars), particularly between MWIRK and the GPH doppelgänger toxins.

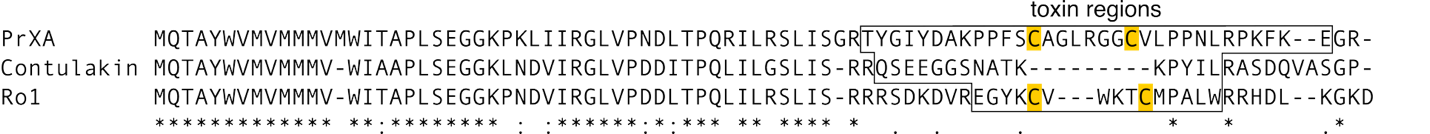

**Fig. S45. Multiple sequence alignment of C superfamily toxins.** Consomatin Ro1 resembles the ancestral SSRP neuropeptide that was initially recruited into the venom and represents a SSRP doppelgänger toxin. Following recruitment, the SSRP superfamily diversified to include Contulakin-G, a neurotensin receptor agonist, and PrXA, a nicotinic acetylcholine receptor antagonist. Cysteine connected by disulfide bond are highlighted yellow. The gene superfamily that encompasses all these toxins is known as the C superfamily.

Table S1.

**Assembly statistics**

| Assembly | C_textile_scaffolded.yes-cor.v2.fa.masked |
| --- | --- |
| # contigs (>= 0 bp) | 2618 |
| # contigs (>= 1000 bp) | 2615 |
| # contigs (>= 5000 bp) | 2515 |
| # contigs (>= 10000 bp) | 2483 |
| # contigs (>= 25000 bp) | 1784 |
| # contigs (>= 50000 bp) | 1268 |
| Total length (>= 0 bp) | 3550089287 |
| Total length (>= 1000 bp) | 3550088231 |
| Total length (>= 5000 bp) | 3549942712 |
| Total length (>= 10000 bp) | 3549712027 |
| Total length (>= 25000 bp) | 3537168836 |
| Total length (>= 50000 bp) | 3519148590 |
| # contigs | 2616 |
| Largest contig | 179254051 |
| Total length | 3550088828 |
| GC (%) | 43.56 |
| N50 | 91776639 |
| N90 | 60457912 |
| auN | 94127367.6 |
| L50 | 16 |
| L90 | 34 |
| # N's per 100 kbp | 95.53 |

Table S2. Terrier annotation of transposable elements

| **Class** | **Total length** | **Percent** |
| --- | --- | --- |
| DNA | 435560601 | 12.269313 |
| DNA/Academ | 18456561 | 0.519903 |
| DNA/CMC | 12568272 | 0.354036 |
| DNA/Dada | 1442689 | 0.040639 |
| DNA/Harbinger | 27531940 | 0.775548 |
| DNA/Kolobok | 1865086 | 0.052538 |
| DNA/MULE | 20595748 | 0.580162 |
| DNA/Maverick | 17551033 | 0.494395 |
| DNA/Merlin | 167329 | 0.004713 |
| DNA/PiggyBac | 6731327 | 0.189615 |
| DNA/Sola | 5476964 | 0.154281 |
| DNA/TcMar | 45320170 | 1.276625 |
| DNA/hAT | 33186656 | 0.934835 |
| LINE | 30695838 | 0.864671 |
| LINE/CR1 | 28822714 | 0.811907 |
| LINE/CRE | 228257 | 0.00643 |
| LINE/I | 125804702 | 3.543794 |
| LINE/L1 | 36720358 | 1.034376 |
| LINE/L2 | 29678001 | 0.836 |
| LINE/R1 | 137009 | 0.003859 |
| LINE/R2 | 24404001 | 0.687437 |
| LINE/RTE | 88749879 | 2.499997 |
| LINE/RTE-BovB | 393497 | 0.011084 |
| LINE/Rex-Babar | 714181 | 0.020118 |
| LINE/Tad1 | 4207607 | 0.118524 |
| LTR | 55354216 | 1.559274 |
| LTR/Copia | 1105963 | 0.031154 |
| LTR/DIRS | 94374387 | 2.658433 |
| LTR/ERV | 14552025 | 0.409916 |
| LTR/Gypsy | 129054698 | 3.635344 |
| LTR/Pao | 2180516 | 0.061423 |
| Low_complexity | 18269953 | 0.514647 |
| PLE | 58962997 | 1.660929 |
| RC | 16663500 | 0.469394 |
| SINE/tRNA | 23515569 | 0.66241 |
| Satellite | 7013343 | 0.197559 |
| Simple_repeat | 273777651 | 7.712047 |
| Structural_RNA | 107058 | 0.003016 |
| Unknown | 424055306 | 11.94522 |

Table S3. Metazoan BUSCO on the proteome

C:90.9%[S:34.4%,D:56.5%],F:2.0%,M:7.1%,n:954

867	Complete BUSCOs (C)

328	Complete and single-copy BUSCOs (S)

539	Complete and duplicated BUSCOs (D)

19	Fragmented BUSCOs (F)

68	Missing BUSCOs (M)

954	Total BUSCO groups searched

Table S4.

**Venom composition from transcriptome**

| Superfamily | Transcripts | TPM |
| --- | --- | --- |
| O2 | 40 | 211191.67 |
| T | 29 | 142314.16 |
| O1 | 15 | 102780.53 |
| M | 18 | 83848.5 |
| J | 6 | 40082.84 |
| H | 7 | 37813.08 |
| A | 9 | 34466.73 |
| I1 | 6 | 22709.09 |
| MSRLF | 10 | 14574.79 |
| MKAVA | 1 | 10400.35 |
| I2 | 4 | 10255.12 |
| P | 5 | 9216.89 |
| U | 3 | 8200.25 |
| B2 | 1 | 7834.61 |
| MMLFM | 2 | 5332.58 |
| E | 1 | 2743.98 |
| MLSML | 2 | 2574.16 |
| Insulin | 3 | 2292.18 |
| I4 | 2 | 1215.07 |
| Conodipine | 5 | 1101.04 |
| Prohormone_4 | 2 | 923.05 |
| S | 1 | 782.95 |
| N | 1 | 585.52 |
| Conopressin | 3 | 580.03 |
| Conorfamide | 1 | 537.51 |
| L11 | 1 | 109.06 |
| MASEG | 1 | 96.14 |
| Conikotikot | 2 | 92.8 |
| Conocap | 1 | 90.89 |
| F | 2 | 64.36 |
| O3 | 2 | 24.23 |
| DivMKVAVVLLVS | 2 | 12.63 |
| MGSVP | 1 | 10.09 |

Table S5.

**ToxCodAn-genome full toxin gene annotations**

| Superfamily | # genes |
| --- | --- |
| O2 | 53 |
| O1 | 34 |
| J | 24 |
| M | 17 |
| H | 16 |
| I1 | 7 |
| Conodipine | 7 |
| MSRLF | 6 |
| Conorfamide | 6 |
| Prohormone | 4 |
| A | 4 |
| N | 3 |
| I2 | 3 |
| MWIRK | 2 |
| I4 | 2 |
| Conopressin | 2 |
| S | 1 |
| MMLFM | 1 |
| L11 | 1 |
| Insulin | 1 |
| E | 1 |
| Conocap | 1 |
| B2 | 1 |
| B | 1 |

**Table S6**

**Known doppelgänger toxins and references**

| **Doppelganger toxin** | **Signaling peptide(s)** | **Reference** |
| --- | --- | --- |
| Conopressin | Oxytocin/ Vasopressin (OT/VP) | ^2^ |
| Contulakin-G | Neurotensin (NTS) | ^3^ |
| Cono-NPY | Neuropeptide-F/Y (NPF/Y) | ^4^ |
| ConoCAP | Crustacean cardioactive peptide (CAP) | ^5^ |
| ConoMAP | Myoactive tetradecapeptide (MAP) | ^6^ |
| Con-Insulin | Insulin | ^7^ |
| - | Elevenin (L11) | ^8^ |
| - | Prohormone-4 (PH4) | ^8^ |
| - | Thryostimulin | ^8^ |
| Consomatin | Somatostatin and related peptides (SSRP) | ^9^ |
| Triangle toxin | Triangle doppelgänger related peptide (DREP) | ^10^ |
| CHH toxin | Crustacean Hyperglycemic Hormone (CHH) | ^10^ |
| Tail toxin | Tail DREP | ^10^ |
| Medial toxin | Medial DREP | ^10^ |
| Hairpin toxin | Hairpin DREP | ^10,11^ |

**Table S7. Accession numbers for RNAseq data used in this study**

| **Species** | **Accession number** | **type** |
| --- | --- | --- |
| *Canariensis* | GCA_033310375.1 | WGS |
| *Textile* | GCA_049309315.1 | WGS |
| *Textile* | pending | WGS |
| *Ventricosus* | GCA_018398815.2 | WGS |
| *Abbreviatus* | SRR14407584 | RNAseq |
| *Ammiralis* | SRR14921183 | RNAseq |
| *Arenatus* | SRR2609544 | RNAseq |
| *Aristophanes* | SRR14407587 | RNAseq |
| *Bayani* | SRR13781584 | RNAseq |
| *Betulinus* | SRR2124878 | RNAseq |
| *Boavistensis* | SRR11807497 | RNAseq |
| *Canariensis* | SRR25393579 | RNAseq |
| *Consors* | SRR27489861 | RNAseq |
| *Cordigera* | pending | RNAseq |
| *Coronatus* | SRR2609545 | RNAseq |
| *Cuneolus* | SRR11807496 | RNAseq |
| *Ebraeus* | SRR2609538 | RNAseq |
| *Ermineus* | SRR27489862 | RNAseq |
| *Furvus* | SRR22829302 | RNAseq |
| *Galeao* | SRR11807500 | RNAseq |
| *Generalis* | SRR8821430 | RNAseq |
| *Geographus* | SRR16493595 | RNAseq |
| *Gloriamaris* | SRR5499408 | RNAseq |
| *Grahami* | SRR11807507 | RNAseq |
| *Guanche* | SRR11807502 | RNAseq |
| *Imperialis* | SRR2609542 | RNAseq |
| *Infinitus* | SRR11807493 | RNAseq |
| *Judaeus* | SRR17653514 | RNAseq |
| *Litteratus* | SRR6378469 | RNAseq |
| *Lividus* | SRR2609539 | RNAseq |
| *Magus (adult)* | SRR8195628 | RNAseq |
| *Magus (juvenile)* | SRR23824615 | RNAseq |
| *Marmoreus* | SRR2609532 | RNAseq |
| *Miliaris* | SRR1544627 | RNAseq |
| *Miruchae* | SRR11807495 | RNAseq |
| *Obscurus* | SRR28746436 | RNAseq |
| *Pomacea canaliculata* | GCA_003073045.1 | WGS |
| *Pymorhynchus buccinoides* | GCA_017654935.2 | WGS |
| *Quercinus* | SRR2609537 | RNAseq |
| *Rattus* | SRR2609540 | RNAseq |
| *Raulsilvai* | SRR11807492 | RNAseq |
| *Rolani* | SRR16493597 | RNAseq |
| *Sponsalis* | SRR2609541 | RNAseq |
| *Striatus* | SRR8195630 | RNAseq |
| *Terebra* | SRR8195627 | RNAseq |
| *Textile* | SRR8195629 | RNAseq |
| *Textile FT* | pending | RNAseq |
| *Textile HP* | pending | RNAseq |
| *Textile NR* | pending | RNAseq |
| *Textile SG* | pending | RNAseq |
| *Textile VB* | pending | RNAseq |
| *Textile VG* | pending | RNAseq |
| *Tribblei* | SRR1803937 | RNAseq |
| *Trochulus* | SRR11807506 | RNAseq |
| *Varius* | SRR2609543 | RNAseq |
| *Ventricosus* | SRR13740844 | RNAseq |
| *Verdensis* | SRR11807498 | RNAseq |
| *Virgo* | SRR2608262 | RNAseq |

Table S8. Parameters and PacBio sequencing yields for each of the three HiFi libraries.

|  | Sheared DNA pre-treatment | Average size | BluePippin size-selection cutoff | HiFi yield | HiFi reads | Polymerase read length |
| --- | --- | --- | --- | --- | --- | --- |
| Library 1 | NA | 19 kbp | 10 kb | 24.7 Gb | 1.83 M | 39.7 kb |
| Library 2 | powerClean | 26 kbp | 10 kb | 15.6 Gb | 1.05 M | 26.3 kb |
| Library 3 | powerClean | 13 kbp | 6 kb | 38.4 Gb | 4.54 M | 42.6 kb |

Movie S1.

*Conus textile* hunting on a coral reef at night in the Philippines.

Movie S2.

*Conus textile* hunting and feeding on a turbinidae.

Movie S3.

*Conus textile* hunting *Conus eburneus.*

Movie S4.

*Conus textile* hunting *Conus floridulus*.

Data S1. (separate file)

Annotated *Conus* neuropeptides

Data S2. (separate file)

New calcitonin doppelgänger toxins

Data S3. (separate file)

New egg-laying hormone doppelgänger toxins

Data S4. (separate file)

New FxRIamide doppelgänger toxins

Data S5. (separate file)

New GGNamide doppelgänger toxins

Data S6. (separate file)

New HFAamide doppelgänger toxins

Data S7. (separate file)

New NdWFamide doppelgänger toxins

Data S8. (separate file)

New neuropeptide KY doppelgänger toxins

Data S9. (separate file)

Genomic loci for MMLFM, conoNPY, and NPF/Y genes in *C. textile*
